# The Host Cell’s Endoplasmic Reticulum Proteostasis Network Profoundly Shapes the Protein Sequence Space Accessible to HIV Envelope

**DOI:** 10.1101/2021.04.24.441266

**Authors:** Jimin Yoon, Emmanuel E. Nekongo, Jessica E. Patrick, Angela M. Phillips, Anna I. Ponomarenko, Samuel J. Hendel, Vincent L. Butty, C. Brandon Ogbunugafor, Yu-Shan Lin, Matthew D. Shoulders

**Author notes:** These authors contributed equally to this work (Jimin Yoon, Emmanuel E. Nekongo). To whom correspondence may be addressed: Matthew D. Shoulders, Department of Chemistry, Massachusetts Institute of Technology, 77 Massachusetts Avenue, 16-573A, Cambridge, MA 02139.

## Abstract

The sequence space accessible to evolving proteins can be enhanced by cellular chaperones that assist biophysically defective clients in navigating complex folding landscapes. It is also possible, however, for proteostasis mechanisms that promote strict quality control to greatly constrain accessible protein sequence space. Unfortunately, most efforts to understand how proteostasis mechanisms influence evolution rely on artificial inhibition or genetic knockdown of specific chaperones. The few experiments that perturb quality control pathways also generally modulate the levels of only individual quality control factors. Here, we use chemical genetic strategies to tune proteostasis networks via natural stress response pathways that regulate levels of entire suites of chaperones and quality control mechanisms. Specifically, we upregulate the unfolded protein response (UPR) to test the hypothesis that the host endoplasmic reticulum (ER) proteostasis network shapes the sequence space accessible to human immunodeficiency virus-1 (HIV) envelope (Env) protein. Elucidating factors that enhance or constrain Env sequence space is critical because Env evolves extremely rapidly, yielding HIV strains with antibody and drug escape mutations. We find that UPR-mediated upregulation of ER proteostasis factors, particularly those controlled by the IRE1-XBP1s UPR arm, globally reduces Env mutational tolerance. Conserved, functionally important Env regions exhibit the largest decreases in mutational tolerance upon XBP1s activation. This phenomenon likely reflects strict quality control endowed by XBP1s-mediated remodeling of the ER proteostasis environment. Intriguingly and in contrast, specific regions of Env, including regions targeted by broadly neutralizing antibodies, display enhanced mutational tolerance when XBP1s is activated, hinting at a role for host proteostasis network hijacking in potentiating antibody escape. These observations reveal a key function for proteostasis networks in decreasing instead of expanding the sequence space accessible to client proteins, while also demonstrating that the host ER proteostasis network profoundly shapes the mutational tolerance of Env in ways that could have important consequences for HIV adaptation.

## Introduction

Protein mutational tolerance is constrained by the biophysical properties of the evolving protein. Selection to maintain proper protein folding and structure purges a large number of otherwise possible mutations that would be functionally beneficial [1–5]. It is no surprise, then, that cellular proteostasis networks play a key role in defining the protein sequence space accessible to client proteins. [6–17]. For example, much attention has been given to the phenomenon of chaperones increasing sequence space accessible to their client proteins, likely by promoting the folding of protein variants with biophysically deleterious amino acid substitutions [7–11]. Most efforts in this area have focused specifically on how the activities of the heat shock proteins Hsp90 and Hsp70 can expand protein sequence space, in part owing to the availability of specific inhibitors that enable straightforward comparative studies of protein evolution in the presence versus the absence of folding assistance.

In contrast to chaperones increasing sequence space, one might anticipate that protein folding quality control factors would constrain the sequence space accessible to evolving client proteins. For example, promoting the rapid degradation and removal of slow-folding or aberrantly folded protein variants could cut off otherwise accessible evolutionary trajectories [16–18], especially if those variants might have still maintained some level of function if instead allowed to persist in the cellular environment. Unfortunately, efforts to understand potential contributions of quality control to shaping protein sequence space are limited. This gap in understanding is particularly problematic because natural cellular mechanisms to remodel proteostasis networks function via stress-responsive transcription factors [19, 20], not via inhibition or repression of individual chaperones. These stress responses tune the levels both chaperones and quality control mechanism simultaneously. Our understanding of how the potentially competing phenomena of upregulated chaperones expanding sequence space versus upregulated quality control restricting sequence space manifest in the context of various evolving client proteins remains shallow.

Here, we sought to evaluate whether and how the unfolded protein response (UPR)-regulated endoplasmic reticulum (ER) proteostasis network influences the sequence space accessible to membrane proteins processed by the secretory pathway. In particular, we aimed to use chemical genetic control of the UPR to broadly modulate the composition of the ER proteostasis network, and then use deep mutational scanning (DMS) to assess how such perturbations alter the accessible protein sequence space. We chose human immunodeficiency virus-1 (HIV) envelope (Env), a trimeric surface glycoprotein that is folded and quality-controlled by the ER, as our model protein. We selected Env because its rapid evolution during HIV infections plays a critical role in HIV developing drug and host cell antibody resistance [21–23]. Additionally, Env interacts extensively with various components of the ER proteostasis network, including the ER chaperones calnexin [24] and calreticulin [25], binding immunoglobulin protein (BiP) [26], and ER alpha mannosidase to initiate ER-associated degradation (ERAD) [27, 28], suggesting the strong potential for the host’s ER proteostasis network to shape Env’s accessible sequence space.

Importantly, recent work has revealed that the cellular proteostasis network can indeed impact the sequence space of not just endogenous client proteins, but also viral proteins that hijack their host’s proteostasis machinery [29–33]. This relationship has critical evolutionary and therapeutic implications, because mutational tolerance is directly associated with the ability of a virus to evade the host’s innate and adaptive immune responses, as well as antiviral drugs [34–40]. Early work in this area focused on how viruses like influenza and poliovirus hijack the host’s heat shock response-regulated cytosolic chaperones to enhance their mutational tolerance [29–31]. More recently, we discovered that host UPR-mediated upregulation of the ER proteostasis network increases the mutational tolerance of influenza A hemagglutinin specifically at febrile temperatures [32]. Aside from the hemagglutinin work, no comprehensive studies testing the influence of the ER proteostasis network on client protein evolution, whether viral or endogenous, are available.

In this study, we used chemical genetic tools to specifically upregulate the inositol requiring enzyme-1 / X-box binding protein-1 spliced (IRE1-XBP1s) and the activating transcription factor 6 (ATF6) transcriptional arms of the UPR separately or in tandem [41]. This approach provided user-defined modulation of the composition of the host’s ER proteostasis network. We hypothesized that the resulting distinct host ER proteostasis environments would critically influence the protein sequence space accessible to Env. We tested this hypothesis using DMS and observed a global decrease in Env mutational tolerance upon UPR upregulation. The effect proved particularly strong upon XBP1s-mediated enhancement of the ER proteostasis environment. In addition, we observed that sites with different structural or functional roles respond differently to UPR upregulation. For example, despite the general reduction in mutational tolerance across the Env sequence, which was especially strong for conserved regions, a number of sites still exhibited an increase in mutational tolerance. These sites displaying increased mutational tolerance included a number of sites targeted by broadly-neutralizing antibodies.

Altogether, this work provides experimental evidence that the composition of the ER proteostasis network profoundly shapes the protein sequence space accessible to membrane proteins. Furthermore, it demonstrates for the first time that the combined upregulation of chaperones and quality control factors can actually greatly decrease mutational tolerance of the client protein. Critically, the details of the interaction between the host cell proteostasis network and viral proteins differ from one protein to another, and can vary even within different regions of the same protein.

## Results

### Chemical genetic control of ER proteostasis network composition during HIV infection

We began by generating a cell line in which HIV could robustly replicate and we could chemically activate the UPR’s IRE1-XBP1s and ATF6 transcriptional responses separately or simultaneously, in an ER stress-independent manner. We selected the IRE1-XBP1s and ATF6 arms of the UPR for chemical control because they (rather than the PERK arm of the UPR) are largely responsible for defining levels of ER chaperones and quality control factors [20, 41, 42] that would likely influence Env folding, degradation, and secretion. We sought ER stress-independent activation of these transcription factors rather than stress-mediated, global UPR induction, owing to the pleiotropic effects of chemical stressors and the non-physiologic, highly deleterious consequences of inducing high levels of protein misfolding in the secretory pathway [19, 32, 41, 43].

To allow for robust replication of HIV, we chose human T cell lymphoblasts (SupT1 cells). SupT1 cells support high levels of HIV replication in cell culture, likely due to the lack of cytidine deaminase activity that can cause hypermutation of HIV DNA [44]. To attain control of the IRE1-XBP1s pathway in this HIV replication-competent cell line, we stably introduced the gene encoding the XBP1s transcription factor under control of the tetracycline repressor into SupT1 cells (**Fig 1A**). In the resulting cells, the IRE1-XBP1s transcriptional response can be induced simply by treatment with the tetracycline repressor-binding small molecule doxycycline (dox). We then used a second, orthogonal chemical genetic strategy to regulate the ATF6 transcriptional response using another small molecule. Specifically, we transduced the XBP1s-inducible SupT1 cells with a gene encoding an *Escherichia coli* dihydrofolate reductase (DHFR)-based destabilizing domain fused to the active form of ATF6 (amino acid residues 1–373 of ATF6; **Fig 1A**). The DHFR.ATF6 fusion protein is constitutively expressed but rapidly degraded by the proteasome, owing to the largely unfolded nature of the destabilizing domain version of DHFR [45], preventing any ATF6 transcriptional activity. However, treatment with the small molecule trimethoprim (TMP) stabilizes the folded state of the DHFR destabilizing domain and thereby induces ATF6 transcriptional activity [46]. We termed these cells SupT1^DAX^ cells (**Fig 1A**), with the DAX signifier indicating the inclusion of both the <u>D</u>HFR.<u>A</u>TF6 and <u>X</u>BP1s constructs.

**Fig 1.**
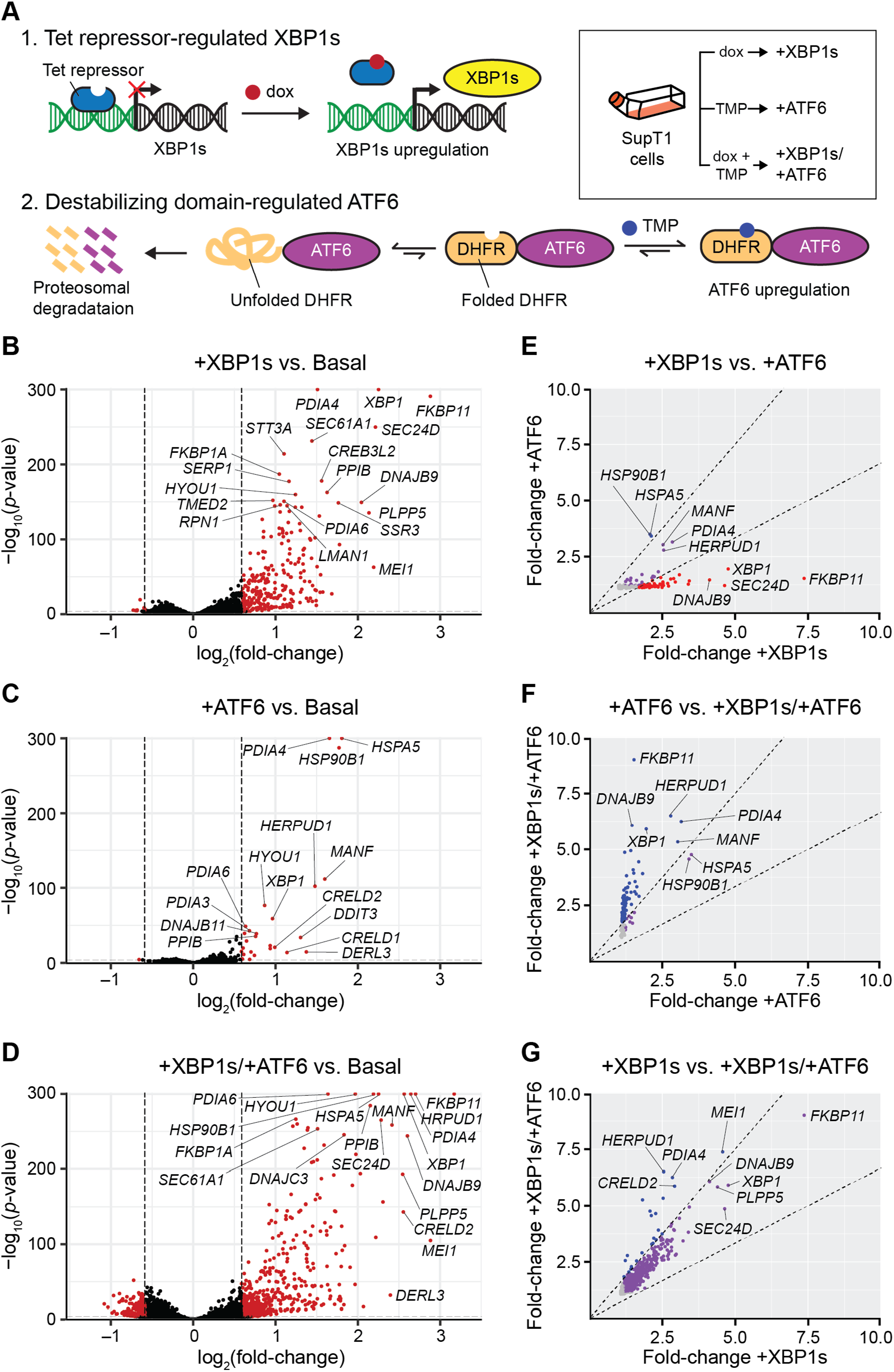
Stress-independent activation of XBP1s, ATF6, or XBP1s and ATF6 creates four distinct ER prote-ostasis environments in SupT1^DAX^ cells (basal, +XBP1s, +ATF6, +XBP1s/+ATF6). (**A**) Chemical genetic strategy to orthogonally regulate XBP1s and ATF6 in SupT1^DAX^ cells. (**B–D**) RNA-Seq analysis of the transcriptomic consequences of (**B**) XBP1s, (**C**) ATF6, and (**D**) XBP1s/ATF6 induction. Transcripts that were differentially expressed under each condition based on a >1.5-fold change in expression level (for dox-, TMP-, or dox and TMP-treated versus vehicle-treated cells) and a non-adjusted *p*-value <10^−10^ are plotted in red, with select transcripts labeled. The lowest nonzero *p*-value recorded was 10^−291^; therefore, *p*-values = 0 were replaced with *p*-values = 1.00 × 10^−300^ for plotting purposes. Transcripts for which *p*-values could not be calculated owing to extremely low expression or noisy count distributions were excluded from plotting. (**E–G**) Comparison of transcript fold-change upon (**E**) +XBP1s versus +ATF6 (**F**) +ATF6 versus +XBP1s/+ATF6, and (**G**) +XBP1s versus +XBP1s/+ATF6 remodeling of the ER proteostasis network. Only transcripts with false discovery rate-adjusted *p*-value <0.05 and fold-increase >1 in both of the indicated conditions are plotted. Dashed lines indicate a 1.5-fold filter to assign genes as selectively induced by the proteostasis condition on the *x*-axis (red), *p*-axis (blue), or lacking selectivity (purple). Transcripts with fold-increase <1.2 in either proteostasis environment are colored in grey to indicate low differential expression.

With stably engineered SupT1^DAX^ cells in hand, we anticipated that activation of the XBP1s and ATF6 transcriptional responses, either separately or together, would create four distinct ER proteostasis environments (basal, XBP1s-activated, ATF6-activated, and XBP1s/ATF6 both activated) that could then be used to assess potential consequences for Env mutational tolerance. We treated SupT1^DAX^ cells with DMSO (vehicle), dox (to activate the XBP1s transcriptional response), TMP (to activate the ATF6 transcriptional response), and both dox and TMP (to simultaneously activate the XBP1s and ATF6 transcriptional responses) for 18 h and used RNA-Seq to evaluate resultant changes in the transcriptome (see **S1 Table** for complete results). We applied gene set enrichment analysis [47] to the RNA-Seq results using the MSigDB c5 collection, and found that gene sets related to ER stress, Golgi trafficking, and ERAD were highly enriched upon induction of XBP1s and/or ATF6 (**S2 Table**). In contrast, gene sets that serve as markers of other stress responses (e.g., the heat shock response) were not enriched, consistent with a highly selective, stress-independent induction of UPR transcriptional responses.

To identify transcripts that were differentially expressed in the modified ER proteostasis environments, we compared the transcriptome of SupT1^DAX^ cells treated with dox, TMP, or both dox and TMP to the basal transcriptome (**Figs 1B–D** and **S1 Table**). We observed upregulation of 223 transcripts upon XBP1s induction (+XBP1s), 24 transcripts upon ATF6 induction (+ATF6), and 436 transcripts upon simultaneous induction of XBP1s and ATF6 (+XBP1s/+ATF6); here, upregulation was defined as a change in expression level >1.5-fold relative to the basal environment with a non-adjusted *p*-value <10^-10^. For all three treatment conditions, the up-regulated transcripts were strongly biased towards UPR-regulated components of the ER proteostasis network. Furthermore, transcripts known to be targeted primarily by XBP1s were strongly upregulated upon dox treatment (e.g., *SEC24D* and *DNAJB9*), whereas transcripts known to be targeted primarily by ATF6 were more strongly upregulated upon TMP treatment (e.g., *HSP90B1*) [41, 48, 49]. Moreover, genes known to be targets of XBP1s and ATF6 heterodimers, such as *HERPUD1* [41, 50], were upregulated to the highest extent only when both XBP1s and ATF6 were induced in tandem.

To analyze the extent to which these three perturbations (+XBP1s, +ATF6, and +XBP1s/+ATF6) engendered unique ER proteostasis environments, we cross-compared the mRNA fold-changes owing to each treatment. As expected, XBP1s or ATF6 activation resulted in the upregulation of partially overlapping gene sets. XBP1s activation notably caused an extensive remodeling of the entire ER proteostasis network, whereas ATF6 activation resulted in targeted upregulation of just a select subset of ER chaperones (**Fig 1E**). The combined activation of XBP1s and ATF6 provided access to a third environment where specific transcripts were more strongly upregulated than upon the single activation of either transcription factor alone (**Figs 1F** and **1G**). These results are consistent with prior work showing that ATF6 induction causes upregulation of fewer transcripts than XBP1s [41, 49], as well as with the ability of XBP1s and ATF6 to synergistically upregulate a larger set of transcripts than either transcription factor alone, likely via heterodimerization of the transcription factors [41, 50, 51]. Taken together, our RNA-Seq results show that we can access four distinctive ER proteostasis environments for Env mutational tolerance experiments via chemical genetical control of XBP1s and ATF6 (basal, +XBP1s, +ATF6, and +XBP1s/+ATF6).

Finally, we assessed whether these perturbations of the ER proteostasis environment had deleterious effects on cell viability or restricted HIV replication, as we had previously observed inhibition of HIV replication upon upregulation of the heat shock response [52]. To address the former, we activated XBP1s and/or ATF6 in SupT1^DAX^ cells and measured resazurin metabolism 72 h post-drug treatment (**S1 Fig A**). We observed that induction of XBP1s and ATF6, either separately or simultaneously, did not alter the metabolic activity of SupT1^DAX^ cells, consistent with no deleterious effects on cell viability. To address the latter, we used the TZM-bl assay to quantify HIV infectious titer (**S1 Fig B**). Specifically, we used TZM-bl reporter cells containing the *E. coli* β-galactosidase gene under the control of an HIV long-terminal repeat sequence [53]. When these cells are infected with HIV, the HIV Tat transactivation protein induces expression of β-galactosidase, which cleaves the chromogenic substrate (X-Gal) and causes infected cells to appear blue in color. The infectious titer increased marginally by approximately 3.5-fold when XBP1s was induced, either alone or together with ATF6. Induction of ATF6 alone did not affect HIV infectious titer. Thus, ER proteostasis network perturbation via XBP1s and/or ATF6 activation did not deleteriously impact HIV replication.

### Deep mutational scanning of Env in four distinctive host ER proteostasis environments

We next sought to apply DMS to Env to test our hypothesis that the composition of the host’s ER proteostasis network plays a central role in determining the mutational tolerance of Env. For this purpose, we employed a previously developed set of three replicate Env proviral plasmid libraries [22], created by introducing random codon mutations at amino acid residues 31–702 of the Env protein (note that the HXB2 numbering scheme [54] is used throughout). In these libraries, the N-terminal signal peptide and the C-terminal cytoplasmic tail were excluded from mutagenesis owing to their dramatic impact on Env expression and/or HIV infectivity [22].

We generated biological triplicate viral libraries from these mutant Env plasmid libraries by transfecting the plasmid libraries into HEK293T cells and then harvesting the passage 0 (p0) viral supernatant after 4 d. Deep sequencing of the three p0 viral libraries showed that 84% of all possible amino-acid mutations were observed at least three times in at least one of the triplicate libraries, consistent with prior work [22, 36]. Next, to establish a genotype–phenotype link we passaged the p0 transfection supernatants in SupT1 cells at a very low multiplicity of infection (MOI) of 0.005 infectious virions/cell. We next performed batch competitions of each individual Env viral library in SupT1^DAX^ cells in each of the four different ER proteostasis environments: basal, +XBP1s, +ATF6, and +XBP1s/+ATF6 (**Fig 2A**). Briefly, SupT1^DAX^ cells were treated for 18 h with vehicle, dox, TMP, or both dox and TMP to generate the intended ER proteostasis environment, followed by infection with p1 viral supernatant at a MOI of 0.005 infectious virions/cell. We used this MOI to minimize co-infection of individual cells and thereby maintain the genotype–phenotype link. 96 h post-infection, we extracted non-integrated viral DNA, and then generated PCR amplicons of *Env* [22]. Finally, we deep-sequenced the amplicons using barcoded-subamplicon sequencing and analyzed the sequencing reads using the dms_tools2 suite (https://jbloomlab.github.io/dms_tools2/) [55, 56].

**Fig 2.**
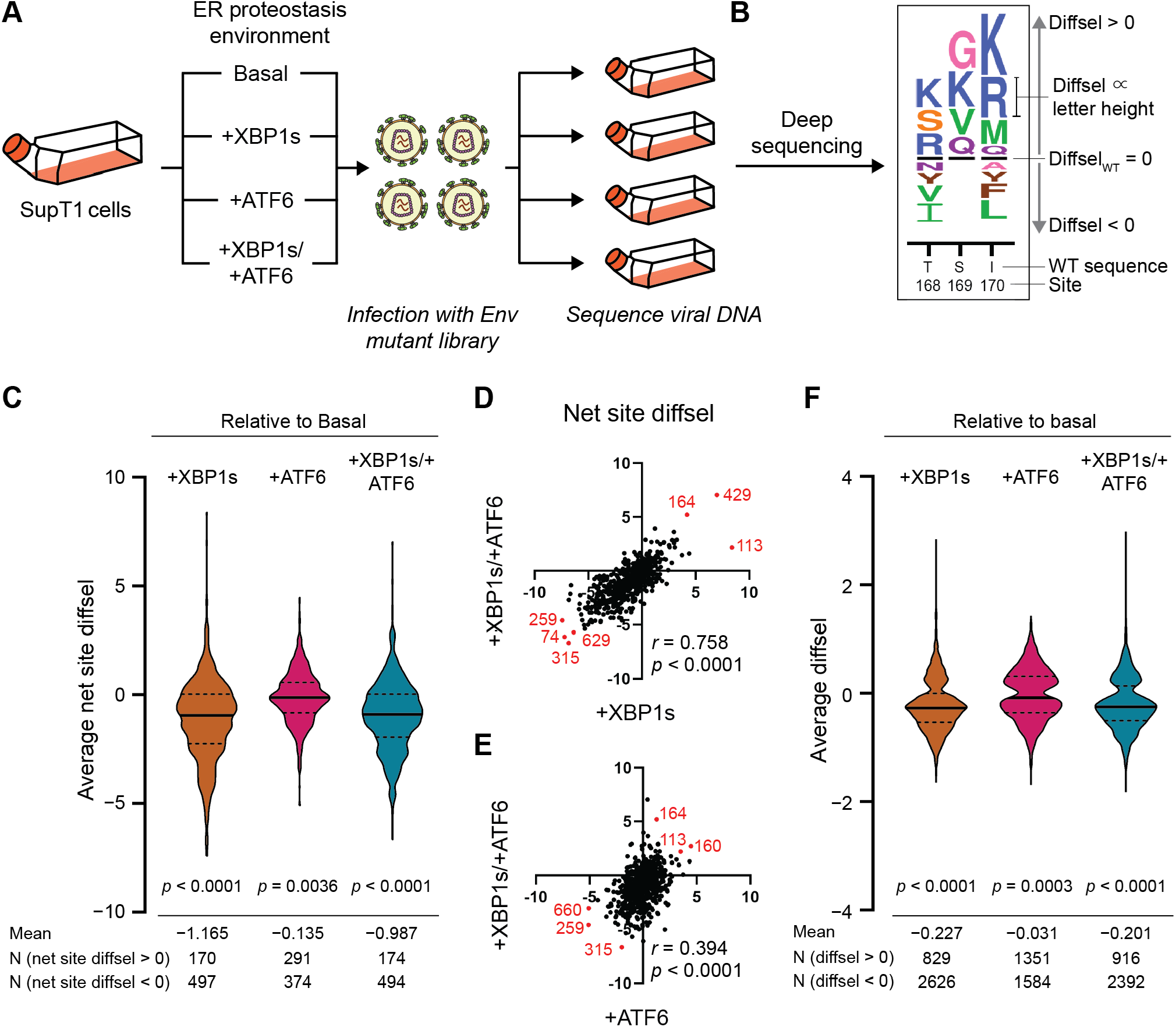
Upregulation of the host cell’s ER proteostasis environment generally reduces mutational tolerance across the Env protein sequence. **(A)** Scheme for deep mutational scanning of Env in four distinctive ER proteostasis environments (basal, +XBP1s, +ATF6, +XBP1s/+ATF6). SupT1^DAX^ cells were pre-treated with DMSO (Basal), dox (+XBP1s), TMP (+ATF6), or both dox and TMP (+XBP1s/+ATF6) 18 h prior to infection with biological triplicate Env libraries. 4 d post-infection, cells were harvested and non-integrated viral DNA was sequenced to quantify diffsel of Env variants. **(B)** Diffsel for each amino acid variants can be visualized in a sequence logo plot. The black horizontal lines represent the diffsel for the wild-type amino acid at that site, and the height of the amino acid letter abbreviations are proportional to the diffsel of each variant in the remodeled ER proteostasis environment relative to the basal environment. Variants that are relatively enriched in the indicated ER proteostasis environment (positive diffsel) are located above the black horizontal line. Variants that are relatively depleted in the indicated ER proteostasis environment (negative diffsel) are located below the black horizontal line. (**C**) Net site diffsel for all Env sites in three perturbed ER proteostasis environments, averaged over biological triplicates. The black horizontal lines on the violin plots indicate the median (solid line) or the first and the third quartiles (dashed lines) of the distribution. The significance of deviation from null (net site diffsel = 0, no selection) was tested using a one-sample *t*-test, with two-tailed *p*-values shown. The mean of distribution, as well as the number of sites with net site diffsel >0 or <0, are listed below the distribution. (**D–E**) Correlation for net site diffsel values between (**D**) +XBP1s/+ATF6 versus +XBP1s and (**E**) +XBP1s/+ATF6 versus +ATF6, normalized to the basal proteostasis environment. Pearson correlation coefficients *r* and corresponding *p*-values are shown. Select sites with highly positive or highly negative net site diffsel values in both proteostasis environments are marked in red and labeled. (**F**) Diffsel for individual Env variants in three perturbed ER proteostasis environments, averaged over biological triplicates. The black horizontal lines on the violin plots indicate the median (solid line) or the first and the third quartiles (dashed lines) of the distribution. The significance of deviation from null (diffsel = 0, no selection) was tested using a one-sample *t*-test, with twotailed *p*-values shown. The mean of distribution, as well as the number of sites with diffsel >0 and <0, are listed below the distribution.

To identify amino acid variants that were differentially enriched or depleted in a given ER proteostasis selection condition (+XBP1s, +ATF6, or +XBP1s/+ATF6) relative to the basal ER proteostasis environment, we quantified differential selection (diffsel). Diffsel was calculated by taking the logarithm of the variant’s enrichment in the selection condition relative to its enrichment in the basal ER proteostasis network condition (**Fig 2B**). In addition, to decipher reliable signal from experimental noise, we filtered the DMS data using a previously described and validated two-step strategy [32]. First, we removed variants that were not present in all three pre-selection replicate viral libraries. That is, we eliminated even those variants that were strongly enriched or depleted in two replicates if they were not present in the starting library of the third replicate. Second, we removed variants that exhibited diffsel in opposite directions in any of the biological triplicates. Using the second filter, we typically removed variants that were minimally affected by the selection, displaying slightly positive diffsel values in one replicate but slightly negative diffsel values in another. By applying these two filters, we were able to focus subsequent analyses only on Env variants that exhibited robust, reproducible diffsel across biological triplicates of the same ER proteostasis network conditions (3,455 variants for +XBP1s, 2,935 variants for +ATF6, and 3,308 variants for +XBP1s/+ATF6).

### XBP1s-mediated ER proteostasis network remodeling causes a strong net decrease in the mutational tolerance of Env, whereas ATF6 has minimal effects

To evaluate our hypothesis that the composition of the host’s ER proteostasis network critically shapes Env mutational tolerance, we first analyzed the ‘net site diffsel’ in each host ER proteostasis environment. Net site diffsel is the sum of individual mutational diffsel values for a given Env site. Thus, a positive net site diffsel indicates that mutational tolerance at a given Env site is quantitatively increased in an enhanced host ER proteostasis environment relative to the basal ER proteostasis environment. In contrast, a negative net site diffsel indicates that mutational tolerance is decreased in an enhanced host ER proteostasis environment. For example, the net site diffsel for site 169 (**Fig 2B**) would be the sum of the diffsel values for G, K, V, and Q, which would be positive and therefore we would conclude that mutational tolerance increased at site 169.

Using the filtered Env DMS data sets, we calculated net site diffsel at each Env position averaged across the three biological replicates of our experiment (**Fig 2C**). Strikingly, the +XBP1s ER proteostasis environment globally, substantially, and significantly reduced mutational tolerance across the entire Env protein (mean net site diffsel = −1.165, *p*-value < 0.0001). Combined activation of both XBP1s and ATF6 had a similar effect, again substantially and significantly reducing Env mutational tolerance (mean net site diffsel = −0.987, *p*-value < 0.0001). The magnitude of mean net site diffsel was approximately 15-fold larger upon XBP1s activation than we previously observed for increased mutational tolerance in influenza hemagglutinin in an XBP1s-activated ER proteostasis environment at 37 °C [32]. Thus, Env mutational tolerance is exceptionally sensitive to XBP1s-mediated ER proteostasis network upregulation, to a much greater extent than hemagglutinin. In contrast, the +ATF6 ER proteostasis environment, while still mildly reducing mutational tolerance across Env, had a less substantial global effect (mean net site diffsel = −0.135, *p*-value = 0.0036). The latter result suggests that the reduced Env mutational tolerance observed in the +XBP1s/+ATF6 ER proteostasis environment was largely driven by ER proteostasis factors targeted by XBP1s. Indeed, the Pearson correlation coefficient *r* was substantially higher between the net site diffsel values observed in the +XBP1s versus +XBP1s/+ATF6 environments (*r* = 0.758; **Fig 2D**) than between those observed in the +ATF6 versus +XBP1s/+ATF6 (*r* = 0.394; **Fig 2E**). This observation aligns well with our RNA-Seq data, in which we observed substantially more overlap between the ER proteostasis network transcriptome remodeling caused by XBP1s activation versus the combination of XBP1s and ATF6 activation, compared to ATF6 activation versus the combination of XBP1s and ATF6 activation (**Fig 1F** and **G**).

It is important to note that in a net site diffsel analysis we quantify the relative enrichment of all amino acid variants combined to assess mutational tolerance at a given Env site. Consequently, a decrease in mutational tolerance as measured by net site diffsel could be caused by a single amino acid variant that was strongly disfavored or, alternatively, by many variants being disfavored relative to wild-type. Therefore, to test if individual amino acid variants also reveal a global trend towards reduced mutational fitness, we plotted the individual diffsel values for all Env variants. We again observed reduced mutational fitness of the majority of Env variants whenever XBP1s was induced, indicating that the effect is largely driven by a general loss of mutational tolerance rather than by large effect sizes for just a few specific amino acid variants (**Fig 2F**).

In sum, there is a striking decrease in mutational tolerance across much of Env upon XBP1s-mediated remodeling of the host’s ER proteostasis network. This broad and substantive trend should not, however, mask the fact that many sites displayed strongly enhanced mutational tolerance upon not just XBP1s activation but also ATF6 activation (e.g., S164, D113) (**Fig 2C**; N net site diffsel > 0). Finally, it should be noted that although ATF6 activation had minimal global consequences for Env mutational tolerance, there were still a number of sites where reduced net site diffsel (e.g., L259, R315) was observed across all three enhanced ER proteostasis environments (**Fig 2D** and **2E**).

### Investigation of Env sites and variants most strongly impacted by the host’s ER proteostasis network

To visualize the relative fitness of individual amino acid variants in each host ER proteostasis environment, we generated sequence logo plots across the entire Env sequence (**Fig 3A**, **S2 Fig**, and **S3 Fig**). The relative enrichment for each amino acid variant (diffsel) was calculated from our filtered data sets by averaging across three biological replicates. The unfiltered, unaveraged full sequence logo plots for each replicate and condition are also provided in **S2 File**.

**Fig 3.**
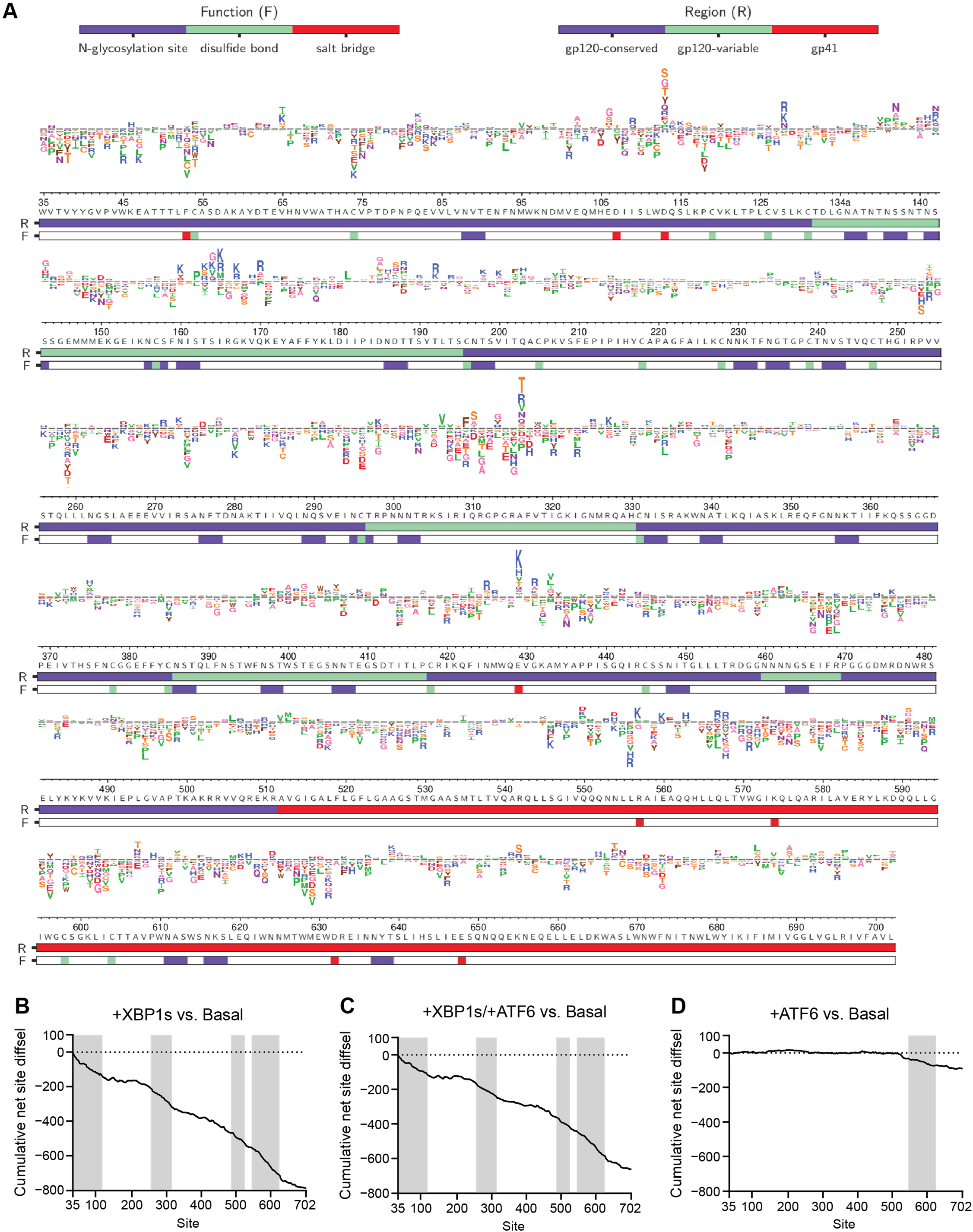
Differential selection (diffsel) across Env upon remodeling of the host’s ER proteostasis network. (**A**) Logo plot displaying averaged diffsel for +XBP1s normalized to the basal proteostasis environment. The height of the amino-acid abbreviation corresponds to the magnitude of diffsel. The amino-acid abbreviations are colored based on the side-chain properties: negatively charged (D, E; red), positively charged (H, K R; blue), polar uncharged (C, S, T; yellow), small nonpolar (A, G; pink), aliphatic (I, L, M, P, V; green), and aromatic (F, W, Y; brown). The numbers and letters below the logos indicate the Env site in HXB2 numbering and the identity of the wild-type amino acid for that site, respectively. The color bar below the logos indicates the function (F) that the site is involved in (*N*-glycosylation site (blue), disulfide bond (green), or salt bridge (red)) or the region (R) of Env that the site belongs to (gp120-conserved (blue), gp120-variable (green), or gp41 (red); the sites that belong to the five variable loops of gp120 were categorized as ‘gp120-variable’, and the sites that are not included in the five variable loops were categorized as ‘gp120-conserved’). Only variants that were present in all three pre-selection viral libraries and exhibited diffsel in the same direction across all three biological triplicates are plotted here. Unfiltered logo plots for each individual replicate are provided in **S2 File**. (**B–D**) Cumulative net site diffsel across Env sites for (**B**) +XBP1s, (**C**) +XBP1s/+ATF6, and (**D**) +ATF6, normalized to the basal proteostasis environment. Regions where the decrease in mutational tolerance is particularly prominent are shaded in grey (35–120, 260–320, 490–530, and 550–630 for (**B**) and (**C**), 550–630 for (**D**)).

Several features of these logo plots are immediately noteworthy. First, the global and relatively similar reduction in mutational tolerance caused by XBP1s activation (**Fig 3A**) and combined XBP1s and ATF6 activation (**S2 Fig**) is readily observed. To visualize this phenomenon and highlight specific regions in which the effect size is particularly large, we plotted cumulative net site diffsel against Env sites (**Fig 3B–D**). We observed that the decrease in mutational tolerance is most prominent around sites 35–120, 260–320, 490–530, and 550– 630 when XBP1s was activated alone (**Fig 3A** and **3B**) or together with ATF6 (**S2 Fig** and **Fig 3C**), as indicated by the steeper slopes in those regions. For ATF6 activation alone (**S3 Fig** and **Fig 3D**), the reduction in mutational tolerance was focused close to the C-terminus around sites 550–660. We assessed whether this differential impact of ER proteostasis mechanisms is based on surface accessibility of sites, but did not observe any correlation between net site diffsel and surface accessibility across Env sites for either the Env monomer or the trimer (**S4 Fig** and **S3 Table**). Second, although the general trend towards reduced mutational tolerance is quite striking, it is also apparent that there are specific positions where either XBP1s- or ATF6-mediated ER proteostasis network enhancement strongly enhanced mutational tolerance at a given site (e.g., D113) or enhanced the fitness of a specific mutation (e.g., I309F). Third, the stronger impacts of XBP1s activation compared to ATF6 activation are apparent (**Fig 3A** versus **S3 Fig** and **Fig 3B** versus **3D**), as is the overall similarity of the impacts of XBP1s activation to the simultaneous activation of both XBP1s and ATF6 (**Fig 3A** versus **S2 Fig** and **Fig 3B** versus **3C**).

To assess whether or not the global decrease in mutational tolerance could be attributed to specific structural or functional regions, we calculated average net site diffsel for individual functional/structural groups. These groups include (1) the entire gp120 and gp41 subunits, (2) the conserved and variable regions of gp120, where the conserved region is defined as the region that does not belong to the five variable loops of gp120, (3) the five variable loops of gp120 individually (denoted V1–5), (4) regions responsible for viral membrane fusion, and (5) other sites with important functional and structural roles (**Fig 4**, **S5 Fig**, and **S6 Fig**; see corresponding references for assignment of these regions in **S4 Table**).

**Fig 4.**
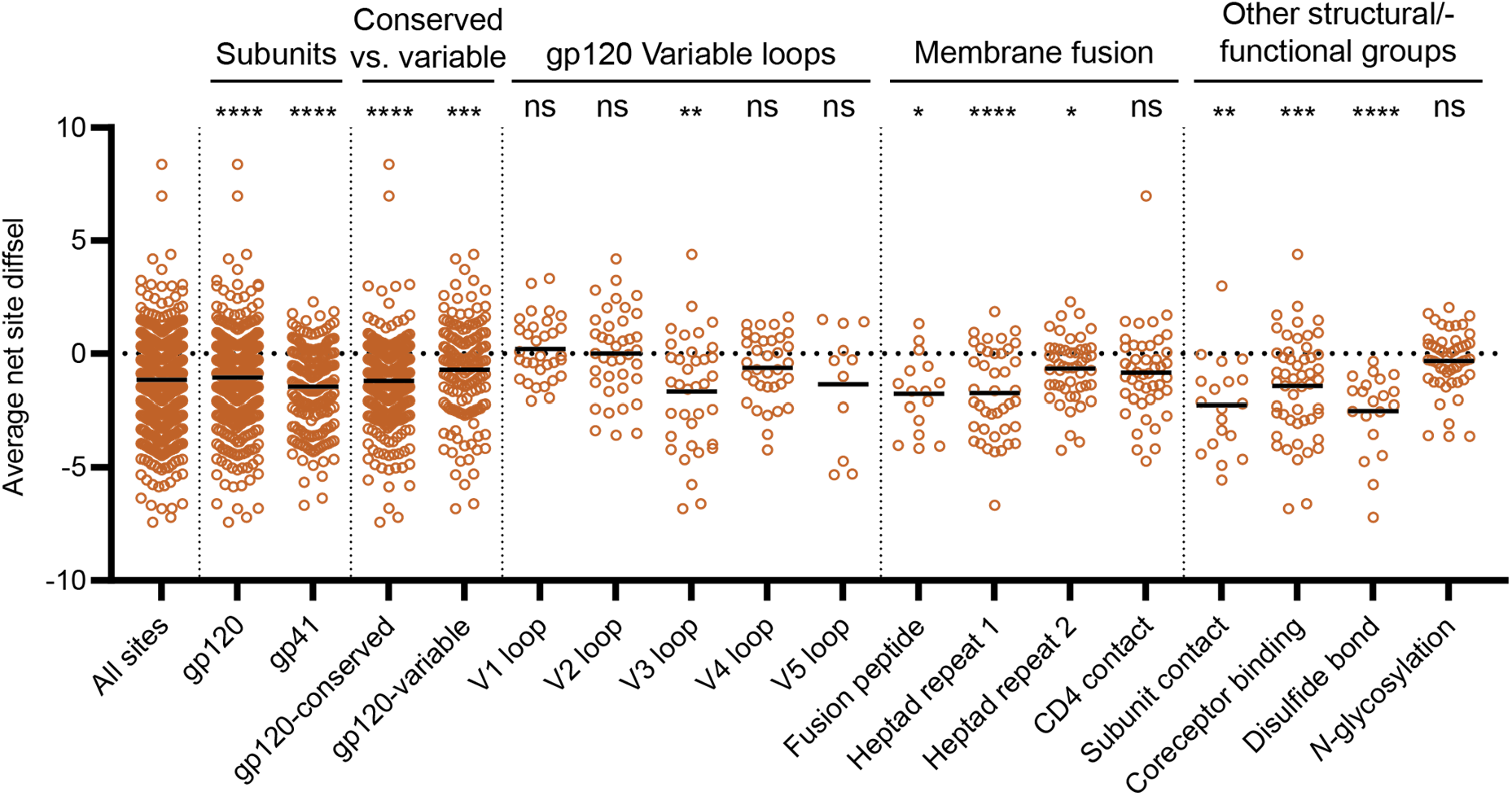
Impact of XBP1s induction on mutational tolerance varies across Env structural elements. Average net site diffsel for the +XBP1s ER proteostasis environment normalized to the basal ER proteostasis environment, where medians of distributions are indicated by black horizontal lines. Sites are sorted by subunits, conserved vs. variable regions of gp120, five variable loops of gp120, regions important for membrane fusion, and other structural/functional groups. For ‘Conserved vs. variable’, the sites that belong to the five variable loops of gp120 were categorized as ‘gp120-variable’, and the sites that are not included in the five variable loops were categorized as ‘gp120-conserved’. Significance of deviation from null (net site diffsel = 0, no selection) was tested using a one-sample *t*-test. The derived *p*-values were Bonferroni-corrected for 18 tests and *, **, ***, and **** represent adjusted two-tailed *p*-values of <0.05, <0,01, <0.001, and <0.0001.

We focused first on the consequences of XBP1s induction, because the effects were larger than for ATF6 induction and similar to the consequences of dual induction (**Fig 4**). We observed a decrease in mutational tolerance for both gp120 and gp41, indicating that XBP1s upregulation impacts both subunits of Env, albeit gp41 more strongly (**Fig 4**; ‘Subunits’). Within the gp120 subunit, there was a stronger decrease in mutational tolerance for the regions that did not belong to any variable loops (gp120-conserved) than there was for the variable loops (gp120-variable), although both conserved and variable regions exhibited a loss of mutational tolerance (**Fig 4**; ‘Conserved versus variable’). Among the five variable loops of gp120, the more conserved V3 loop exhibited the strongest negative net site diffsel (**Fig 4**; ‘gp120 Variable loops’) [57]. Further notable within the V3 loop, we observed a particularly large decrease in mutational tolerance for sites that are highly conserved, such as the GPGR motif or the hydrophobic patch whose disruption causes gp120 shedding (**Fig 5A** and **5B**) [58]. Indeed, across the entire Env protein positions with high Shannon entropy exhibited increases in mutational tolerance more frequently than conserved positions (**S7 Fig**). These observations indicate that conserved regions in Env generally experience stronger selection pressure when the ER proteostasis network is upregulated than do variable regions.

**Fig 5.**
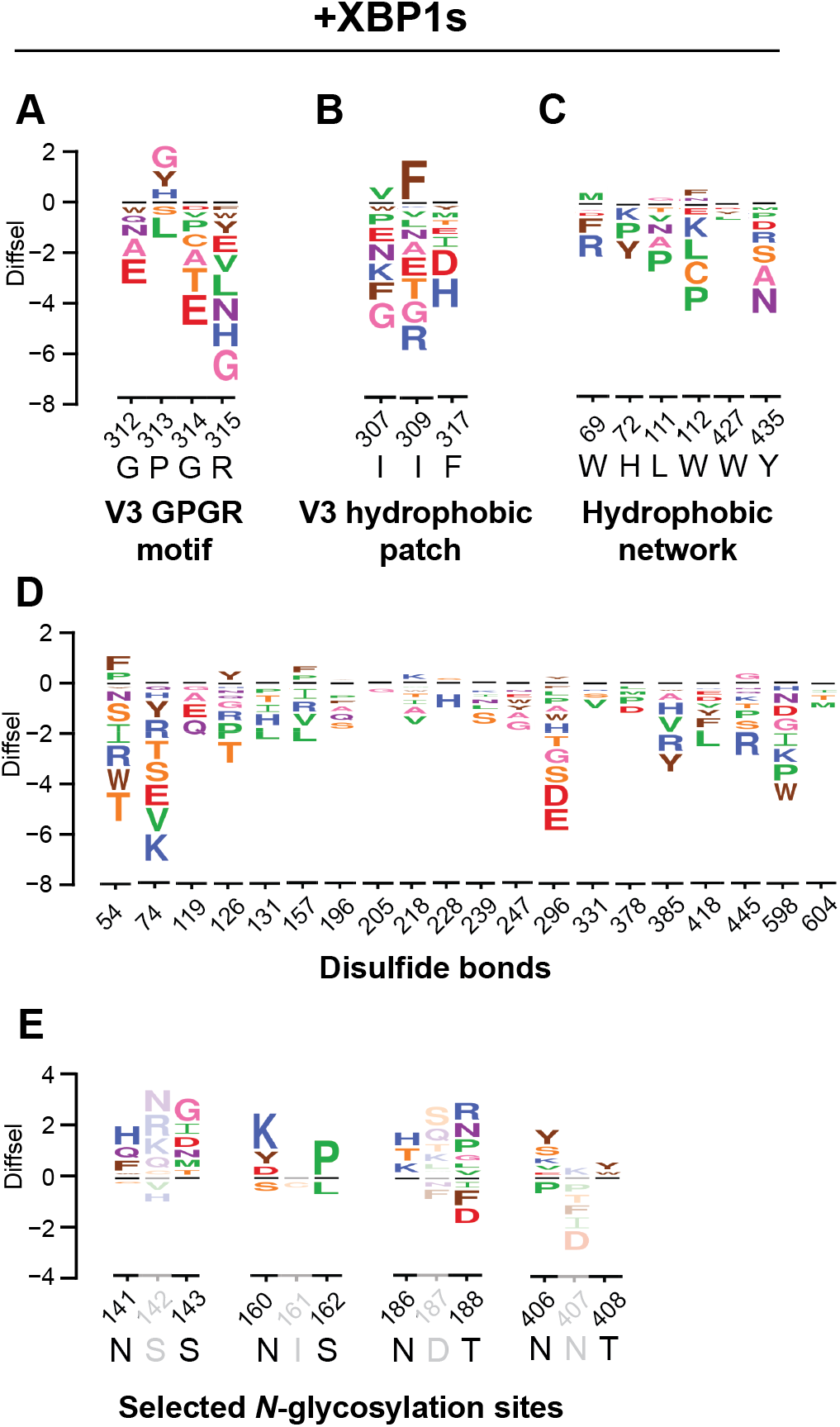
Diverse functional elements of Env respond differently to XBP1s induction. Selected sequence logo plots for the +XBP1s ER proteostasis environment normalized to the basal ER proteostasis environment for (**A**) the conserved GPGR motif of the V3 loop, (**B**) the hydrophobic patch of the V3 loop, (**C**) the hydrophobic network of gp120 important for CD4 binding, (**D**) selected *N*-glycosylation sequons (N-X-S/T) that exhibited positive net site diffsel in all three remodeled proteostasis environments and (**E**) cysteine residues participating in disulfide bonds. The height of the amino-acid abbreviation corresponds to the magnitude of diffsel. The numbers and letters below the logos indicate the Env site in HXB2 numbering and the wild-type amino acid for that site, respectively. Only variants that were present in all three pre-selection viral libraries and exhibited diffsel in the same direction across the biological triplicates are plotted. All logo plots were generated on the same scale.

We next scrutinized Env regions directly involved in membrane fusion, since the principal function of Env in the HIV replication cycle is to facilitate host cell entry via the fusion of viral and host membranes. Briefly, upon binding to cell surface CD4 receptor and coreceptor, the fusion peptide in gp41 is inserted into the cell membrane and the two heptad repeat domains form a three-stranded coiled-coil that allows the anchoring of Env to the host cell membrane [59]. With the exception of CD4 contact sites, regions participating in membrane fusion (**Fig 4**; ‘Membrane fusion’) experienced decreased mutational tolerance upon XBP1s induction. In addition, the hydrophobic network of gp120 that undergoes conformational changes upon CD4 binding to trigger membrane fusion [60] exhibited negative net site diffsel (**Fig 5C**).

Lastly, we focused further attention on regions of Env that may play important roles in Env folding and stability. We observed a significant decrease in mutational tolerance for sites participating in the gp120–gp41 subunit contact (**Fig 4**; ‘Subunit contact’). Next, we asked what the consequences of XBP1s induction are for disulfide bonds and *N*-glycosylation sequons. Particularly noteworthy, we observed that every single cysteine residue involved in disulfide bonds exhibited negative net site diffsel upon XBP1s induction (**Fig 4**; ‘Disulfide bond’ and **Fig 5D**), consistent with the notion that the XBP1s-remodeled ER proteostasis environment quite strictly quality controls disulfide bond formation in Env. The results were different for *N*-glycosylation sequons, even though these residues can also promote ER protein folding and quality control by providing access to the ER’s lectin-based chaperone network [61]. We observed an approximately equal number of sites in *N*-glycosylation sequons that displayed positive and negative net site diffsel upon XBP1s induction (**Fig 4**; ‘*N*-glycosylation’). In fact, several *N*-glycosylation sequons displayed positive net site diffsel across all three enhanced ER proteostasis environments (**Fig 5E**, **S8 Fig E** and **S8 Fig J**). Among those *N*-glycosylation sequons displaying positive net site diffsel, all except N160 are highly variable [62]. These observations add to the evidence that mutational tolerance is more strongly constrained in conserved regions than in variable regions upon upregulation of the host’s ER proteostasis machinery.

The trends observed for the simultaneous induction of XBP1s and ATF6 largely overlapped with XBP1s induction only (**S5 Fig** and **S8 Fig A–E**), except that CD4 contact sites exhibited a statistically significant decrease in mutational tolerance whereas subunit contact sites did not. Consistent with the less striking reduction in mutational tolerance observed upon ATF6 induction (**Fig 2C**), we observed that the impact of ATF6 induction was minimal across Env sites when we assessed structural/functional groups independently (**S6 Fig** and **S8 Fig F–J**). Only the gp41 subunit exhibited a small, yet statistically significant decrease in mutational tolerance (**S6 Fig**; ‘Subunits’).

Finally, to evaluate structural regions whose mutational tolerance was particularly impacted by host ER proteostasis network remodeling, we mapped net site diffsel values onto the Env crystal (**Fig 6**). Whereas mutationally intolerant sites were distributed throughout the Env trimer, sites with enhanced mutational tolerance upon XBP1s induction were located primarily at the apex of the Env trimer (**Fig 6A**). For instance, N160, S128, and D185 were among the sites with the highest positive net site diffsel. Indeed, although the magnitude of enhanced mutational tolerance varied, these sites exhibited positive net site diffsel in all host ER proteostasis conditions tested. N160, S128, and D185 had similar net site diffsel values when XBP1s was induced (**Fig 6A**) or when XBP1s and ATF6 were induced simultaneously (**Fig 6B**), but N160 exhibited substantially higher mutational tolerance when ATF6 was induced (**Fig 6C**). Notably, N160 belongs to the V2 apex, a well-characterized epitope targeted by the broadly neutralizing antibodies PG9 [63], CH01 [64], CAP256.09 [65], and PGT145 [66], and elimination of the N160 glycan was shown to confer antibody escape [37]. In addition, I165K, a fusion peptide inhibitor resistance mutation [64], was the single variant with the highest positive diffsel when XBP1s and ATF6 were induced simultaneously and the third highest positive diffsel when XBP1s was induced alone. These observations suggest that upregulation of host ER proteostasis factors, although generally constraining Env mutational tolerance, can still strongly enhance mutational tolerance in regions of the Env protein in which adaptive mutations are essential, including mutations at certain antibody- or drug-targeted regions of Env.

**Fig 6.**
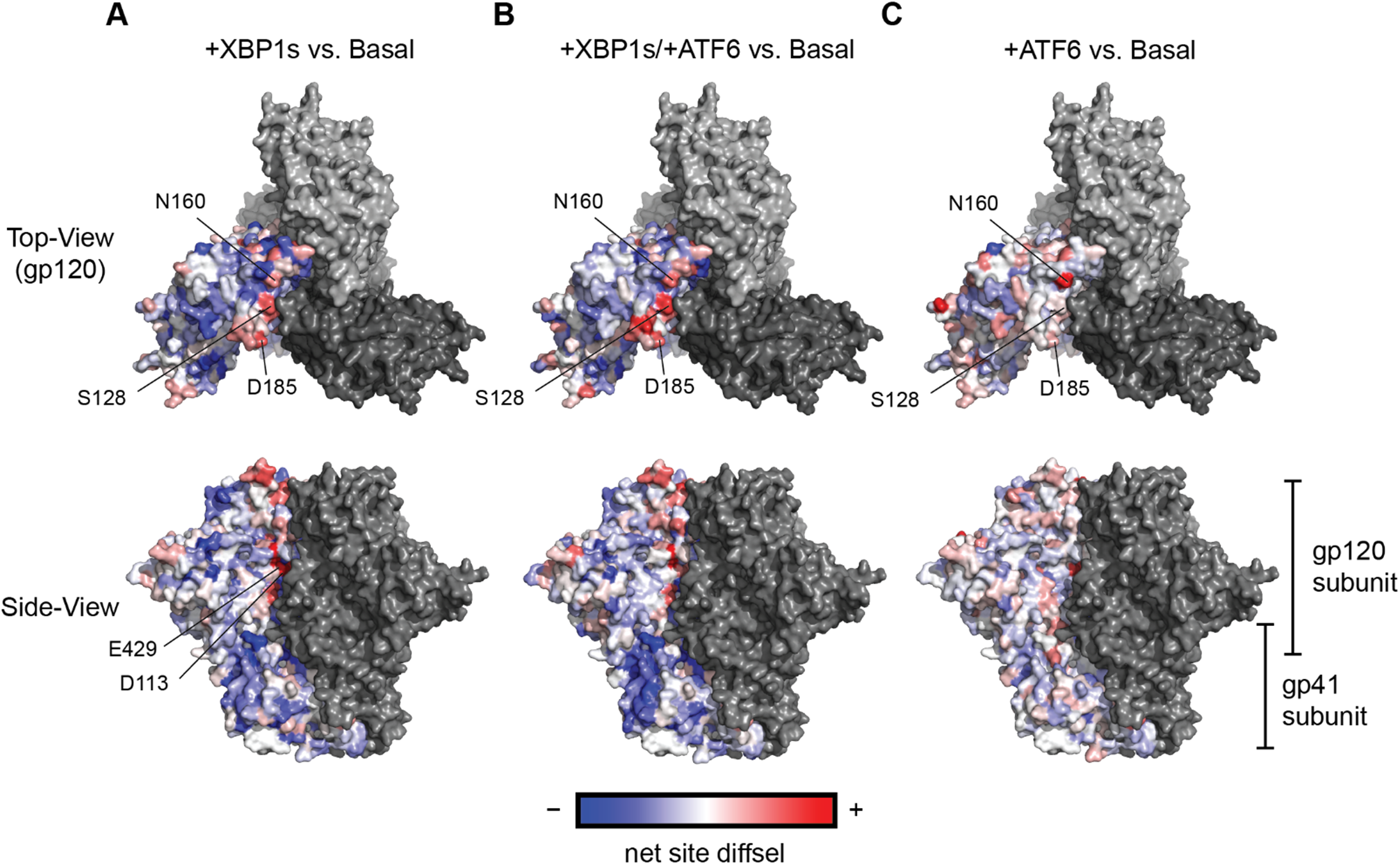
Env sites with positive net site diffsel are clustered at the trimer apex. Average net site diffsel values across Env for (**A**) +XBP1s (**B**) +XBP1s/+ATF6, and (**C**) +ATF6, normalized to the basal ER proteostasis environment, are mapped onto Env trimer crystal structure (PDBID 5FYK) [87]. One monomer is colored using net site diffsel as the color spectrum; negative net site diffsel residues are colored in blue, and positive net site diffsel residues are colored in red. The remainder of the Env trimer is colored in grey.

## Discussion

Our results provide the first experimental evidence that UPR-mediated upregulation of ER proteostasis network can globally reduce the mutational tolerance of client proteins. The primary ER proteostasis factors involved in driving this effect in Env are XBP1s-regulated, as the broad-scale effects of ATF6 activation are more muted (**Fig 2C** and **2F**).

Importantly, this observation is consistent with the impacts of cellular quality control factors on protein mutational tolerance, where the available protein sequence space is restricted through degradation and reduced trafficking of aberrantly folded protein variants [16–18]. Previous studies established that Env is readily targeted to and degraded by ERAD [27, 28, 67], suggesting that destabilizing Env variants may be subjected to more rapid removal by quality control factors in an enhanced ER proteostasis environment. Indeed, conserved regions of Env exhibit particularly large decrease in mutational tolerance upon XBP1s induction (**Fig 4** and **S5 Fig**), where mutations are more likely to cause protein misfolding. We note that the LAI strain of HIV used in this study could have had lower mutational tolerance than HIV strains on average, as it was isolated from a chronically infected individual and potentially accumulated a significant number of deleterious mutations. In future studies, it will be interesting to examine effects in additional HIV strains.

This work augments the emerging evidence that host ER proteostasis machinery can fundamentally define the mutational tolerance of viral membrane proteins. Prior to this study, the consequences of ER proteostasis network composition for the mutational tolerance of a membrane protein, whether viral or endogenous, had only ever been investigated for one other protein – influenza hemagglutinin [32]. We show that the host ER proteostasis network also impacts Env mutational tolerance, implying the potential that this relationship is applicable across multiple RNA viruses. Moreover, the present work reveals that the interaction between host proteostasis and viral proteins is highly nuanced, and the outcome can differ for each viral pathogen. For example, hemagglutinin mutational tolerance is enhanced at febrile temperatures upon XBP1s activation, with very minimal effects at a permissive temperature [32]. Unlike hemagglutinin, the majority of Env sites exhibited strongly decreased mutational tolerance upon upregulation of host ER proteostasis factors, in this case even at a permissive temperature.

Looking deeper into our observations for Env itself, this study highlights several Env regions that merit further investigation with respect to their roles in Env folding and structure. For example, we found that sites that constitute *N*-glycosylation sequons exhibited both positive and negative net site diffsel (**Fig 5E**, **S8 Fig E**, and **S8 Fig J**). While *N*-glycans in Env are important for antibody shielding and viral replication [68–71], there have been various reports on whether *N*-glycans are indispensable for proper folding of Env [71, 72]. The specific *N*-glycan sites that proved particularly sensitive to XBP1s upregulation are likely to play some important role in the folding, quality control, and/or trafficking of Env. It will be interesting to explore the specific biophysical mechanisms underlying our observations in future work.

Finally, we find that different sites within a single viral protein can respond differently to the selection pressure imposed by the host ER proteostasis network (**Fig 4**, **Fig 5**, **S2 Fig**, **S3 Fig**, and **S8 Fig**). Contrary to the global trend in decreased mutational tolerance, we observed many Env sites with positive net site diffsel, especially at the trimer apex of Env (**Fig 6**). We discovered that N160, where a glycan is installed that is obligatory for binding of the vast majority of V2 apex broadly neutralizing antibodies [73], exhibited enhanced mutational tolerance in all three proteostasis environments and particularly when ATF6 was induced alone. We also observed that I165K, an Env variant known to be fusion peptide inhibitor resistant [64], exhibited highly positive diffsel upon XBP1s induction. These observations indicate that, although the majority of Env sites exhibited depletion of variants, important antibody- or drug-escape variants may be enriched upon upregulation of host ER proteostasis network mechanisms. Thus, the host ER proteostasis environment can strongly influence the mutational tolerance of specific Env variants that are of therapeutic interest.

In conclusion, our results establish that stress response-mediated upregulation of proteostasis networks can actually restrict rather than increase accessible client protein sequence space, in contrast to most prior work focused on the effects of individual chaperones. We also find that evolutionary interactions between viral proteins and host proteostasis factors are specific to the virus type, as well as to specific regions of the viral protein. We anticipate this knowledge will prove particularly valuable for ongoing efforts to target host proteostasis network components for antiviral therapeutics [52, 74–78] and for the design of proteostasis network-targeted therapeutic adjuvants that can prevent the emergence of viral variants that confer immune system escape or drug resistance. More broadly, the principles observed here seem likely to prove generally applicable, not just to viral proteins but also endogenous client proteins

## Materials and Methods

### Data availability

RNA-Seq data are available in GEO with accession number GSE171356. The Python script used to perform DMS data analysis and generate the sequence logo plots is provided in a series of IPython notebooks in (https://github.com/yoon-jimin/2021_HIV_Env_DMS). FASTQ files from DMS sequencing are available from the Sequence Read Archive (http://www.ncbi.nlm.nih.gov/sra; SRP314168; BioProject PRJNA720817).

### Cell culture

Human T lymphoblasts (SupT1 cells; ATCC) were grown in RPMI-1640 medium (Corning), supplemented with 10% heat-inactivated fetal bovine serum (FBS, Cellgro), 1% penicillin/streptomycin/glutamine (Cellgro) at 37 °C with 5% CO_2_(g). TZM-bl reporter cells (NIH AIDS Research and Reference Reagent Program; Cat. no. 1470) were cultured in DMEM (Corning) supplemented with 10% heat-inactivated FBS, 1% penicillin/streptomycin/glutamine at 37 °C with 5% CO_2_(g). Cell lines were periodically tested for mycoplasma using the MycoSensor PCR Assay Kit (Agilent).

### Plasmids to engineer SupT1^DAX^ cells

The following lentiviral destination vectors were used for stable cell line construction: pLenti6/V5 Dest Gateway with a tetracycline repressor insert (Invitrogen) and blasticidin resistance, pLenti CMV/TO Zeocin DEST with either human XBP1s insert (Addgene), and pLenti CMV hygromycin DEST with a DHFR.ATF6(1-373) fusion, as previously described [41].

### Stable cell line engineering

For the construction of SupT1^DAX^ cells, SupT1 cells were first transduced with lentivirus encoding a blasticidin-resistant tetracycline repressor and then with lentivirus encoding zeocin resistant XBP1s. Transduction was performed by spinoculation with 2 μg/mL polybrene (Sigma-Aldrich) at 1,240 × *g* for 1–1.5 h. Heterostable cell lines expressing the tetracycline repressor and XBP1s were then selected using 10 μg/mL blasticidin (Gibco) and 50 μg/mg zeocin (Invitrogen). Single colony lines were derived from the heterostable population by seeding 30–40 cells in a 96-well plate in 100 μl of RPMI media without antibiotics for 10–14 days. Clonal populations were then selected and expanded in 24-well plates in 500 μL of RPMI containing 10 μg/mL blasticidin and 50 μg/mL zeocin. Cells were grown to confluency and then screened based on functional testing of the XBP1s construct using RT-PCR (described below) with or without 2 μg/mL doxycycline (dox; Alfa Aesar). The selected SupT1 single colony cell line encoding tetracycline-inducible XBP1s was then transduced with lentivirus encoding DHFR.ATF6(1–373) via the spinoculation protocol described above and stable cells were selected using 400 μg/mL hygromycin B (Gibco). The heterostable populations were then treated with vehicle, 2 μg/mL dox, 10 μM trimethoprim (TMP; Alfa Aesar), or 2 μg/mL dox and 10 μM trimethoprim and screened for function using qPCR (described below) to obtain the final stably engineered SupT1^DAX^ cell line.

### qPCR

SupT1^DAX^ cells were seeded at a density of 2 × 10^5^ cells/well in a 6-well plate in RPMI media and treated with 0.01 % DMSO, 2 μg/mL dox, 10 μM TMP, or 2 μg/mL dox and 10 μM TMP for 18 h. As a positive control for unfolded protein response activation, SupT1^DAX^ cells were treated with 10 μg/mL tunicamycin (Tm, Sigma-Aldrich) for 6 h. Cellular RNA was harvested using the Omega RNA Extraction Kit with Homogenizer Columns (Omega Bio-tek). 1 μg RNA was used to prepare cDNA using random primers (total reaction volume = 20 μL; Applied Biosystems High-Capacity Reverse Transcription Kit). The reverse transcription reaction was diluted to 80 μL with water, and 2 μL of each sample was used for qPCR with 2 × Sybr Green (Roche) and primers for human *RPLP2* (housekeeping gene), *HSPA5* (BIP), *HSP90B* (GRP94), *DNAJB9* (ERDJ4), and *SEC24D* (**S1 Data**). For qPCR data analysis, all gene transcripts were normalized to that of *RPLP2* and the fold-change in expression relative to DMSO-treated cells was calculated.

### RNA-Seq

SupT1^DAX^ cells were seeded in a 6-well plate (Corning) at a density of 5 x 10^5^ cells/well in RPMI media in quadruplicate. The cells were treated with 0.01 % DMSO, 2 μg/mL dox, 10 μM TMP, or 2 μg/mL dox and 10 μM TMP for 24 h. Cellular RNA was harvested using the RNeasy Plus Mini Kit with QIAshredder homogenization columns (Qiagen). RNA-Seq libraries were prepared using the Kapa mRNA HyperPrep RNA-seq library construction kit (Kapa/Roche), with 6 min fragmentation at 94 °C and nine PCR cycles of final amplification and duplex barcoding. Libraries were quantified using the Fragment Analyzer and qPCR before being sequenced on an Illumina HiSeq 2000 using 40-bp single-end reads in High Output mode.

Analyses were performed using previously described tools and methods [79]. Reads were aligned against hg19 (Feb., 2009) using bwa mem v. 0.7.12-r1039 [RRID:SCR_010910] with flags –t 16 –f, and mapping rates, fraction of multiply-mapping reads, number of unique 20-mers at the 5’ end of the reads, insert size distributions and fraction of ribosomal RNAs were calculated using bedtools v. 2.25.0 [RRID:SCR_006646] [80]. In addition, each resulting bam file was randomly down-sampled to a million reads, which were aligned against hg19, and read density across genomic features were estimated for RNA-Seq-specific quality control metrics. For mapping and quantitation, reads were aligned against GRCh38/ENSEMBL 89 annotation using STAR v. 2.5.3a with the following flags-runThreadN 8 –runMode alignReads –outFilter-Type BySJout –outFilterMultimapNmax 20 –alignSJoverhangMin 8 –alignSJDBoverhangMin 1 –outFilterMismatchNmax 999 – alignIntronMin 10 –alignIntronMax 1000000 –alignMatesGapMax1000000 –outSAMtype BAM SortedBy-Coordinate –quantMode TranscriptomeSAM with –genomeDir ointing to a 75nt-junction GRCh38 STAR suffix array [81]. Gene expression was quantitated using RSEM v. 1.3.0 [RRID:SCR_013027] with the following flags for all libraries: rsem-calculate-expression –calc-pme –alignments -p 8 –forward-prob 0 against an annotation matching the STAR SA reference [82]. Posterior mean estimates (pme) of counts and estimated RPKM were retrieved.

For differential expression analysis, dox-, TMP-, or dox and TMP-treated SupT1^DAX^ cells were compared against vehicle-treated SupT1^DAX^ cells. Differential expression was analyzed in the R statistical environment (R v.3.4.0) using Bioconductor’s DESeq2 package on the protein-coding genes only [RRID:SCR_000154] [83]. Dataset parameters were estimated using the estimateSizeFactors(), and estimateDispersions() functions; read counts across conditions were modeled based on a negative binomial distribution, and a Wald test was used to test for differential expression (nbinomWaldtest(), all packaged into the DESeq() function), using the treatment type as a contrast. Shrunken log_2_ fold-changes were calculated using the lfcShrink function. Fold-changes and *p*-values were reported for each protein-coding gene. Gene ontology analyses were performed using the online DAVID server, according to tools and methods presented by Huang *et al* [79]. The volcano plots were generated using EnhancedVolcano (**Fig 1B–D**; https://github.com/kevinblighe/EnhancedVolcano).

### Gene Set Enrichment Analysis (GSEA)

Differential expression results from DESeq2 were retrieved, and the “stat” column was used to pre-rank genes for GSEA analysis. These “stat” values reflect the Wald’s test performed on read counts as modeled by DESeq2 using the negative binomial distribution. Genes that were not expressed were excluded from the analysis. GSEA (desktop version, v3.0) [47, 84] was run in the pre-ranked mode against MSigDB 7.0 C5 (Gene Ontology) set, using the official gene symbol as the key, with a weighted scoring scheme, normalizing by meandiv, with 8958 gene sets retained, and 5000 permutations were run for *p*-value estimation. Selected enrichment plots were visualized using a modified version of ReplotGSEA, in R (https://github.com/PeeperLab/Rtoolbox/blob/master/R/ReplotGSEA.R).

### Resazurin metabolism assay

SupT1^DAX^ cells were seeded in 96-well plates (Corning) at a density of 1.5 × 10^5^ cells/well in RPMI media and then treated with 0.1% DMSO, 2 μg/mL dox, 10 μM TMP, or 2 μg/mL dox and 10 μM TMP. 72 h post-treatment, 50 μL RPMI containing 0.025 mg/mL resazurin sodium salt (Sigma) was added to the wells and mixed thoroughly. After 2 h of incubation, resorufin fluorescence (excitation 530 nm; emission 590 nm) was quantified using a Take-3 plate reader (BioTeK). Experiments were conducted in biological quadruplicate.

### HIV titering

TZM-bl reporter cells were seeded at a density of 2.5 × 10^4^ cells/well in 48-well plates. After 5 h, the cells were infected with 100 μL of serially diluted infectious HIV viral inoculum containing 10 μg/ml polybrene. Each sample was used to infect four technical replicates. After 48 h, the viral supernatant was removed and the cells were washed twice with PBS and then fixed with 4% paraformaldehyde (Thermo Scientific) for 20 min. The fixed cells were washed twice with PBS and then stained with 4 mM potassium ferrocyanide, 4 mM ferricyanide, and 0.4 mg/mL 5-bromo-4-chloro-3-indolyl-*p*-d-galactopyranoside (X-Gal) in PBS at 37 °C for 50 min. The cells were washed with PBS, blue cells were counted manually under a microscope, and infectious titers were calculated based on the number of blue cells per volume of viral inoculum.

### Deep mutational scanning

Three biological replicate HIV libraries were generated from three previously prepared independent Env mutant plasmid libraries (a generous gift from Prof. Jesse Bloom, University of Washington) following the previously reported protocol [22]. For DMS, SupT1^DAX^ cells were seeded in T175 vented tissue culture flasks (Corning) at a density of 1.0 × 10^8^ cells/flask in RPMI media. The cells were pre-treated with 0.01% DMSO, 2 μg/mL dox, 10 μM TMP, or 2 μg/mL dox and 10 μM TMP for 18 h. Pre-treated cells were infected with the p1viral libraries at a MOI of 0.005 based on the infectious (TZM-bl) titers. In addition, one flask was either mock-infected (negative control) or infected with wild-type virus (to enable error correction for DMS data analysis). To remove unbound virions from culture, 6 h post-infection the cells were pelleted at 2,000 rpm for 5 min, washed twice with 25 mL PBS, and then resuspended in 50 mL of RPMI media treated with 0.01% DMSO, 2 μg/mL dox, 10 μM TMP, or 2 μg/mL dox and 10 μM TMP. Cell pellets were harvested 96 h post-infection by centrifuging the culture at 2,000 rpm for 5 min. Cell pellets were washed twice with PBS and then resuspended in 1 mL of PBS. Aliquots (100 μL) were added to Eppendorf tubes and stored at –80 °C for subsequent DNA extraction.

To generate samples for Illumina sequencing, non-integrated viral DNA was purified from aliquots of frozen SupT1^DAX^ cells using a mini-prep kit (Qiagen) and ~10^7^ cells per prep. PCR amplicons of Env were prepared from plasmid or mini-prepped non-integrated viral DNA by PCR following a previously described protocol [22]. The amplicons were sequenced using barcoded-subamplicon sequencing, dividing Env into nine rather than the previously reported six sub-amplicons. We note that it was necessary to exclude Env amino acid residues 31–34 from analysis because, after PCR optimization, we were unable to identify functional primers for the first sub-amplicon that did not include these sites. As previously described, at least 10^6^ Env molecules were PCR-amplified for preparation of sub-amplicon sequencing libraries to ensure sufficient sampling of viral library diversity [55]. Briefly, this sequencing library preparation method appends unique, random barcodes and part of the Illumina adapter to Env subamplicon molecules. In a second round of PCR, the complexity of the uniquely barcoded subamplicons was controlled to be less than the sequencing depth, and the remainder of the Illumina adapter was appended. The resulting libraries were sequenced on an Illumina HiSeq 2500 in rapid run mode with 2 × 250 bp paired-end reads. The primers used are described in **S1 Data**.

### Deep mutational scanning data analysis

The software dms_tools2 (https://jbloomlab.github.io/dms_tools2/) [56] was used to align the deep-sequencing reads, count the number of times each codon mutation was observed both before and after selection, calculate the diffsel for each Env variant, and generate sequence logo plots (**Fig 3A**, **S2 Fig**, and **S3 Fig**). The IPython notebook for code to perform this analysis is provided in **S1 File**, as well as in https://github.com/yoon-jimin/2021_HIV_Env_DMS. SAA was calculated via PDBePISA (**S4 Fig**) [85] using the crystal structure of BG505 SOSIP.664 (PDBID 5V8M) [86] and aligning to the LAI Env sequence (**S3 Table**). PDBePISA calculates the solvent-accessible surface area of the monomer (‘ASA’) and the solvent-accessible surface area that is buried upon formation of interfaces (‘Buried surface in interfaces’). ‘Buried surface in interfaces’ values were subtracted from ‘ASA’ values to obtain the SAA of each residue. Ligands and antibodies were removed from the PDB file prior to SAA analysis. Site entropy (Shannon entropy) was calculated using the Los Alamos HIV Sequence Database Shannon Entropy-One tool (**S7 Fig**). The calculation was based on the consensus sequence generated from the 7590 HIV-1 Env sequences in the Los Alamos HIV Sequence Database (one sequence per patient up to 2019). The net site diffsel values were mapped onto Env crystal structure (PDBID 5FYK) [87] using PyMOL (**Fig 6**).

### Statistical analyses

Unless indicated otherwise, experiments were performed in biological triplicate with replicates defined as independent experimental entireties (i.e., from plating the cells to acquiring the data). For deep mutational scanning, each biological replicate mutant viral library was prepared from independently generated mutant plasmid libraries, as previously reported [55]. Diffsel values from deep mutational scanning were tested for significance of deviation from zero (no relative enrichment or depletion), using a one-sample *t*-test in Graph Pad Prism. For diffsel values and net site diffsel values, two-tailed *p*-values are reported to assess whether the mean (net site) diffsel for enhanced ER proteostasis environments were significantly different from zero (**Fig 2C** and **2F**). For net site diffsel distributions for specific functional and structural groups, *p*-values were Bonferroni-corrected for 18 tests (**Fig 4**, **S5 Fig**, and **S6 Fig**).

## Acknowledgements

The authors are grateful to Prof. Jesse Bloom and Dr. Hugh Haddox for helpful discussion. This work was supported by a Kwanjeong Graduate Fellowship (to JY), a UNCF-Merck Postdoctoral Fellowship (to EEN), National Science Foundation Graduate Research Fellowships (to AMP and SJH), Tufts University (to YSL), the National Science Foundation CAREER Award (MCB1652390 to MDS), and the National Institutes of Health (1R35GM136354 to MDS). Additional support was provided by the National Cancer Institute Koch Institute Support (core) Grant P30-CA14051 and the National Institute for Environmental Health Sciences MIT Center for Environmental Health Sciences (core) Grant P30-ES002109.

---

**S1 Fig.**
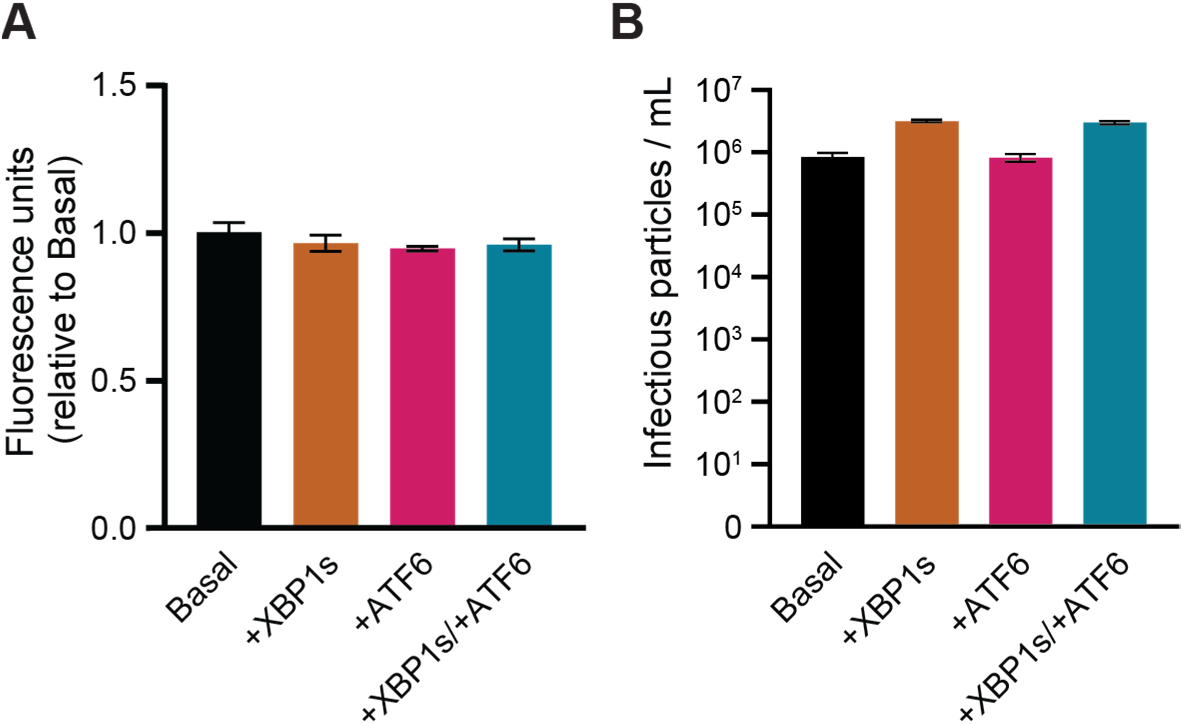
ER proteostasis perturbation has no deleterious effects on cell viability and does not restrict HIV replication. (**A**) Induction of XBP1s, ATF6, or simultaneous induction of XBP1s and ATF6 did not alter metabolic activity of SupT1 cells, as measured by a resazurin assay. The average of biological quadruplicates is plotted with error bars representing the standard error of mean (SEM). (**B**) Induction of XBP1s and simultaneous induction of XBP1s and ATF6 did not restrict and actually slightly increased HIV infectious titers, while induction of ATF6 did not influence HIV replication in SupT1 cells, as measured by TZM-bl infectious units. The average of biological triplicates is plotted with error bars representing the SEM.

**S2 Fig.**
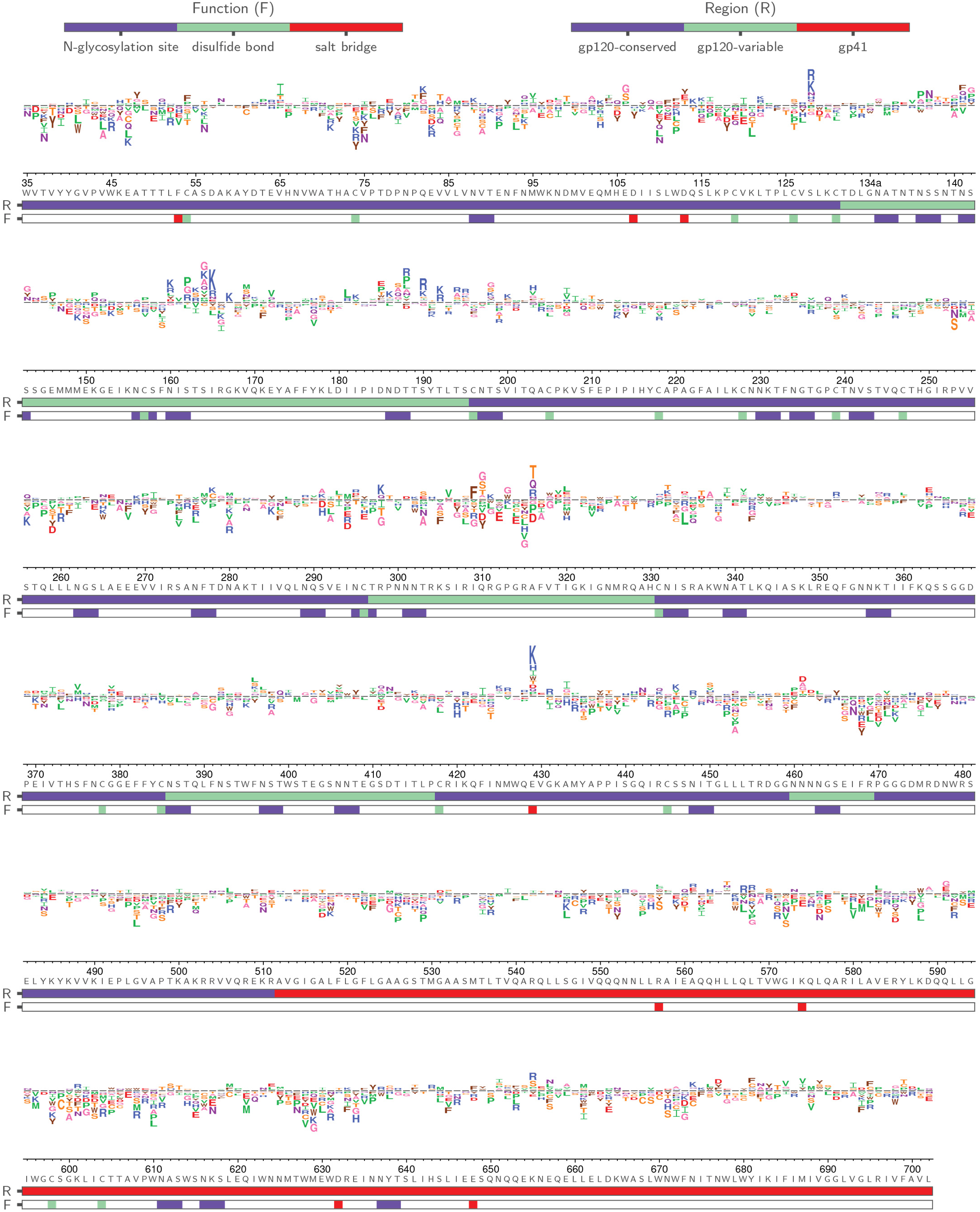
Sequence logo plots reveal diffsel across Env upon simultaneous induction of XBP1s and ATF6. Logo plot displaying averaged diffsel for +XBP1s/+ATF6 normalized to the basal proteostasis environment. The height of the amino-acid abbreviation corresponds to the magnitude of diffsel. The amino-acid abbreviations are colored based on the side-chain properties: negatively charged (D, E; red), positively charged (H, K R; blue), polar uncharged (C, S, T; yellow), small nonpolar (A, G; pink), aliphatic (I, L, M, P, V; green), and aromatic (F, W, Y; brown). The numbers and letters below the logos indicate the Env site in HXB2 numbering and the identity of the wild-type amino acid for that site, respectively. The color bar below the logos indicates the function (F) that the site is involved in (*N*-glycosylation site (blue), disulfide bond (green), or salt bridge (red)) or the region (R) of Env that the site belongs to (gp120-conserved (blue), gp120-variable (green), or gp41 (red); the sites that belong to the five variable loops of gp120 were categorized as ‘gp120-variable’, and the sites that are not included in the five variable loops were categorized as ‘gp120-conserved’). Only variants that were present in all three preselection viral libraries and exhibited diffsel in the same direction across all three biological triplicates are plotted here. Unfiltered logo plots for each individual replicate are provided in **S2 File**.

**S3 Fig.**
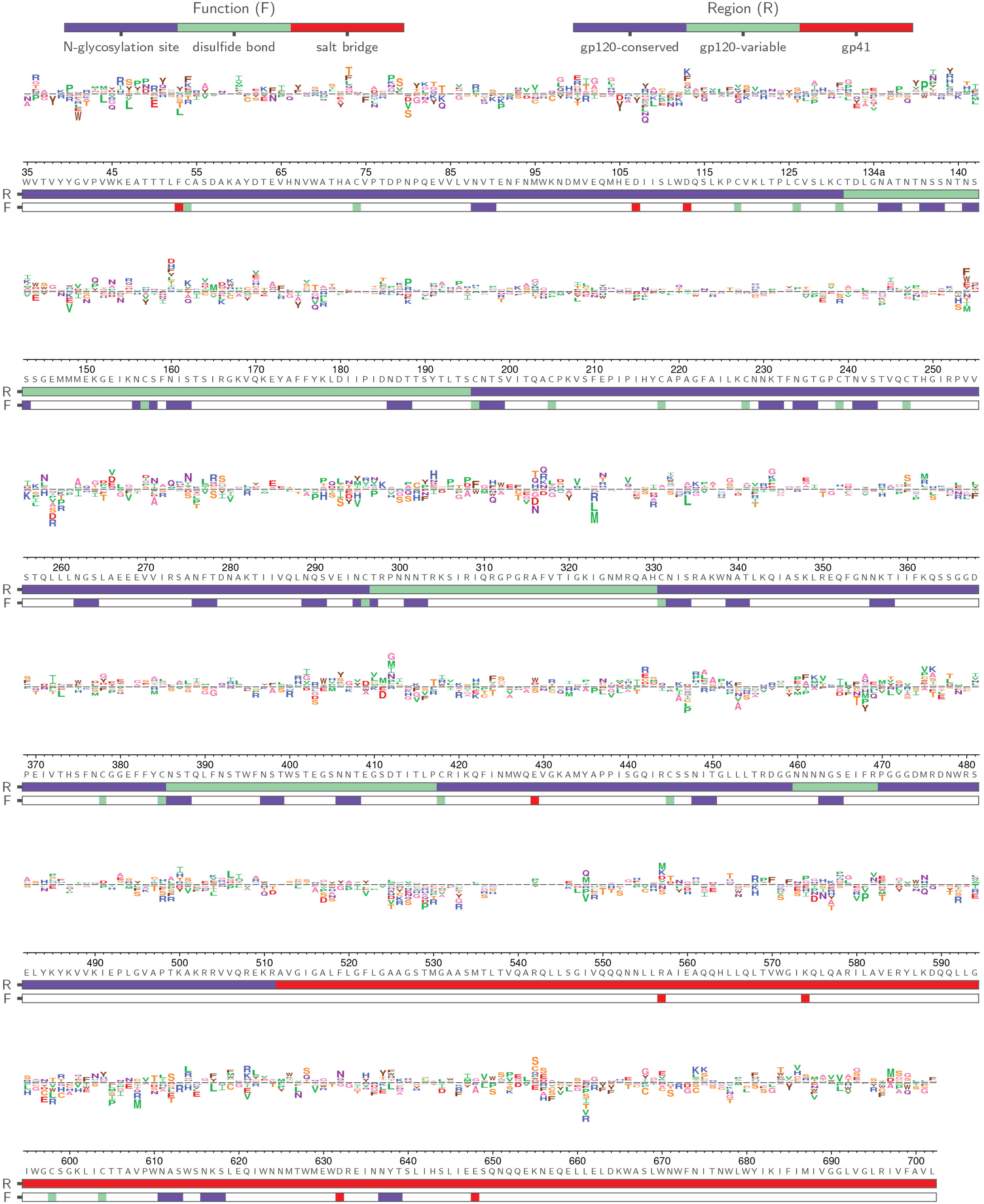
Sequence logo plots reveal diffsel across Env upon induction of ATF6. Logo plot displaying averaged diffsel for +ATF6 normalized to the basal proteostasis environment. The height of the amino-acid abbreviation corresponds to the magnitude of diffsel. The amino-acid abbreviations are colored based on the side-chain properties: negatively charged (D, E; red), positively charged (H, K R; blue), polar uncharged (C, S, T; yellow), small nonpolar (A, G; pink), aliphatic (I, L, M, P, V; green), and aromatic (F, W, Y; brown). The numbers and letters below the logos indicate the Env site in HXB2 numbering and the identity of the wild-type amino acid for that site, respectively. The color bar below the logos indicates the function (F) that the site is involved in (*N*-glycosylation site (blue), disulfide bond (green), or salt bridge (red)) or the region (R) of Env that the site belongs to (gp120-conserved (blue), gp120-variable (green), or gp41 (red); the sites that belong to the five variable loops of gp120 were categorized as ‘gp120-variable’, and the sites that are not included in the five variable loops were categorized as ‘gp120-conserved’). Only variants that were present in all three pre-selection viral libraries and exhibited diffsel in the same direction across all three biological triplicates are plotted here. Unfiltered logo plots for each individual replicate are provided in **S2 File**.

**S4 Fig.**
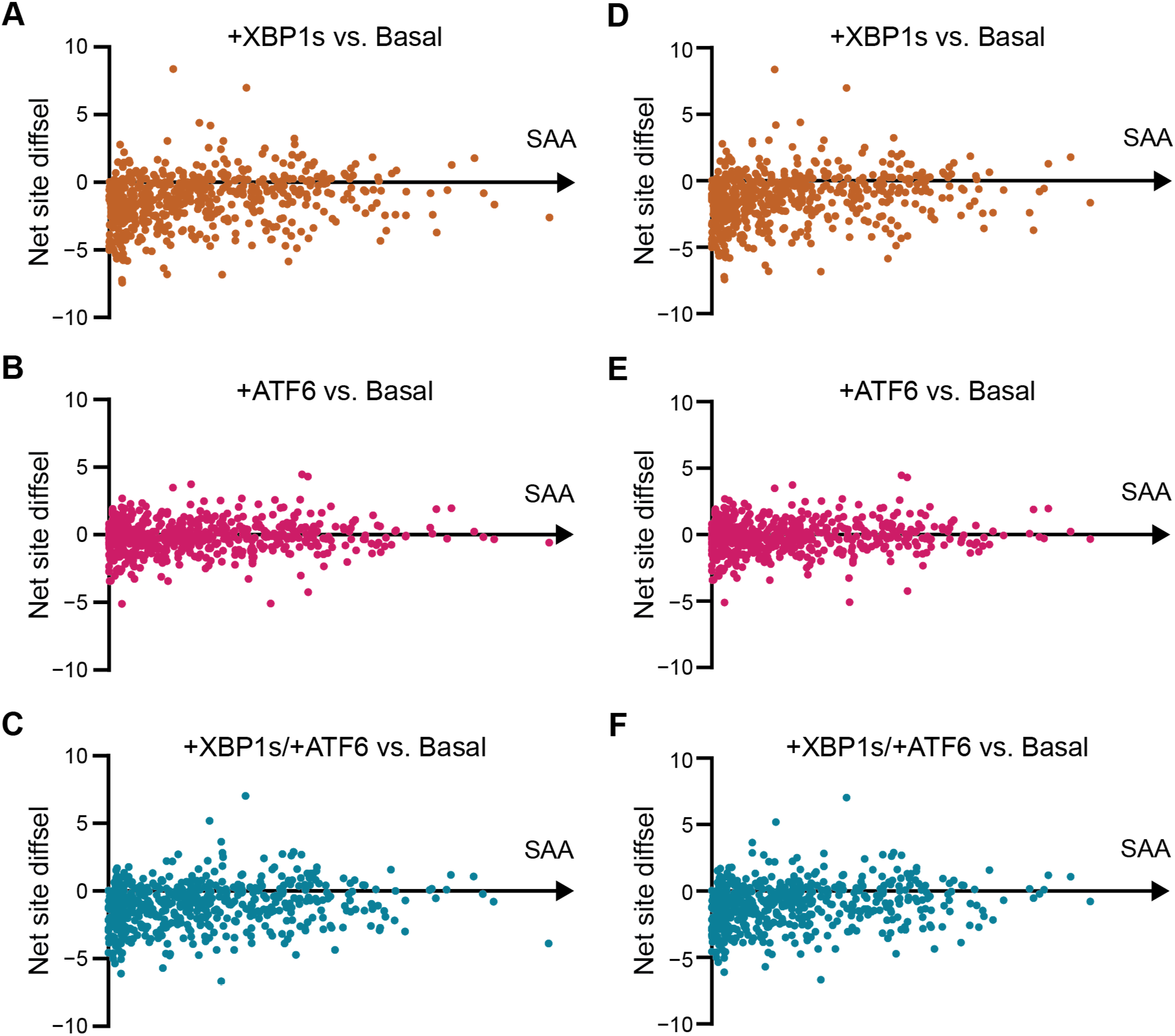
Env net site diffsel is not correlated with surface accessible area (SAA). Average net site diffsel values plotted against the SAA of Env monomer (**A–C**) and trimer (**D–F**). Average net site diffsel values for (**A**, **D**) +XBP1s, (**B**, **E**) +ATF6, and (**C**, **F**) +XBP1s/+ATF6 were normalized to the basal ER proteostasis environment and plotted against the SAA at each site. SAA was calculated using PDBePISA [85] with PDBID 5V8M [86], where SAA = 0 corresponds to a buried site.

**S5 Fig.**
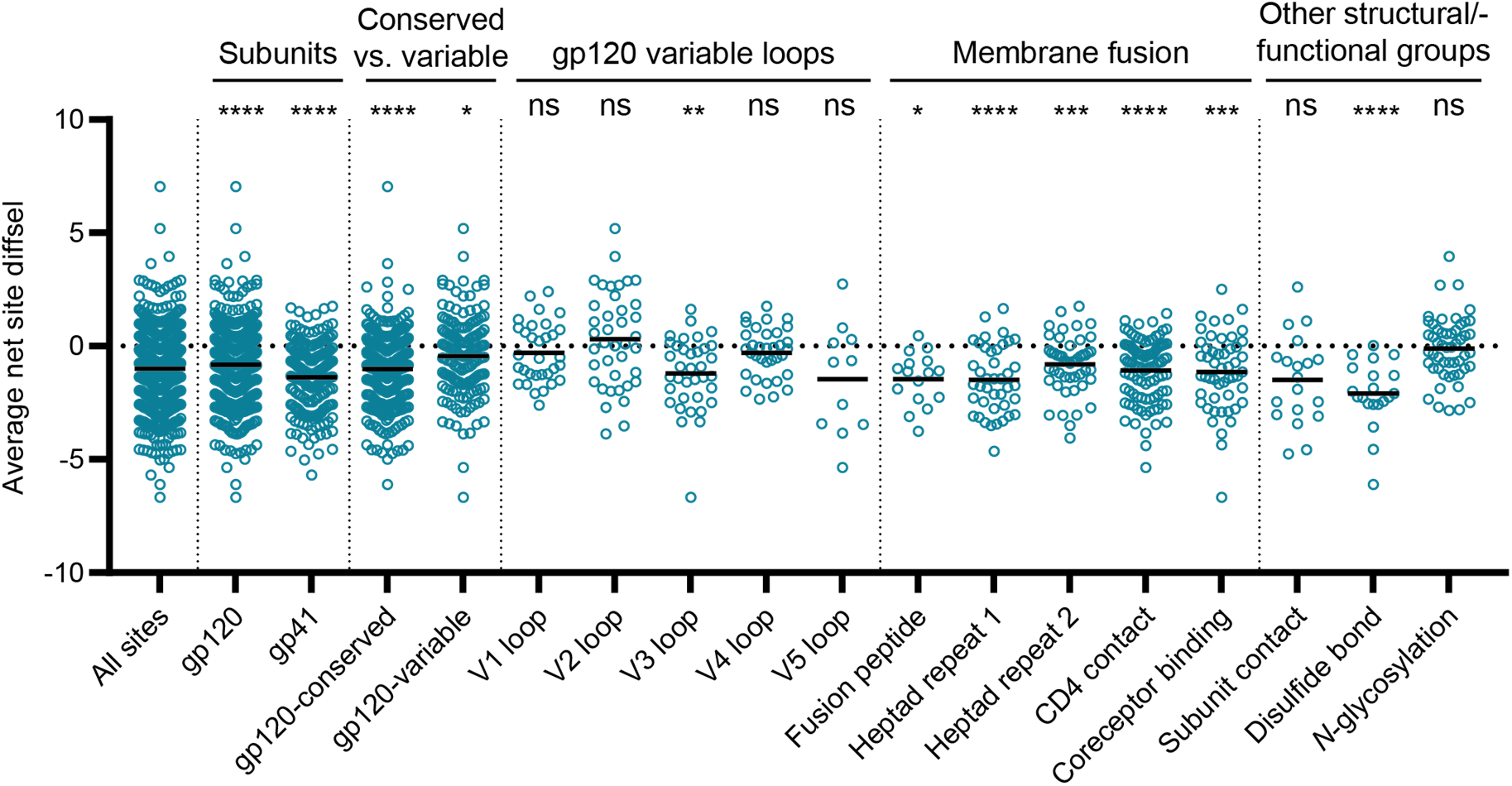
Impact of combined induction of XBP1s andATF6 on mutational tolerance varies across Env structural elements. Average net site diffsel for the +XBP1s/+ATF6 ER proteostasis environment normalized to the basal ER proteostasis environment, where medians of distributions are indicated by black horizontal lines. Sites are sorted by subunits, conserved vs. variable regions of gp120, five variable loops of gp120, regions important for membrane fusion, and other structural/functional groups. For ‘Conserved vs. variable’, the sites that belong to the five variable loops of gp120 were categorized as ‘gp120-variable’, and the sites that are not included in the five variable loops were categorized as ‘gp120-conserved’. Significance of deviation from null (net site diffsel = 0, no selection) was tested using a one-sample Ltest. The derived *p*-values were Bonferroni-corrected for 18 tests and *, **, ***, and **** represent adjusted two-tailed *p*-values of <0.05, <0,01, <0.001, and <0.0001.

**S6 Fig.**
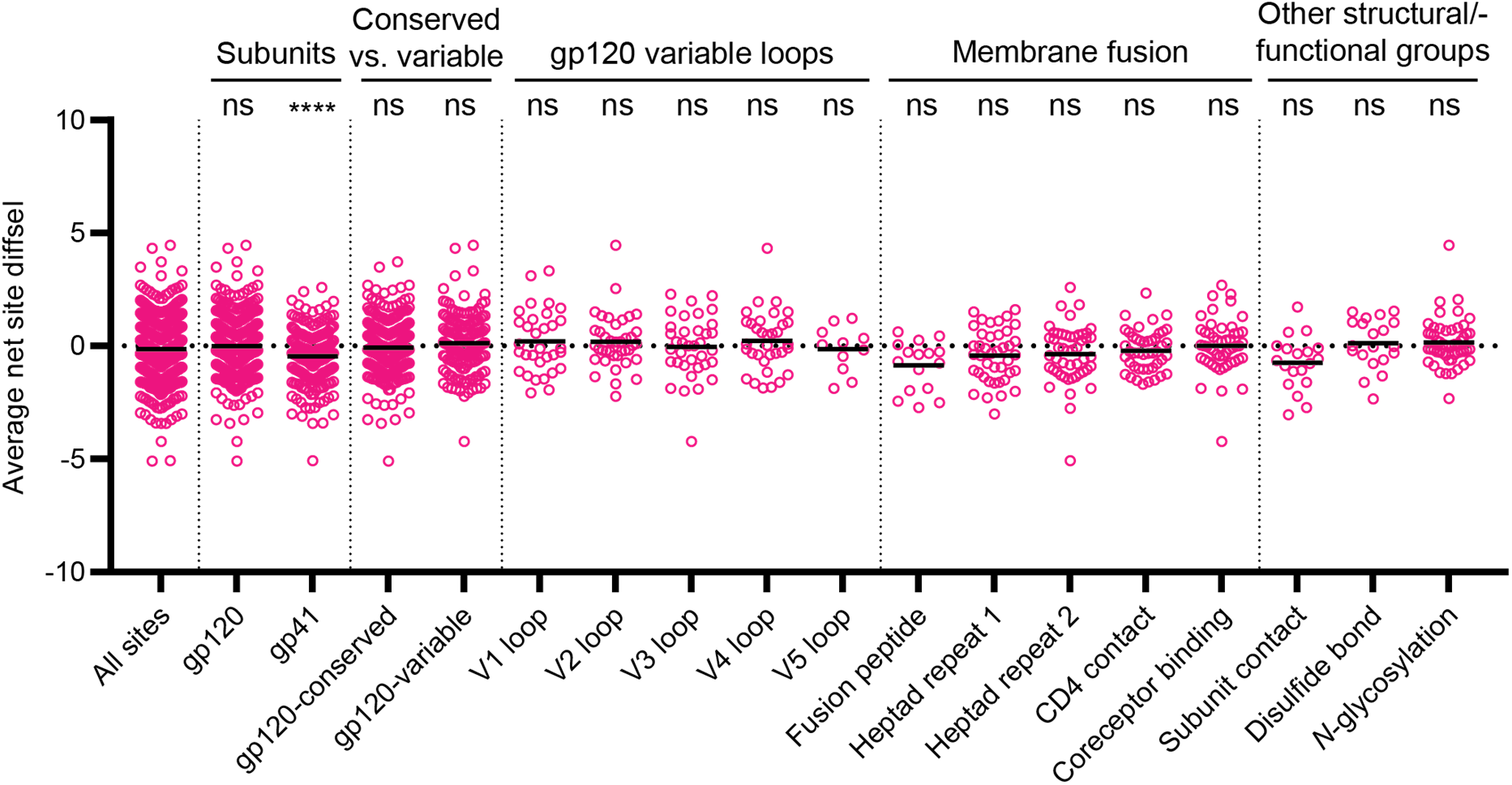
Impact of ATF6 induction on mutational tolerance varies across Env structural elements. Average net site diffsel for the +ATF6 ER proteostasis environment normalized to the basal ER proteostasis environment, where medians of distributions are indicated by black horizontal lines. Sites are sorted by subunits, conserved vs. variable regions of gp120, five variable loops of gp120, regions important for membrane fusion, and other structural/functional groups. For ‘Conserved vs. variable’, the sites that belong to the five variable loops of gp120 were categorized as ‘gp120-variable’, and the sites that are not included in the five variable loops were categorized as ‘gp120-conserved’. Significance of deviation from null (net site diffsel = 0, no selection) was tested using a one-sample *t*-test. The derived *p*-values were Bonferroni-corrected for 18 tests and *, **, ***, and **** represent adjusted two-tailed *p*-values of <0.05, <0,01, <0.001, and <0.0001.

**S7 Fig.**
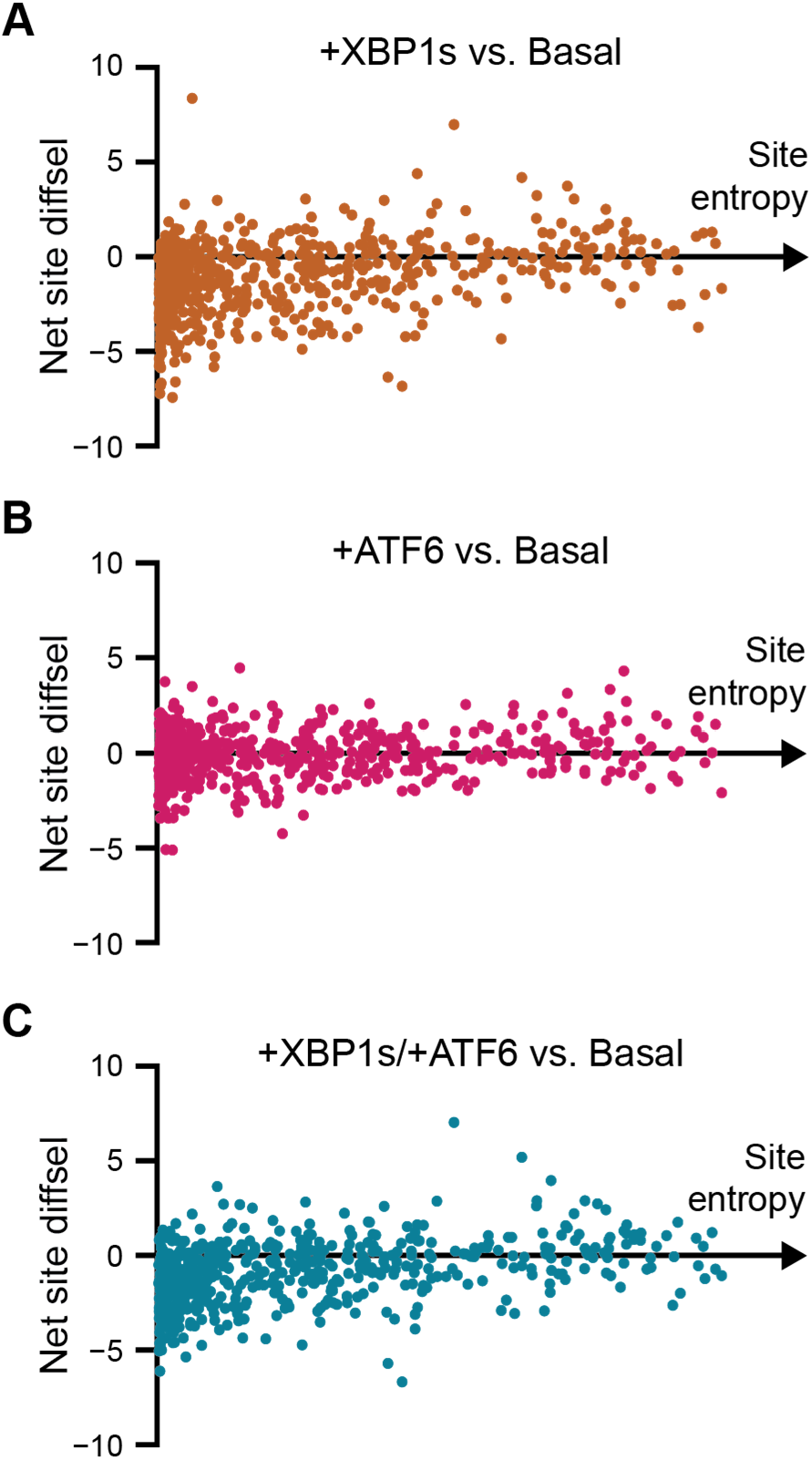
Enhanced mutational tolerance is observed more frequently at sites with high site entropy. Average net site diffsel values across Env for (**A**) +XBP1s (**B**) +ATF6, and (**C**) +XBP1s/+ATF6 are normalized to the basal ER proteostasis environment and plotted against the site entropy at each site.

**S8 Fig.**
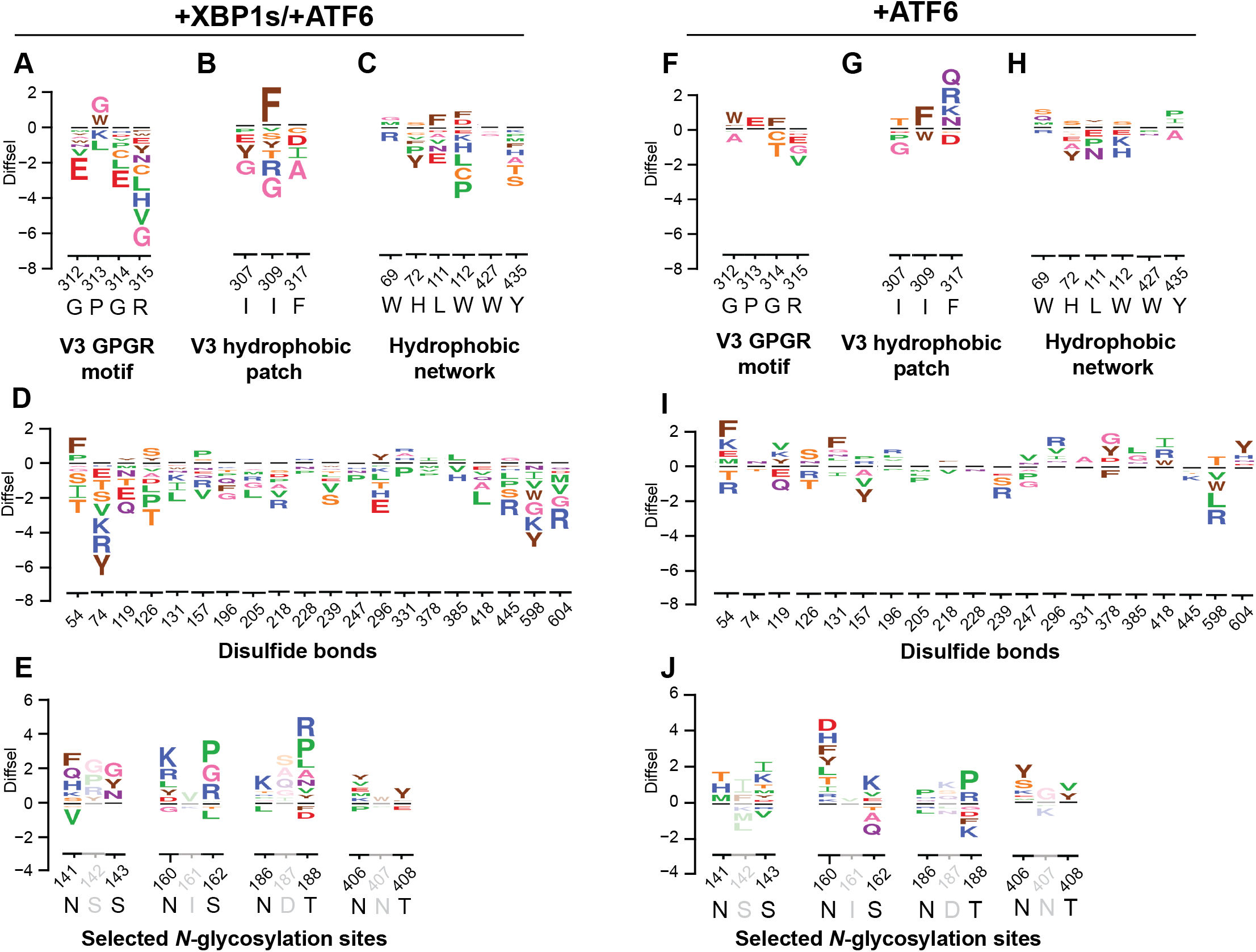
Diverse functional elements of Env respond differently to combined induction of XBP1s and ATF6, and ATF6 induction. Selected sequence logo plots for the +XBP1s/+ATF6 (**A–E**) and +ATF6 (**F–J**) ER proteostasis environments normalized to the basal ER proteostasis environment for (**A**, **F**) the conserved GPGR motif of the V3 loop, (**B**, **G**) the hydrophobic patch of the V3 loop, (**C**, **H**) the hydrophobic network of gp120 important for CD4 binding, (**D**, **I**) selected *N*-glycosylation sequons (N-X-S/T) that exhibited positive net site diffsel in all three remodeled proteostasis environments, and (**E**, **J**) cysteine residues participating in disulfide bonds. The height of the amino-acid abbreviation corresponds to the magnitude of diffsel. The numbers and letters below the logos indicate the Env site in HXB2 numbering and the wild-type amino acid for that site, respectively. Only variants that were present in all three pre-selection viral libraries and exhibited diffsel in the same direction across the biological triplicates are plotted. All logo plots were generated on the same scale.

