## Supplementary material for "The Host Cell’s Endoplasmic Reticulum Proteostasis Network Profoundly Shapes the Protein Sequence Space Accessible to HIV Envelope": analysis_notebook.ipynb

```
# %% [markdown]
# The host cell's ER proteostasis network profoundly shapes the protein sequence space accessible to HIV envelope

# %% [markdown]
## Overview

The first part of this analysis processes FASTQ files generated by Barcoded-subamplicon sequencing and converts them to codon counts, which are the frequencies of mutations at each site of Env. The analysis primarily uses [dms2_batch_bcsubamp](https://jbloomlab.github.io/dms_tools2/dms2_batch_bcsubamp.html).

The second part of this analysis calculates [differential selection](https://jbloomlab.github.io/dms_tools2/diffsel.html#diffsel) (diffsel) from codoncounts.csv files. Briefly, differential selection quantifies the relative enrichment of each amino acid substitution at a given site in the selection condition compared to the mock condition. The analysis primarily uses [dms2_batch_diffsel](https://jbloomlab.github.io/dms_tools2/dms2_batch_diffsel.html).

# %% [markdown]
### Import modules and define general variables

# %% [code]
import os
import sys
import glob
import pandas
from IPython.display import IFrame
from IPython.display import display, HTML
import dms_tools2
import dms_tools2.plot
import dms_tools2.sra
import dms_tools2.utils
import dms_tools2.diffsel
from dms_tools2.ipython_utils import showPDF

print("Using dms_tools2 version {0}".format(dms_tools2.__version__))

# results will go in this directory
resultsdir = './results/' 
if not os.path.isdir(resultsdir):
    os.mkdir(resultsdir)
    
# CPUs to use, use all available
ncpus = -1

# use existing results if available
use_existing = 'yes'

# %% [markdown]
### Define the samples

We define samples needed for processing the Barcoded-subamplicon deep sequencing data. We first create a list of sampleIDs, which are the names of the FASTQ files. We then create a list of sampleNames, which are short descriptions of the FASTQ files and also what the output files will be named. Last, we create a list of R1 (forward read) and R2 (reverse read) files.

We use the following abbreviations to name our samples:

 * Mt1, Mt2, and Mt3 are used to indicate biological replicate 1, 2, and 3, respectively

 * D, T, DT, and V are used to indicate dox-treated (+XBP1s), TMP-treated (+ATF6), dox- and TMP-treated (+XBP1s/+ATF6). and vehicle-treated (Basal) samples, respectively

 * pLAI is used to indicate p0 mutant viral library

 * WT is used to indicate wild-type Env

# %% [code]
# Directory for FASTQ files
fastqdir = '/Users/sjhendel/Documents/FASTQ_Files/HIV_Env_DMS_EN'

# List of sampleIDs
sampleIDs = ['171013Sho_D17-193001','171013Sho_D17-193002','171013Sho_D17-193003',
             '171013Sho_D17-193004','171013Sho_D17-193005','171013Sho_D17-193006',
             '171013Sho_D17-193007','171013Sho_D17-193008','171013Sho_D17-193009',
             '171013Sho_D17-193010','171013Sho_D17-193011','171013Sho_D17-193012',
             '171013Sho_D17-193013','171013Sho_D17-193014','171013Sho_D17-193015',
             '171013Sho_D17-193016','171013Sho_D17-193017']

# List of sampleNames
sampleNames = ['Mt1-V','Mt1-D','Mt1-T','Mt1-DT','Mt2-V','Mt2-D','Mt2-T','Mt2-DT',
                   'Mt3-V','Mt3-D','Mt3-T','Mt3-DT','Wt-V','pLAI-Wt','pLAI-Mt1','pLAI-Mt2','pLAI-Mt3']

# List of R1 and R2 files
r1filename = []
r2filename = []

fastqdir = "/Users/sjhendel/Documents/FASTQ_Files/HIV_Env_DMS_EN/"

for i in range(len(sampleIDs)):
    sampleID = sampleIDs[i]
    r1filename.append('/Users/sjhendel/Documents/FASTQ_Files/HIV_Env_DMS_EN/'+sampleID+'_1_sequence.remerged.fastq')
    r2filename.append('/Users/sjhendel/Documents/FASTQ_Files/HIV_Env_DMS_EN/'+sampleID+'_2_sequence.remerged.fastq')

samples = pandas.DataFrame(list(zip(sampleIDs, sampleNames, r1filename, r2filename)),
               columns =['sampleID', 'name', 'R1', 'R2'])

display(HTML(samples.to_html(index=False)))

# %% [markdown]
### Run dms2_batch_bcsubamp

# %% [code]
# file containing wildtype LAI Env sequence
refseq = './HIV_LAI_Env_reference_seq.fa'
# Specify R1 and R2 read lengths and gene range of each subamplicon for dms_barcodedsubamplicons
alignspecs_ori = ['103,336,45,42',
                  '337,561,45,42',
                  '520,795,43,36',
                  '772,1002,42,42',
                  '994,1221,42,39',
                  '1186,1422,45,39',
                  '1414,1653,39,39',
                  '1651,1902,36,42',
                  '1897,2136,36,36']

alignspecs = ' '.join(alignspecs_ori)
alignspecs = alignspecs.strip()

# raw counts and alignments placed in this directory
rawcountsdir = os.path.join(resultsdir, 'rawcodoncounts')
if not os.path.isdir(rawcountsdir):
    os.mkdir(rawcountsdir)
    
# write sample information to a batch file for dms2_batch_bcsubamplicons
countsbatchfile = os.path.join(rawcountsdir, 'batch.csv')
print("Here is the batch file that we write to CSV format to use as input:")
display(HTML(samples[['name', 'R1', 'R2']].to_html(index=False)))
samples[['name', 'R1', 'R2']].to_csv(countsbatchfile, index=False)

# %% [code]
print('\nNow running dms2_batch_bcsubamp...')
log = !dms2_batch_bcsubamp \
            --batchfile {countsbatchfile} \
            --refseq {refseq} \
            --alignspecs {alignspecs} \
            --outdir {rawcountsdir} \
            --summaryprefix summary \
            --R1trim {200} \
            --R2trim {160} \
            --fastqdir {fastqdir} \
            --ncpus {ncpus} \
            --use_existing {use_existing} 
print("Completed dms2_batch_bcsubamp.")

# %% [markdown]
### Remove sites that are not mutagenized

In the Env mutant libraries we used, the N-terminal signal peptide and the C-terminal cytoplasmic tail were excluded from mutagenesis owing to their dramatic impact on Env expression and/or HIV infectivity. The .codoncounts files were manually modified to exclude these sites (sites 0-34, 708-722) and saved in the directory './results/codoncounts'.

# %% [code]
countsdir = './results/codoncounts'
countsfiles = os.path.join(countsdir, '*')

print("Here are the modified codoncounts files")

for name in glob.glob(countsfiles):
    print(name)

# %% [markdown]
### Run dms2_batch_diffsel

# %% [code]
# diffsel files placed in this directory
diffseldir = os.path.join(resultsdir, 'diffsel')
if not os.path.isdir(diffseldir):
    os.mkdir(diffseldir)

diffselbatchfile = os.path.join(diffseldir, 'batch.csv')

# Specify which samples to use as mock condition and selection condition to calculate differential selection
diffselbatch = pandas.DataFrame.from_records([
    ('XBP1s', 'replicate1', 'Mt1-D_codoncounts.csv', 'Mt1-V_codoncounts.csv'),
    ('ATF6', 'replicate1', 'Mt1-T_codoncounts.csv', 'Mt1-V_codoncounts.csv'),
    ('XBPsATF6', 'replicate1', 'Mt1-DT_codoncounts.csv', 'Mt1-V_codoncounts.csv'),
    ('XBP1s', 'replicate2', 'Mt2-D_codoncounts.csv', 'Mt2-V_codoncounts.csv'),
    ('ATF6', 'replicate2', 'Mt2-T_codoncounts.csv', 'Mt2-V_codoncounts.csv'),
    ('XBPsATF6', 'replicate2', 'Mt2-DT_codoncounts.csv', 'Mt2-V_codoncounts.csv'),
    ('XBP1s', 'replicate3', 'Mt3-D_codoncounts.csv', 'Mt3-V_codoncounts.csv'),
    ('ATF6', 'replicate3', 'Mt3-T_codoncounts.csv', 'Mt3-V_codoncounts.csv'),
    ('XBPsATF6', 'replicate3', 'Mt3-DT_codoncounts.csv', 'Mt3-V_codoncounts.csv'),
    ],
    columns = ['group', 'name', 'sel', 'mock']
    )

# Specify which sample to use as error correction
diffselbatch['err'] = 'Wt-V_codoncounts.csv'

print("Here is the batch file that we write to CSV format to use as input:")
display(HTML(diffselbatch.to_html(index=False)))
diffselbatch.to_csv(diffselbatchfile, index=False)

# %% [code]
print('\nNow running dms2_batch_diffsel...')
log = !dms2_batch_diffsel \
        --summaryprefix summary \
        --batchfile {diffselbatchfile} \
        --outdir {diffseldir} \
        --indir {countsdir} \
        --use_existing {use_existing}
print("Completed dms2_batch_bcsubamp.")
```
