## Supplementary material for "The Host Cell’s Endoplasmic Reticulum Proteostasis Network Profoundly Shapes the Protein Sequence Space Accessible to HIV Envelope": CodonsToAAs.py

```
import sys

def main():
    
    aa={0:'G',1:'A',2:'V',3:'I',4:'L',5:'M',6:'F',7:'Y',8:'W',9:'S',10:'T',11:'N',12:'Q',13:'C',14:'P',15:'H',16:'R',17:'K',18:'D',19:'E'}
    aa2={'G':0,'A':1,'V':2,'I':3,'L':4,'M':5,'F':6,'Y':7,'W':8,'S':9,'T':10,'N':11,'Q':12,'C':13,'P':14,'H':15,'R':16,'K':17,'D':18,'E':19}

    f1 = sys.argv[1] # Input file (Counts file with NNN codons, instead of amino acids)
    f2 = sys.argv[2] # Output file name
    
    length = 0
    with open(f1) as f:
        for line in f:
            length = length + 1
    print(length)
    aaCounts = [[0 for i in range(20)] for j in range(length-1)]
    
    with open(f1) as f:
        firstLine = f.readline().rstrip().split('\t') # Need to keep this in order to skip first line
        firstLine = "POSITION,WT,K,N,K,N,T,T,T,T,R,S,R,S,I,I,M,I,Q,H,Q,H,P,P,P,P,R,R,R,R,L,L,L,L,E,D,E,D,A,A,A,A,G,G,G,G,V,V,V,V,STOP,Y,STOP,Y,S,S,S,S,STOP,C,W,C,L,F,L,F".split(',')
        ## Comment this out if not needed
        print(len(firstLine))
        for lineText in f:
            line = lineText.split(',')
            
            for pos in range(2,len(line)):
                if(firstLine[pos] != "STOP"):
                    aaCounts[int(line[0])-1-34][aa2[firstLine[pos]]] += int(line[pos]) ## The 34 is a special case here, because the HIV Env sample begins at 35
    #print(aaCounts[0])    
    with open(f2,'w') as f:
        for line in aaCounts:
            for pos in range(len(line)):
                f.write(str(line[pos]))
                if(pos<len(line)-1):
                    f.write(',')
            f.write("\n")
    
main()
```
