## Supplementary material for "The Host Cell’s Endoplasmic Reticulum Proteostasis Network Profoundly Shapes the Protein Sequence Space Accessible to HIV Envelope": FilteringStep1.py

```
import sys

def main():
    
    ########### GLOBALS ###########
    
    aa={0:'G',1:'A',2:'V',3:'I',4:'L',5:'M',6:'F',7:'Y',8:'W',9:'S',10:'T',11:'N',12:'Q',13:'C',14:'P',15:'H',16:'R',17:'K',18:'D',19:'E'}
    
    aa2={'G':0,'A':1,'V':2,'I':3,'L':4,'M':5,'F':6,'Y':7,'W':8,'S':9,'T':10,'N':11,'Q':12,'C':13,'P':14,'H':15,'R':16,'K':17,'D':18,'E':19}

    
    ###############################
    
    
    
    
    
    
    ########### INPUTS ###########
    
    dmsdiffsel = sys.argv[1] ### THE DIFFSEL FILE THAT COMES FROM DMS_TOOLS2
    
    libAAcounts = sys.argv[2] ### THE AA COUNTS FILE FROM THE INITIAL LIBRARY
    
    heatmapfilename = sys.argv[3] ### THE HEATMAP FILE NAME
    
    ###############################
    


    
    ########### MAKE UNFILTERED HEATMAP ###########
    
    # Find length of the protein
    diffLength = 0
    with open(dmsdiffsel) as f:
        for line in f:
            diffLength = diffLength + 1
    diffLength = diffLength/20
    
    print("Protein Length: ",diffLength) ### Print length of the protein
       
    
    heatmap = [[0 for i in range(20)] for j in range(diffLength)] # empty heatmap to fill


    with open(dmsdiffsel) as f: # open the dms_tools2 diffsel file
        for textLine in f:
            line = textLine.split(',')
            if(line[0] != 'site'):
                heatmap[int(line[0])-1][aa2[line[2]]] = float(line[3].rstrip()) 
    
    ### Write the unfiltered heatmap


    with open('unfiltered_'+heatmapfilename,"w") as f:
        for line in heatmap:
            for pos in range(len(line)):
                f.write(str(line[pos]))
                if(pos<len(line)-1):
                    f.write(',')
            f.write("\n")         
            
    ################################################
    
    
    
    
    
    
    ########### MAKE FILTERED HEATMAP ###########

    with open(libAAcounts) as f: # Open library amino acid counts file 
        for i in range(diffLength): 
            line = f.readline().split(',')
            for pos in range(len(line)):
                if(line[pos]=='0'):# If the library has zero counts
                    heatmap[i][pos] = -100.00 # make the value -100.00
    
    
    ### Write the filtered heatmap
    
    with open(heatmapfilename,'w') as f:
        for line in heatmap:
            for pos in range(len(line)):
                f.write(str(line[pos]))
                if(pos<len(line)-1):
                    f.write(',')
            f.write("\n")    
            
    #############################################
    
main()
```
