## Supplementary material for "The Host Cell’s Endoplasmic Reticulum Proteostasis Network Profoundly Shapes the Protein Sequence Space Accessible to HIV Envelope": FilteringStep2.py

```
import sys

def main():
    
    ########### GLOBALS ###########
    
    aa={0:'G',1:'A',2:'V',3:'I',4:'L',5:'M',6:'F',7:'Y',8:'W',9:'S',10:'T',11:'N',12:'Q',13:'C',14:'P',15:'H',16:'R',17:'K',18:'D',19:'E'}
    
    aa2={'G':0,'A':1,'V':2,'I':3,'L':4,'M':5,'F':6,'Y':7,'W':8,'S':9,'T':10,'N':11,'Q':12,'C':13,'P':14,'H':15,'R':16,'K':17,'D':18,'E':19}

    
    ###############################
    
    
    
    
    
    
    ########### INPUTS ###########
    
    rep1 = sys.argv[1] ### HEATMAP 1
    
    rep2 = sys.argv[2] ### HEATMAP 2
    
    rep3 = sys.argv[3] ### HEATMAP 3
    
    pickoutName = sys.argv[4] ### Name for pickoutsFile
    
    ###############################
    


    
    # Find length of the protein
    diffLength = 0
    with open(rep1) as f:
        for line in f:
            diffLength = diffLength + 1
    
    print("Protein Length: ",diffLength) ### Print length of the protein
       
    rep1List = [] # The empty new lists
    rep2List = []
    rep3List = []
    
    
    
    
    # Filling the new lists with data from files 
    
    with open(rep1) as f:
        for textLine in f:
            rep1List.append(textLine.rstrip().split(','))
            
    with open(rep2) as f:
        for textLine in f:
            rep2List.append(textLine.rstrip().split(','))  
            
    with open(rep3) as f:
        for textLine in f:
            rep3List.append(textLine.rstrip().split(','))
    
    
    
    
    
    
    pickouts = []
    
    
    for i in range(len(rep1List)):
        for j in range(len(rep1List[i])):
            if(rep1List[i][j] == '-100.0' or rep2List[i][j] == '-100.0' or rep3List[i][j] == '-100.0'): # Checking if any of the three samples has -100.0 at a given position
                rep1List[i][j] = '-100.0'
                rep2List[i][j] = '-100.0'
                rep3List[i][j] = '-100.0'
            elif( (float(rep1List[i][j]) > 0 and float(rep2List[i][j]) > 0 and float(rep3List[i][j]) > 0) or (float(rep1List[i][j]) < 0 and float(rep2List[i][j]) < 0 and float(rep3List[i][j]) < 0) ): # Checking that all three samples go in the same direction
                # IF THEY DO, these variants are referred to as a "pickout"
                # These values will survive through the filtering
                pickouts.append([i,aa[j],(float(rep1List[i][j])+float(rep2List[i][j])+float(rep3List[i][j]))/3,float(rep1List[i][j]),float(rep2List[i][j]),float(rep3List[i][j])])
            else:   # Filtering these out if they are not all in the same direction
                rep1List[i][j] = '-100.0'
                rep2List[i][j] = '-100.0'
                rep3List[i][j] = '-100.0'
    
    # Writing new files
    
    with open("filtered_"+rep1,'w') as f:
        for line in rep1List:
            for pos in range(len(line)):
                f.write(str(line[pos]))
                if(pos<len(line)-1):
                    f.write(',')
            f.write("\n")

    with open("filtered_"+rep2,'w') as f:
        for line in rep2List:
            for pos in range(len(line)):
                f.write(str(line[pos]))
                if(pos<len(line)-1):
                    f.write(',')
            f.write("\n")    
            
    with open("filtered_"+rep3,'w') as f:
        for line in rep3List:
            for pos in range(len(line)):
                f.write(str(line[pos]))
                if(pos<len(line)-1):
                    f.write(',')
            f.write("\n")            

    with open(pickoutName,'w') as f:
        for line in pickouts:
            for pos in range(len(line)):
                f.write(str(line[pos]))
                if(pos<len(line)-1):
                    f.write(',')
            f.write("\n")
    #############################################
    
main()
```
