## Supplementary material for "The Host Cell’s Endoplasmic Reticulum Proteostasis Network Profoundly Shapes the Protein Sequence Space Accessible to HIV Envelope": README.txt

To decipher reliable signal from experimental noise, we filtered the DMS data using the following two-step strategy. This analysis
(1) removes all mutdiffsel values for variants not present in the starting library of the corresponding replicate, replacing that mutdiffsel value with -100 in an array which is then plotted on a heatmap (where -100 is colored in black).
(2) removes mutdiffsel values that are not of the same sign (positive or negative) across replicates. values that are removed are replaced with -100 in the "pickouts" files and are shaded white in the heatmaps.

########################################

#CodonsToAAs.py

This script converts codon counts file with NNN codons to amino acids, which is needed for downstream analysis.

The order of amino acids is: {0:'G',1:'A',2:'V',3:'I',4:'L',5:'M',6:'F',7:'Y',8:'W',9:'S',10:'T',11:'N',12:'Q',13:'C',14:'P',15:'H',16:'R',17:'K',18:'D',19:'E'}

Inputs:
    
    codoncounts ### The pLAI codoncounts file from dms2_batch_bcsubamp
    
    AAcountsname ### The output file name


To run:
	
    python CodonsToAAs.py codoncounts.csv AAcountsname.txt

########################################

#FilteringStep1.py

This script removes all mutdiffsel values for variants not present in the starting library of the corresponding replicate, replacing that mutdiffsel value with -100 in an array which is then plotted on a heatmap. This script creates unfiltered and filtered heatmaps.


Inputs:
    
    dmsdiffsel ### The diffsel file from dms2_batch_diffsel
    
    AAcountsname ### The amino acid counts file from the initial library

    heatmapfilename ### The heatmap file name
    

To run:

    python FilteringStep1.py dmsdiffsel.csv AAcountsname.txt heatmapfilename.txt

#FilteringStep2.py

This script removes mutdiffsel values that are not of the same sign (positive or negative) across replicates. Values that are removed are replaced with -100 in the "pickouts" files.

Inputs:
    
    heatmap1 ### Replicate 1 heatmap
    
    heatmap2 ### Replicate 2 heatmap

    heatmap3 ### Replicate 3 heatmap

    pickoutname ### The pickout file name

To run:

    python FilteringStep2.py heatmap1.txt heatmap2.txt heatmap3.txt pickoutname.txt

#FilteredDiffsel

    This folder contains the final pickout.txt files converted to csv files with column titles. This step was performed manually.
