## Supplementary material for "The Host Cell’s Endoplasmic Reticulum Proteostasis Network Profoundly Shapes the Protein Sequence Space Accessible to HIV Envelope": UnfilteredUnaveragedLogo.ipynb

```
# %% [markdown]
## Overview

We make sequence logo plots using unfiltered, unaveraged differential selection values

We add three overlayes: (1) wildtype sequence (2) site type (i.e. disulfides, *N*-glycosylation sequons, salt bridges), and (3) region (variable loops of gp120, non-variable loops of gp120, gp41)

Prior to generating sequence logo plots, the differential selection files were manually renumbered using [HXB2 numbering](http://people.biology.ucsd.edu/satish/Science/MePapers/korber.pdf). The renumbered diffsel files are used as the input files.

# %% [markdown]
To adjust the logo plots for the best visual clarity, we made the following adjustments to the source codes

<pre>/Users/sam/.local/lib/python3.7/site-packages/weblogolib/__init__.py

* self.logo_margin = 2
* self.stroke_width = 0.5
* self.tic_length = 8
* self.stack_width = std_sizes["medium"]
* self.stack_aspect_ratio = 3
* self.stack_margin = 0.5
* self.pad_right = False
* self.small_fontsize = 6
* self.fontsize = 20
* self.title_fontsize = 20
* self.number_fontsize = 5

/Users/sam/.local/lib/python3.7/site-packages/phydmslib/weblogo.py

* relativestackheight = 1.5
* separatorheight = 0.01
* stackwidth =  5
    *** a stack width any greater will cause letters bleeding over to neighboring sites
* stackaspectratio = 1

in jupyter make the following adjustments:

* --nperline 120 \
* --letterheight 8 \
* --diffselrange -9 9 \  (this may differ on the dataset being used)

# %% [markdown]
### Import modules and define general variables

# %% [code]
import os
from IPython.display import display, HTML
import dms_tools2
import dms_tools2.plot
import dms_tools2.sra
import dms_tools2.utils
import dms_tools2.diffsel
import phydmslib.weblogo
from dms_tools2.ipython_utils import showPDF
import matplotlib

print("Using dms_tools2 version {0}".format(dms_tools2.__version__))

#number of CPUs to use. Use all available
ncpus = -1
#use existing results if they exist
use_existing = 'yes'

# %% [markdown]
### Define input and output directories

# %% [code]
diffseldir = './diffsel'
overlaydir = './overlays'
logodir = './sequencelogo'
if not os.path.isdir(logodir):
    os.mkdir(logodir)

# %% [code]
for proteostasis in ['XBP1s', 'ATF6', 'XBP1sATF6']:
    for replicate in ['1', '2', '3']:
        name = proteostasis + '-replicate' + replicate
        mutdiffsel = os.path.join(diffseldir, name + '_diffsel.csv')
        logoplot = os.path.join(diffseldir, '{0}-replicate{1}_diffsel.pdf'.format(proteostasis, replicate))
        wtoverlayfile = os.path.join(overlaydir, proteostasis + '_wildtypeoverlay.csv')
        regionfile = os.path.join(overlaydir, proteostasis + '_regionoverlay.csv')
        sitetypefile = os.path.join(overlaydir, proteostasis + '_sitetypeoverlay.csv')
        print("\nCreating logo plot for {0} replicate{1} from {2}".format(proteostasis, replicate, mutdiffsel))
        log = !dms2_logoplot \
                --diffsel {mutdiffsel} \
                --name {name} \
                --outdir {logodir} \
                --overlay1 {wtoverlayfile} wildtype wildtype \
                --overlay2 {regionfile} R "Region" \
                --overlay3 {sitetypefile} F "Function" \
                --nperline 113 \
                --letterheight 8 \
                --diffselrange -20 20 \
                --underlay yes \
                --overlaycolormap rainbow \
                --ignore_extracols yes \
                --use_existing yes \
        
print("\nCompleted dms2_batch_seqlogo.")

# %% [markdown]
### +XBP1s vs. Basal replicate 1

# %% [code]
showPDF(logodir + '/XBP1s-replicate1_diffsel.pdf')

# %% [markdown]
### +XBP1s vs. Basal replicate 2

# %% [code]
showPDF(logodir + '/XBP1s-replicate2_diffsel.pdf')

# %% [markdown]
### +XBP1s vs. Basal replicate 3

# %% [code]
showPDF(logodir + '/XBP1s-replicate3_diffsel.pdf')

# %% [markdown]
### +ATF6 vs. Basal replicate 1

# %% [code]
showPDF(logodir + '/ATF6-replicate1_diffsel.pdf')

# %% [markdown]
### +ATF6 vs. Basal replicate 2

# %% [code]
showPDF(logodir + '/ATF6-replicate2_diffsel.pdf')

# %% [markdown]
### +ATF6 vs. Basal replicate 3

# %% [code]
showPDF(logodir + '/ATF6-replicate3_diffsel.pdf')

# %% [markdown]
### +XBP1s/+ATF6 vs. Basal replicate 1

# %% [code]
showPDF(logodir + '/XBP1sATF6-replicate1_diffsel.pdf')

# %% [markdown]
### +XBP1s/+ATF6 vs. Basal replicate 2

# %% [code]
showPDF(logodir + '/XBP1sATF6-replicate2_diffsel.pdf')

# %% [markdown]
### +XBP1s/+ATF6 vs. Basal replicate 3

# %% [code]
showPDF(logodir + '/XBP1sATF6-replicate3_diffsel.pdf')
```
