## Supplementary material for "The Host Cell’s Endoplasmic Reticulum Proteostasis Network Profoundly Shapes the Protein Sequence Space Accessible to HIV Envelope": README.txt

# Deep mutational scanning of HIV Env in remodelled host cell ER proteostasis network

(publication info goes here)

This repository is for the project by Jimin Yoon, Emmanuel Nekongo, and Matthew Shoulders (https://shoulderslab.mit.edu). This project analyzes the impact of the  host ER proteostasis network on the mutational tolerance of human immunodeficiency virus (HIV) envelope (Env). This analysis primarily uses [dms_tools2](https://jbloomlab.github.io/dms_tools2/index.html)

## Abbreviations used

 * Mt1, Mt2, and Mt3 are used to indicate biological replicate 1, 2, and 3, respectively

 * D, T, DT, and V are used to indicate dox-treated (+XBP1s), TMP-treated (+ATF6), dox- and TMP-treated (+XBP1s/+ATF6). and vehicle-treated (Basal) samples, respectively

 * pLAI is used to indicate p0 mutant viral library

 * WT is used to indicate wild-type Env

## Analysis
LAI site numberings are used for analysis steps 1, 2, 3. HXB2 site numberings are used for analysis steps 4-1 and 4-2

### 1. Calculating codon counts from FASTQ files

This analysis processes FASTQ files generated by Barcoded-subamplicon sequencing and converts them to codon counts, which are the frequencies of mutations at each site of Env. This analysis is performed by analysis_notebook.ipynb and primarily uses [dms2_batch_bcsubamp](https://jbloomlab.github.io/dms_tools2/dms2_batch_bcsubamp.html). dms2_batch_bcsubamp documentation also contains detailed explanation of the output files

This analysis requires the following input data

 * The FASTQ files from deep mutational scanning. The FASTQ files can be downloaded from the Sequence Read Archive under SRA project SRP314168 and BioProject PRJNA720817

 * The wildtype Env sequence for the LAI strain used in the experiment. The seqeucne file (LAI_Env_reference.fa) can be found in the main directory

This analysis generates the following output data. The output files can be found in ./results/rawcodoncounts and ./results/codoncounts

 * For each sample: log file, read statistics file, reads-per-barcode file, barcode statistics file, counts file

 * As summary statistics: log file, bcstats file, codonmuttypes file, codonntchanges file, cumulmutcounts file, depth file, mutfreq file, readsperbc file, readstats file, singlentchanges file

### 2. Calculating differential selection

This analysis calculates [differential selection (diffsel)](https://jbloomlab.github.io/dms_tools2/diffsel.html#diffsel) from codoncounts.csv files. Briefly, differential selection quantifies the relative enrichment of each amino acid substitution at a given site in the selection condition compared to the mock condition. This analysis is performed by analysis_notebook.ipynb and primarily uses [dms2_batch_diffsel](https://jbloomlab.github.io/dms_tools2/dms2_batch_diffsel.html). dms2_batch_diffsel documentation also contains detailed explanation of the output files

This analysis requires the following input data

 * codoncounts.csv files generated from dms2_batch_bcsubamp. The input files can be found in ./results/codoncounts

This analysis generates the following output data

 * For each sample: log file, mutdiffsel file, sitediffsel file

 * As summary statistics: meanmutdiffsel, meansitediffsel, medianmutdiffsel, mediansitediffsel, absolutesitediffselcorr, maxmutdiffselcorr, mutdiffselcorr, positivesitediffselcorr, meanmaxdiffsel, meanminmaxdiffsel, meanpositivediffsel, meantotaldiffsel, medianmaxdiffsel, medianminmaxdiffsel, medianpositivediffsel, mediantotaldiffsel

### 3. Filtering diffsel

This analysis filters diffsel data in order to decipher reliable signal from experimental noise. Briefly, this analysis (1) removes all mutdiffsel values for variants not present in the starting library of the corresponding replicate, and (2) removes mutdiffsel values that are not of the same sign (positive or negative) across replicates. This analysis is performed by CodonsToAAs.py, FilteringStep1.py, and FilteringStep2.py, which can be found in ./diffsel_data_processing/Filtering

Details about input/output/code are in each sub-directory

### 4-1. Generating logo plots for filtered, averaged diffsel data

This analysis generates sequence logo plots from the filtered diffsel files from Step 3, averaged across biological triplicate. This analysis is performed by ./diffsel_data_processing/Logoplots/SeqLogo.ipynb and primarily uses [dms2_logoplot](https://jbloomlab.github.io/dms_tools2/dms2_logoplot.html)

This analysis requires the following input data

 * Filtered, averaged diffsel files, which can be found in ./diffsel_data_processing/Logoplots/FilteredDiffsel

 * Overlay files, which can be found in ./diffsel_data_processing/Logoplots/overlays

This analysis generates the following output data

 * Sequence logo plots for each selection vs. mock condition

### 4-2. Generating logo plots for unfiltered, unaveraged diffsel data

This analysis generates sequence logo plots from the unfiltered, unaveraged diffsel files from Step 2. This analysis is performed by ./diffsel_data_processing/UnfilteredUnaveragedLogo/UnfilteredUnaveragedLogo.ipynb and primarily uses [dms2_logoplot](https://jbloomlab.github.io/dms_tools2/dms2_logoplot.html)

This analysis requires the following input data

 * Unfiltered, unaveraged diffsel files, which can be found in ./diffsel_data_processing/UnfilteredUnaveragedLogo/diffsel

 * Overlay files, which can be found in ./diffsel_data_processing/UnfilteredUnaveragedLogo/overlays

This analysis generates the following output data

 * Sequence logo plots for each replicate for each selection vs. mock condition
