## Supplementary figures and images for "The Host Cell’s Endoplasmic Reticulum Proteostasis Network Profoundly Shapes the Protein Sequence Space Accessible to HIV Envelope"

### ATF6-replicate1_diffsel.pdf

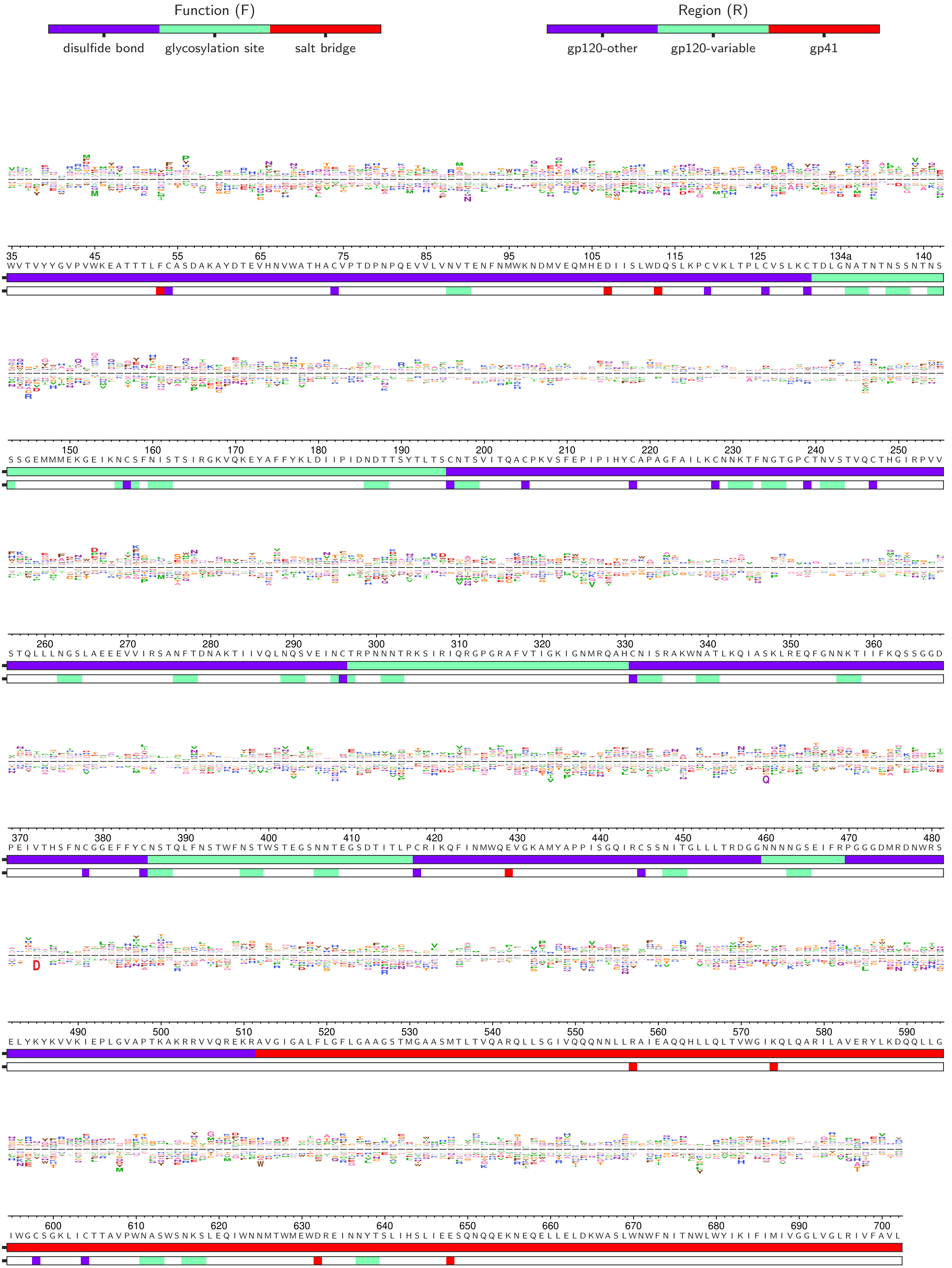

### ATF6-replicate2_diffsel.pdf

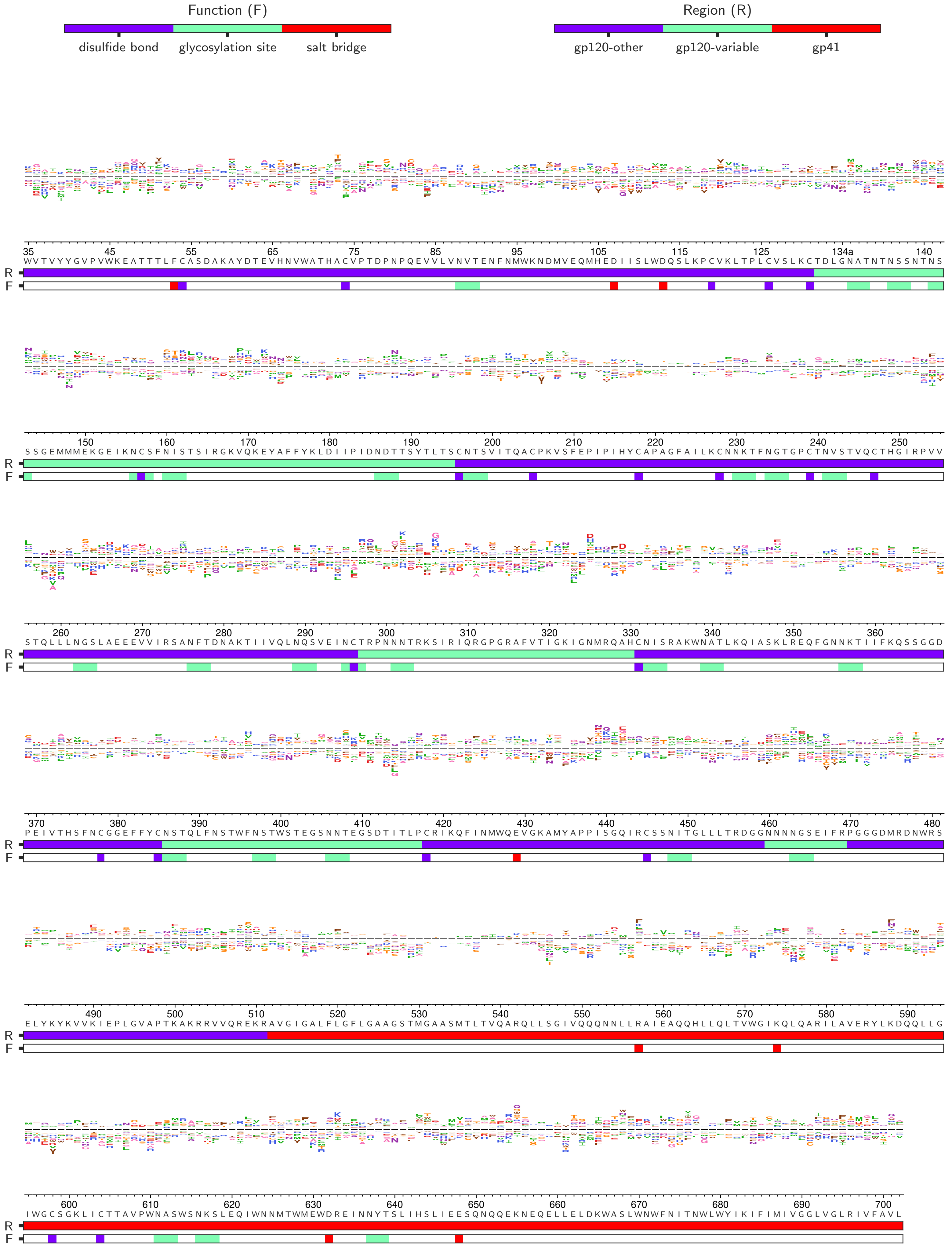

### ATF6-replicate3_diffsel.pdf

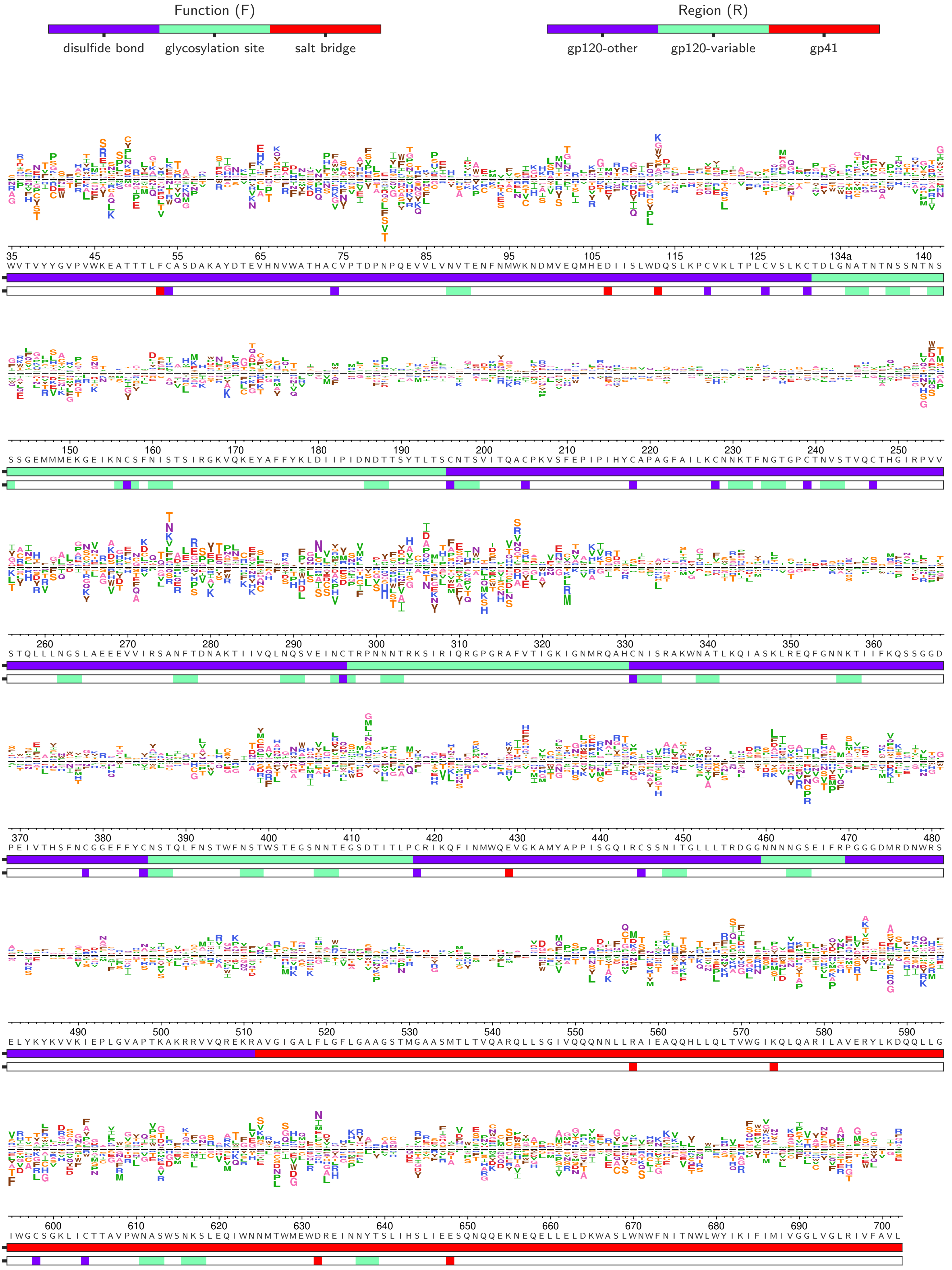

### ATF6_diffsel.pdf

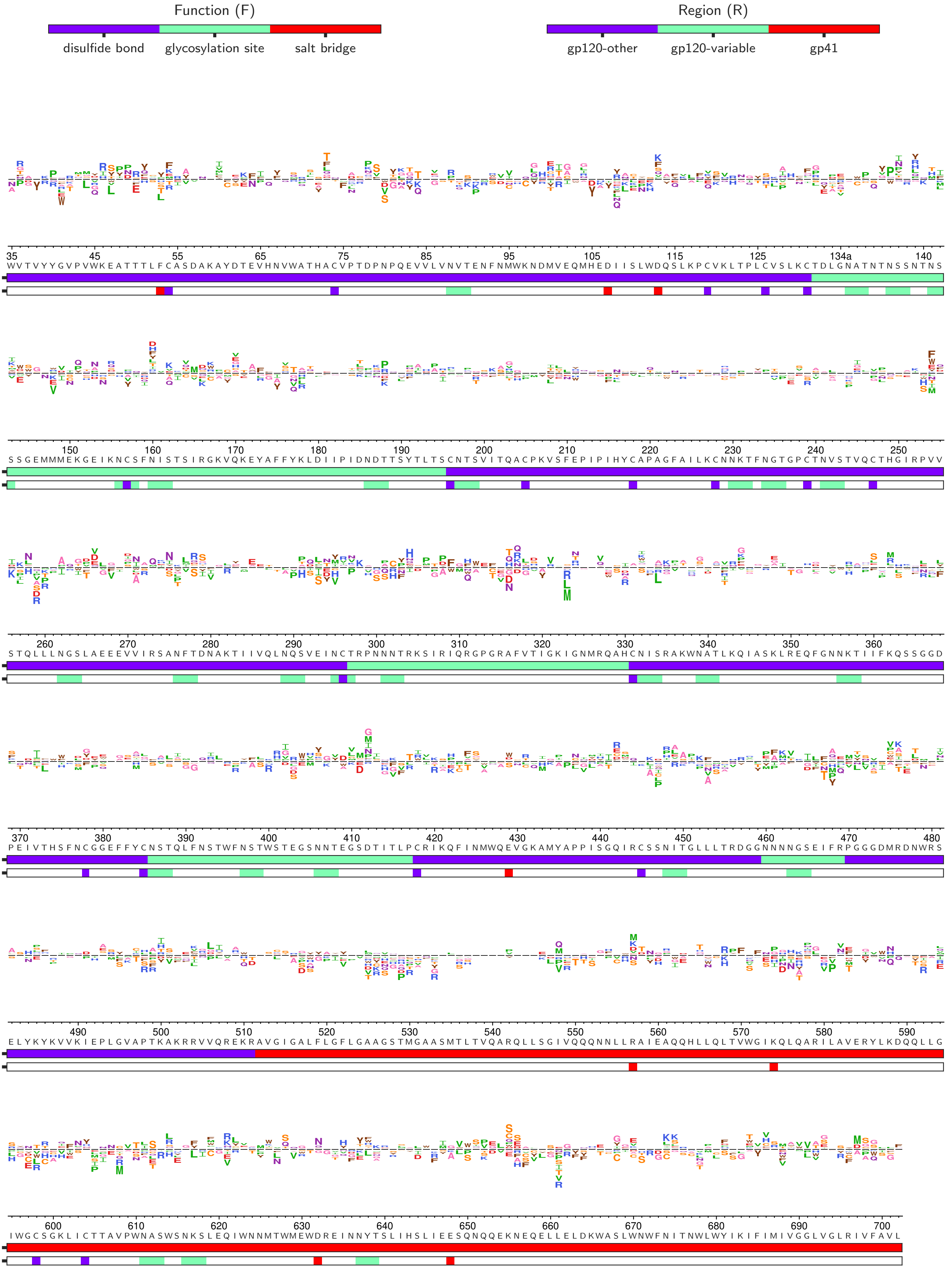

### summary_ATF6-absolutesitediffselcorr.pdf

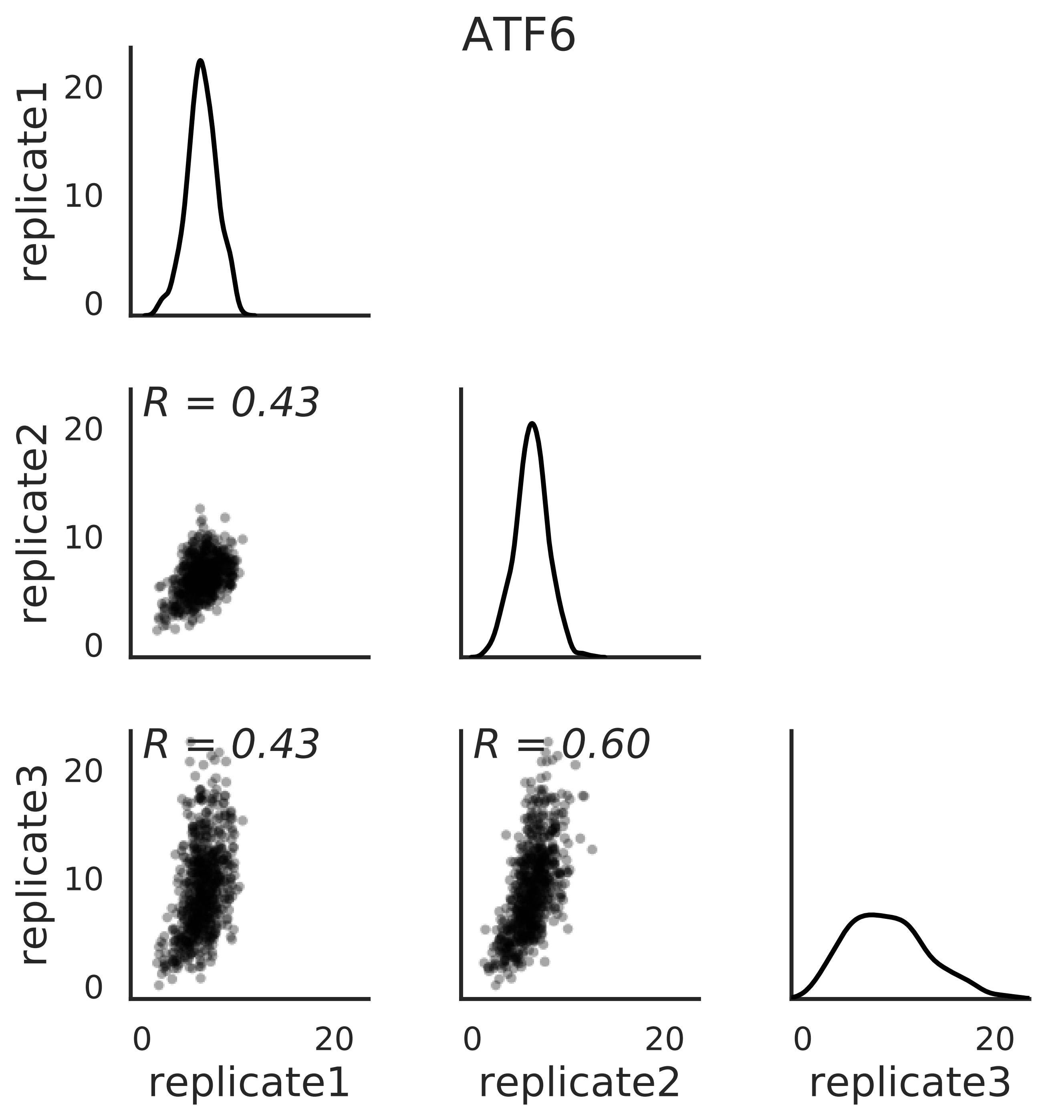

### summary_ATF6-maxmutdiffselcorr.pdf

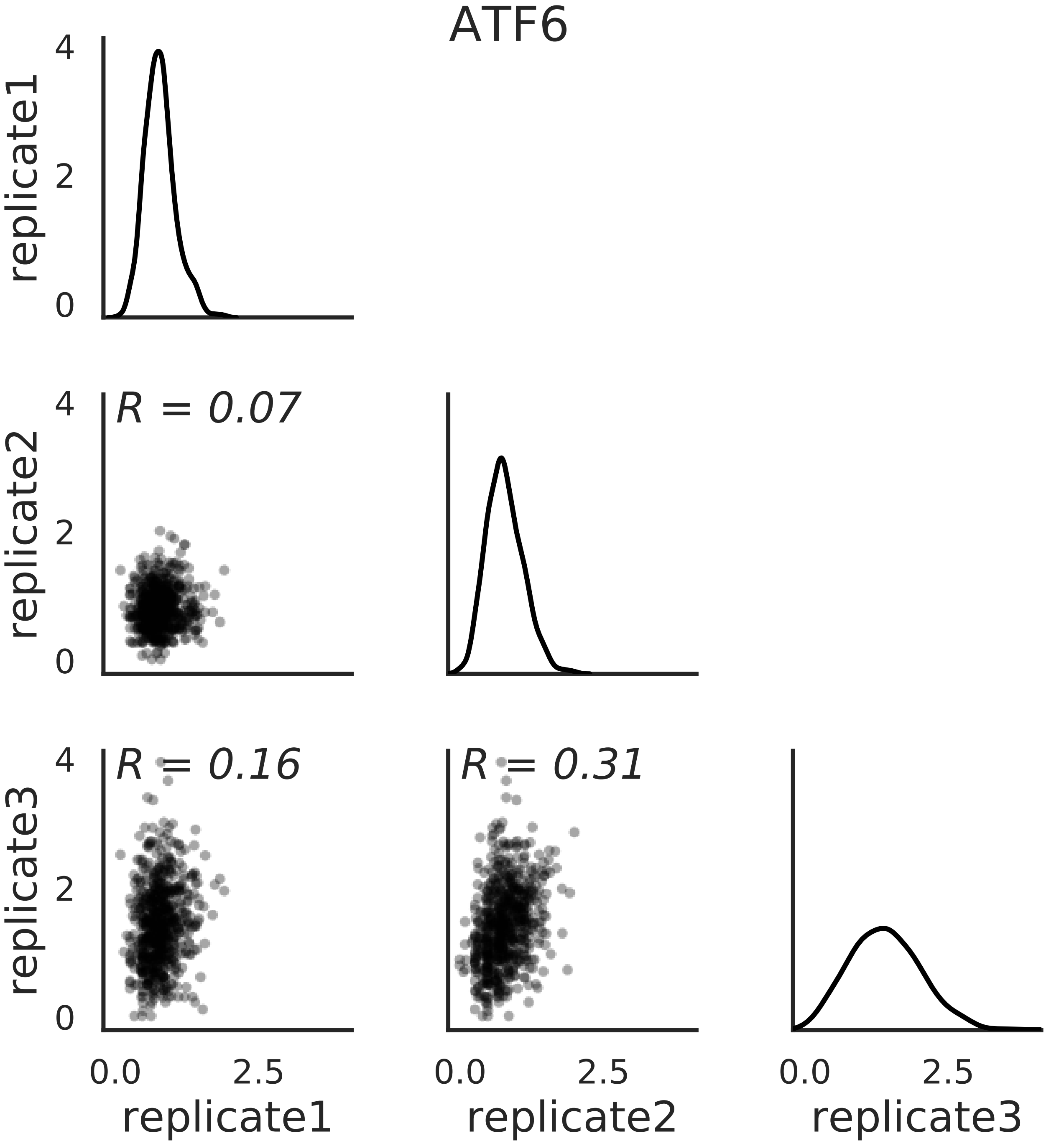

### summary_ATF6-mutdiffselcorr.pdf

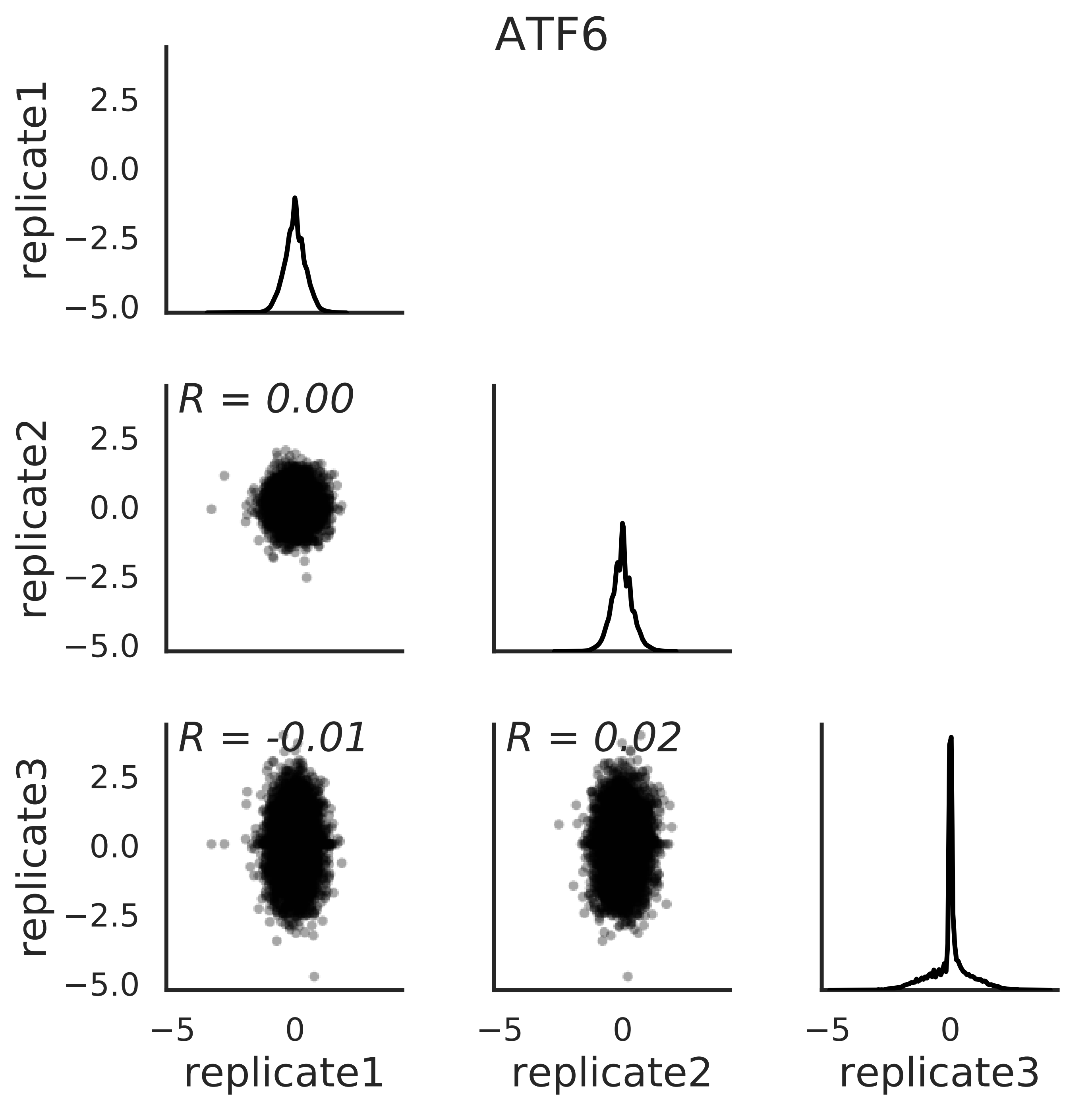

### summary_ATF6-positivesitediffselcorr.pdf

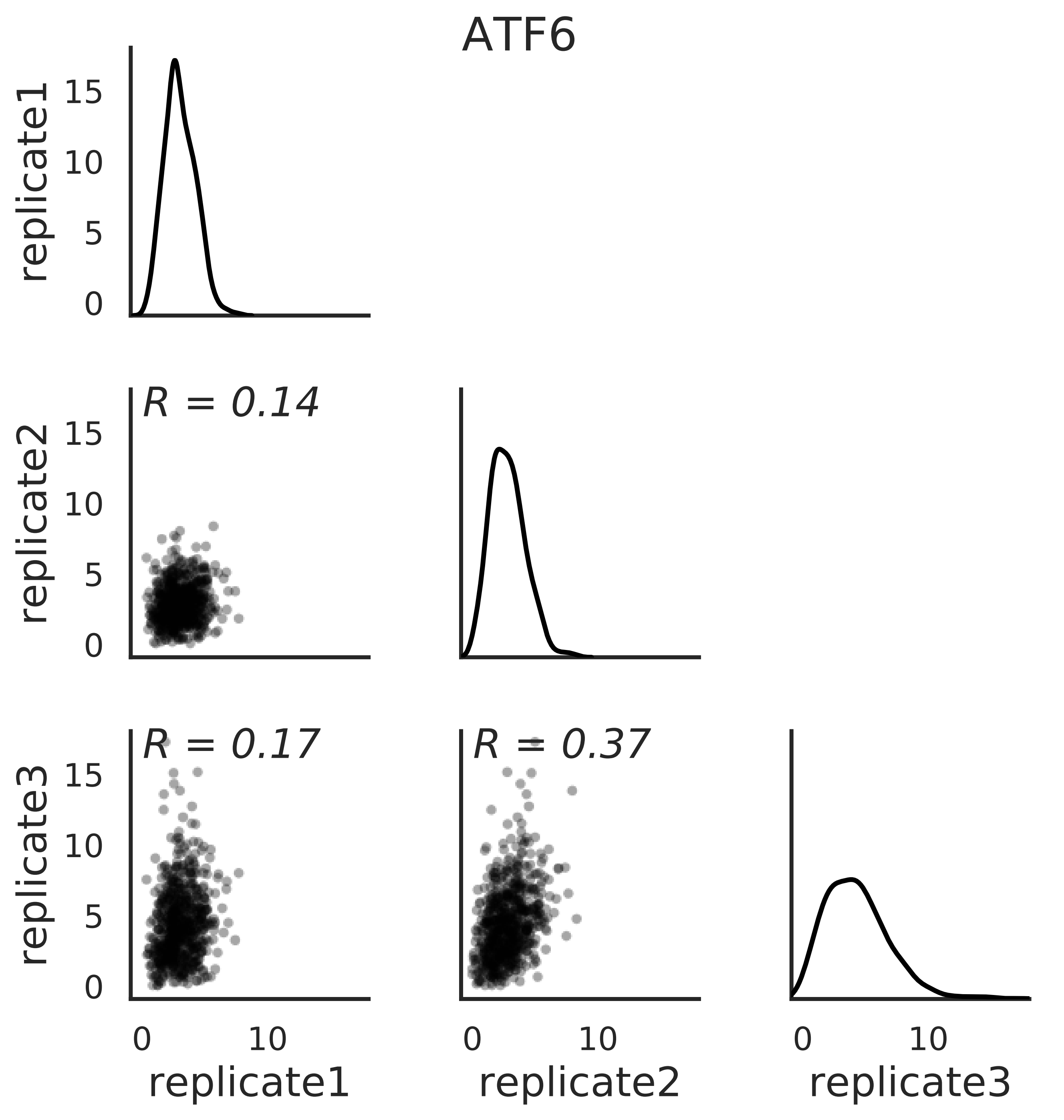

### summary_meanmaxdiffsel.pdf

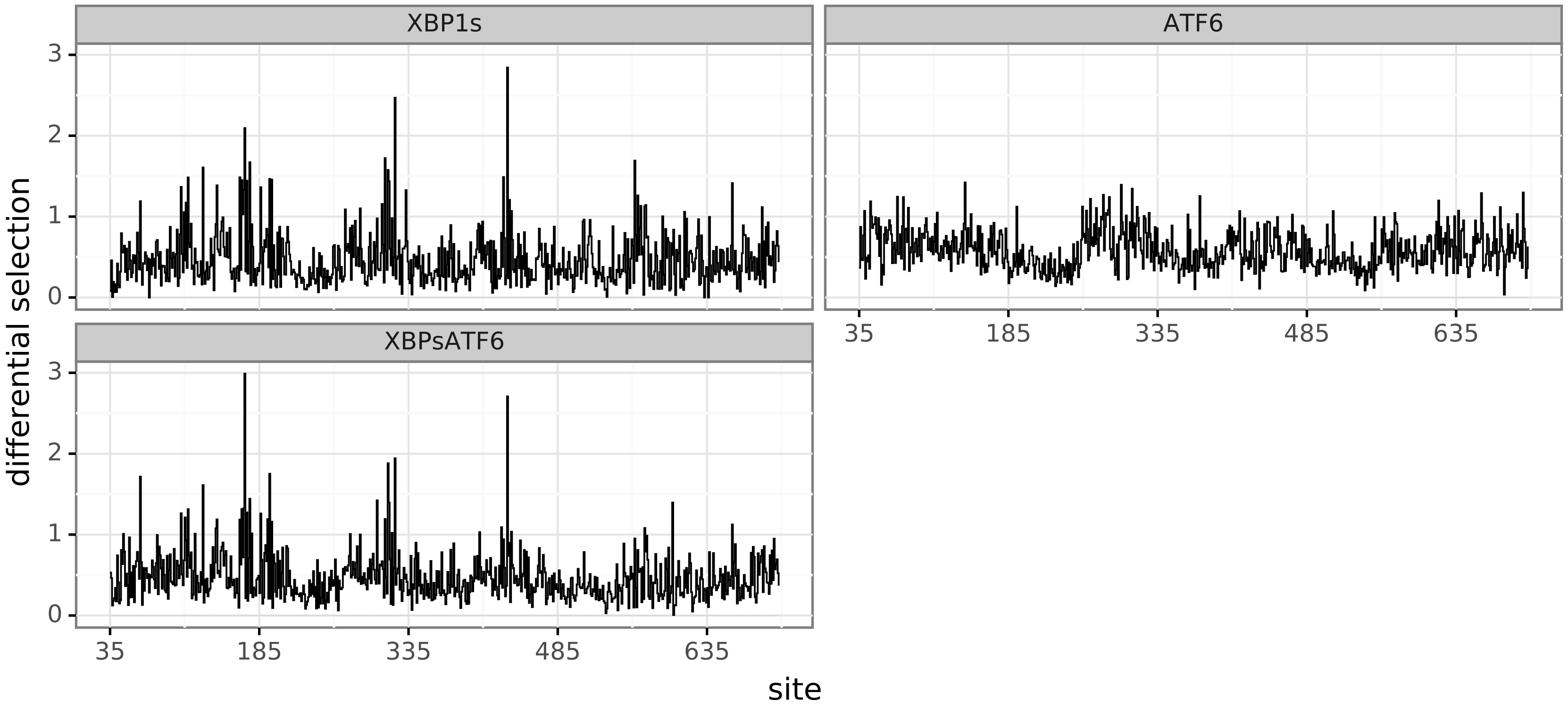

### summary_meanminmaxdiffsel.pdf

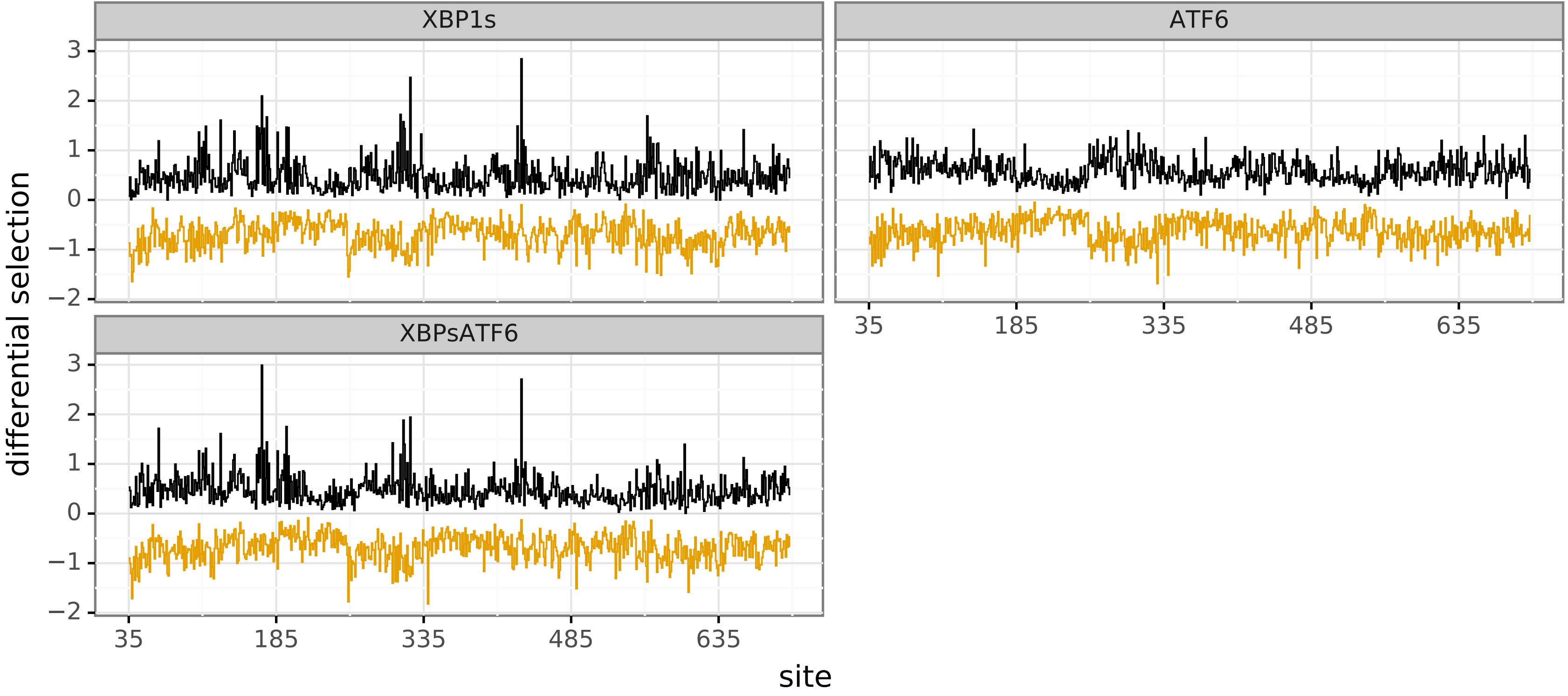

### summary_meanpositivediffsel.pdf

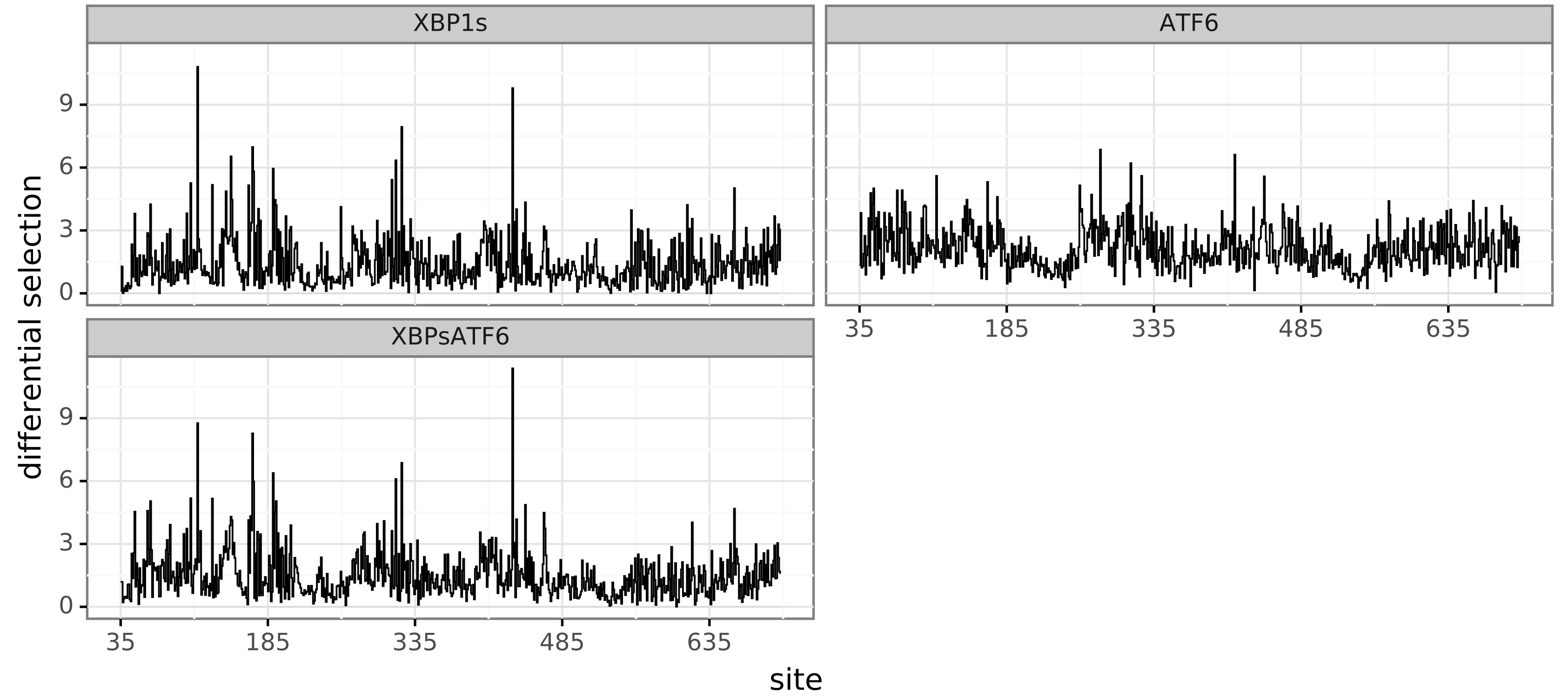

### summary_meantotaldiffsel.pdf

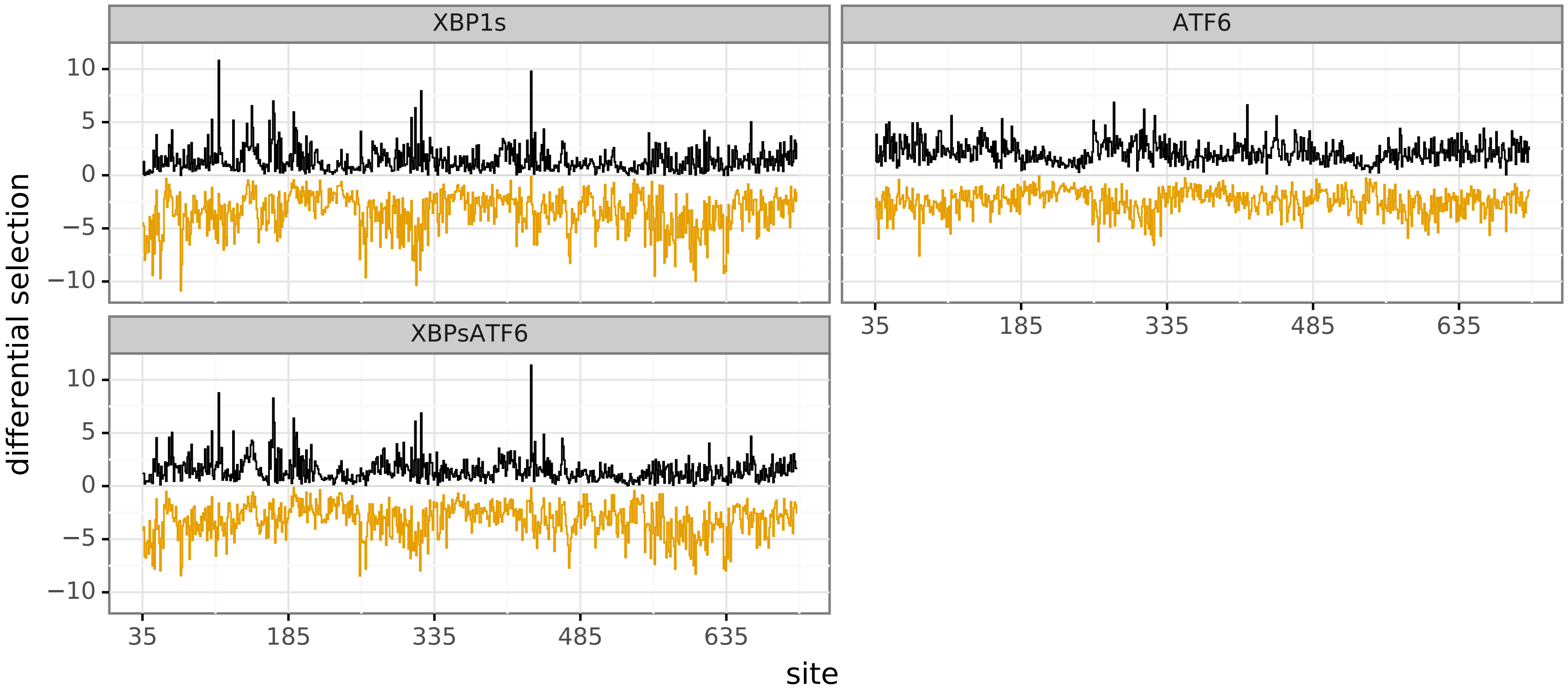

### summary_medianmaxdiffsel.pdf

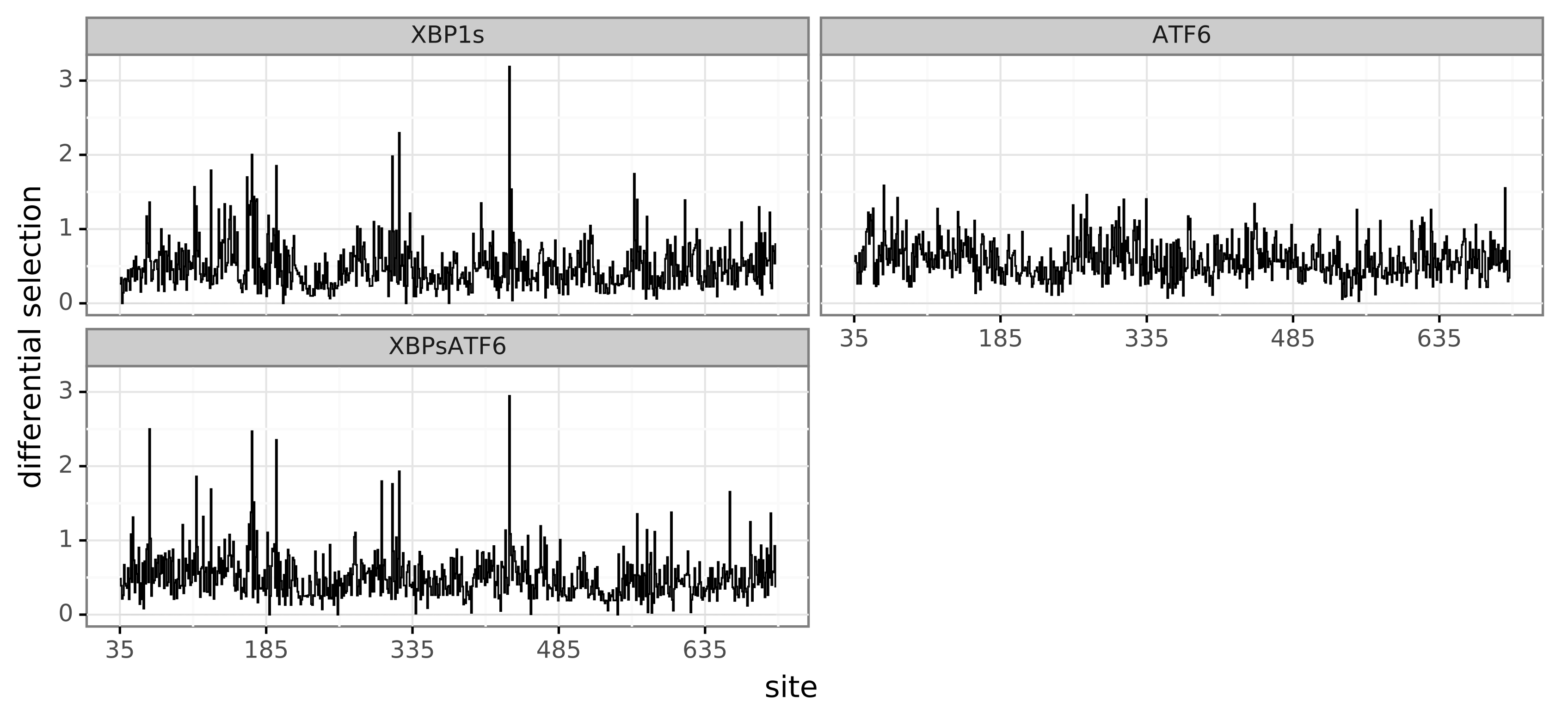

### summary_medianminmaxdiffsel.pdf

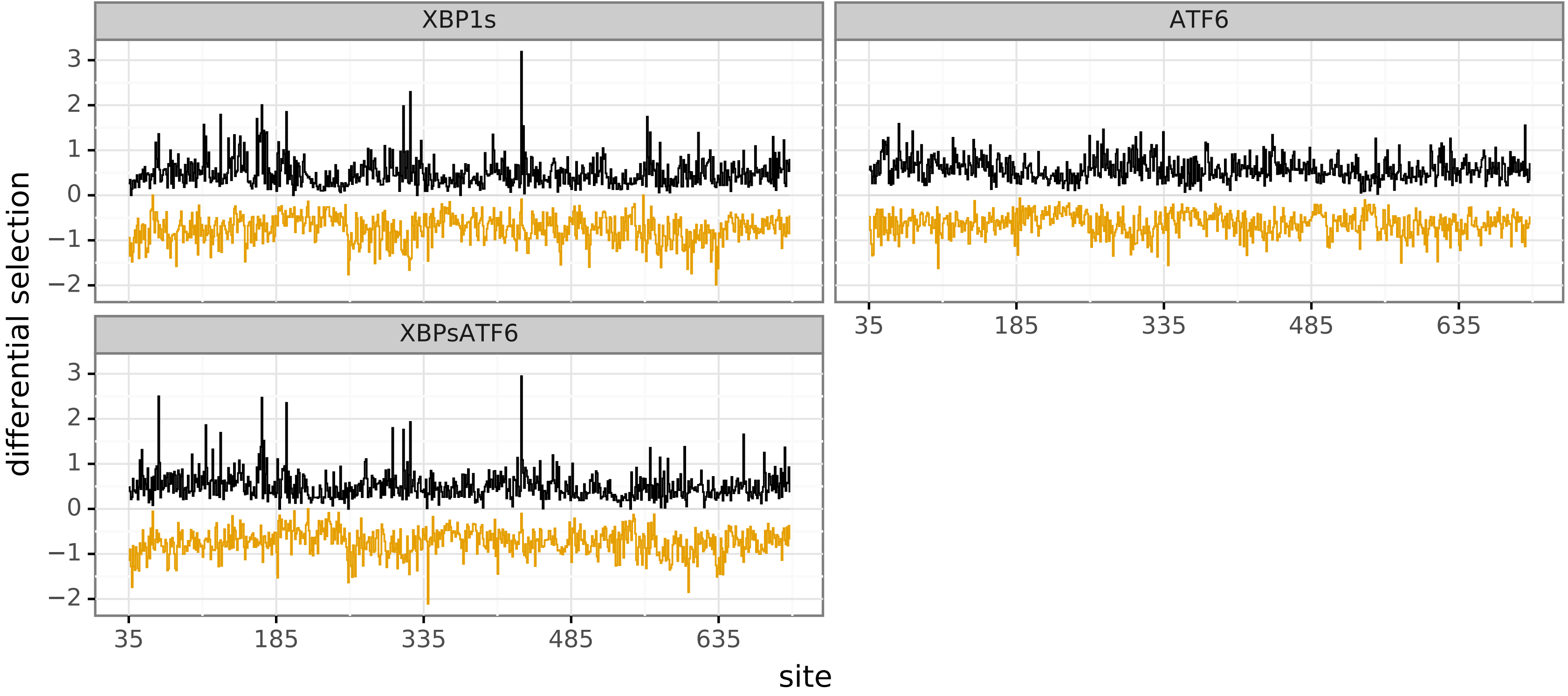

### summary_medianpositivediffsel.pdf

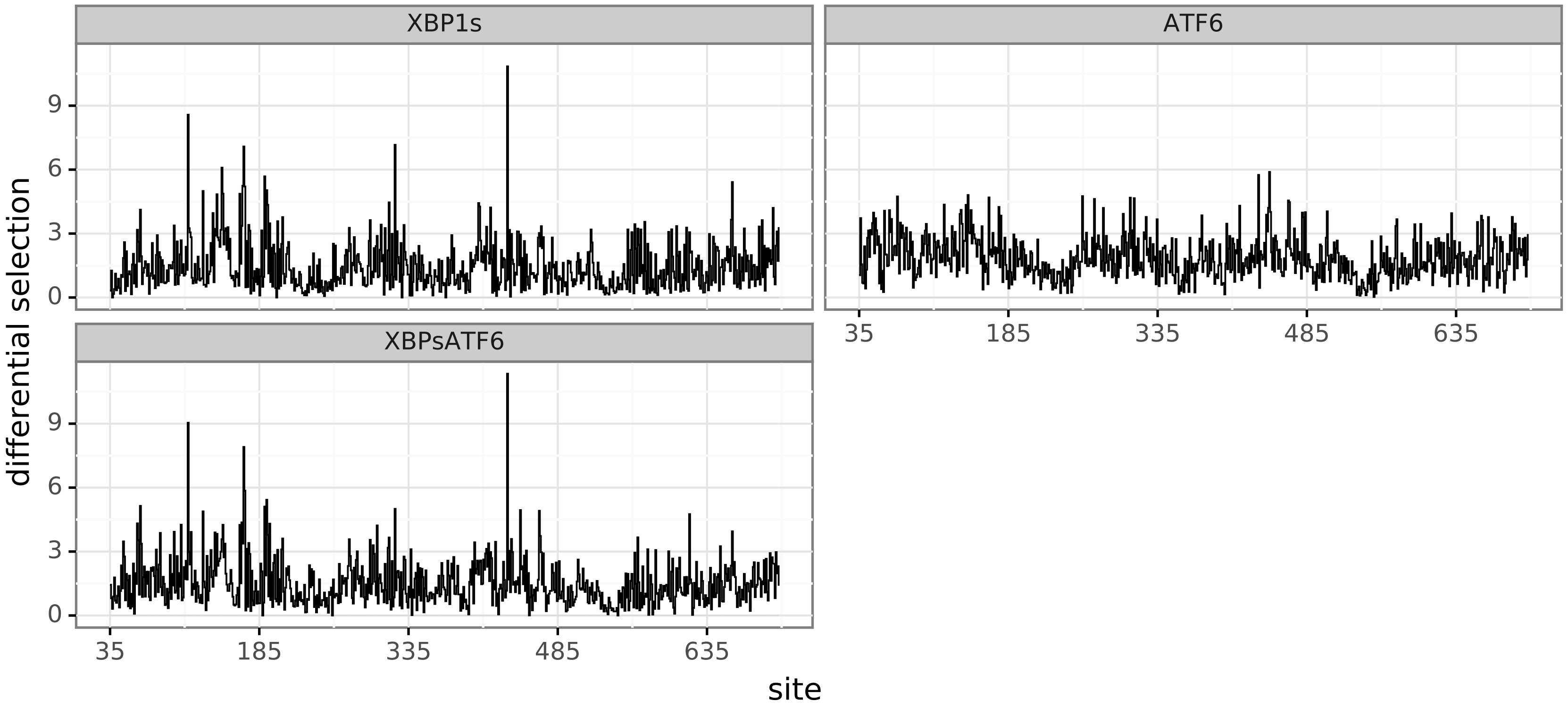

### summary_mediantotaldiffsel.pdf

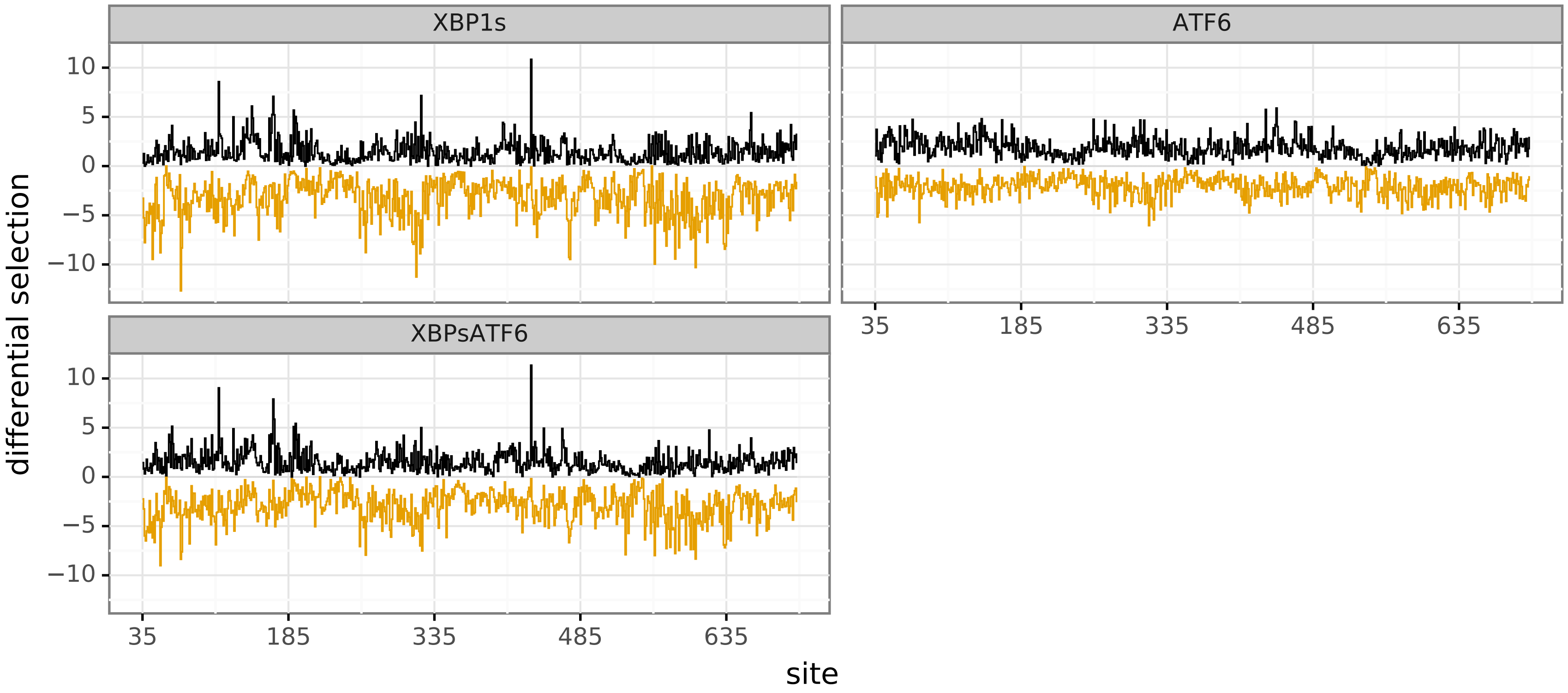

### summary_XBP1s-absolutesitediffselcorr.pdf

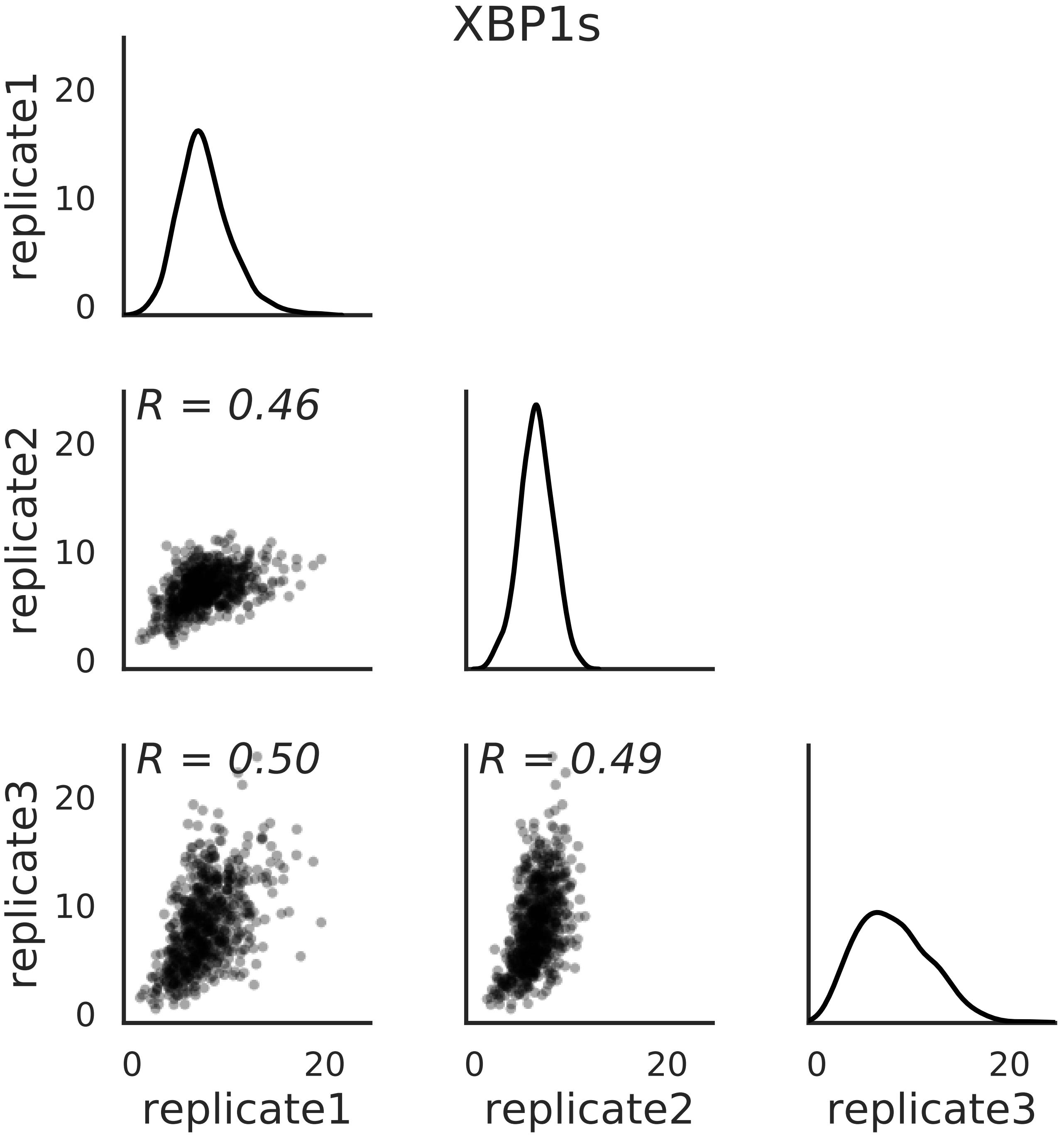

### summary_XBP1s-maxmutdiffselcorr.pdf

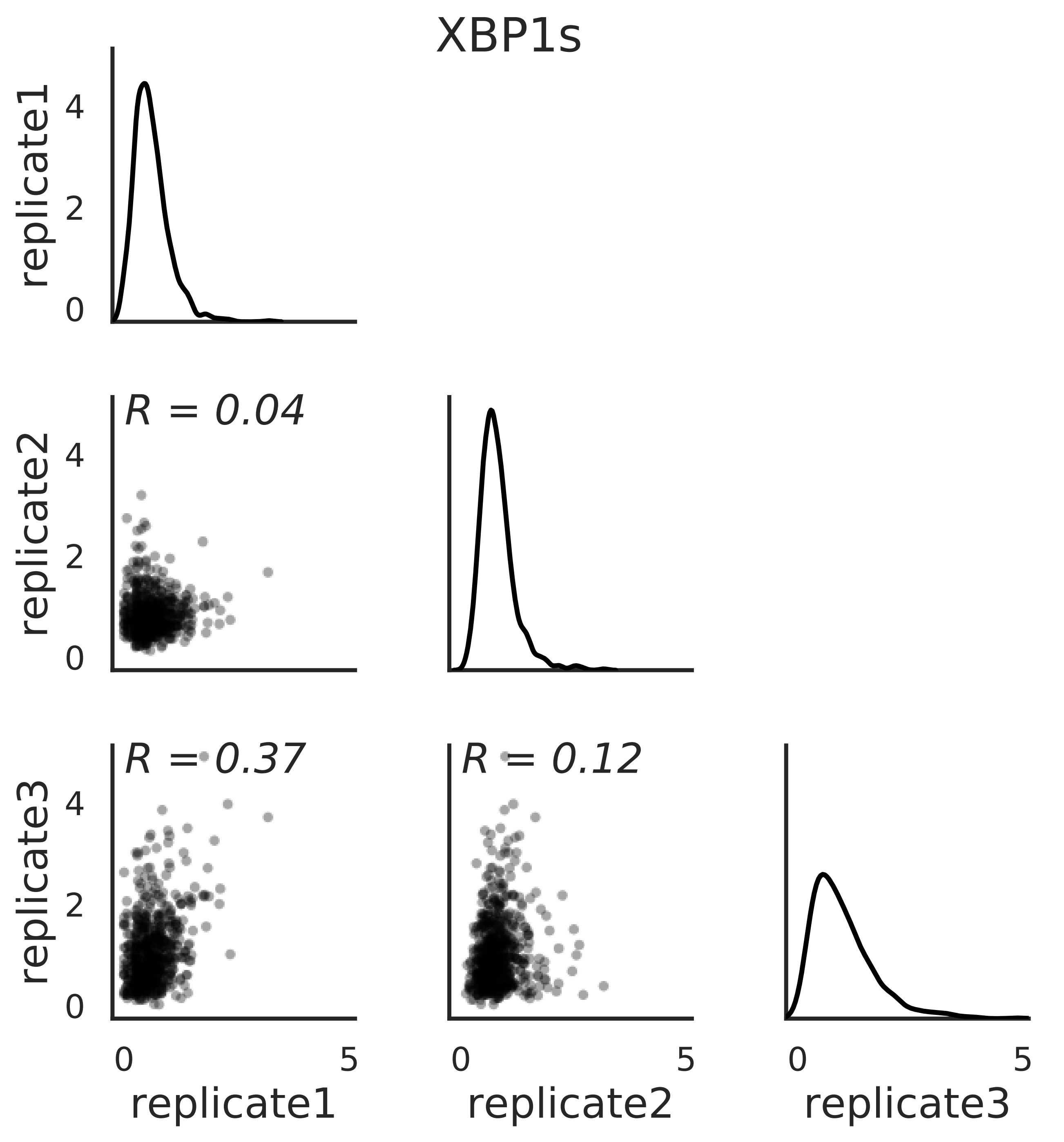

### summary_XBP1s-mutdiffselcorr.pdf

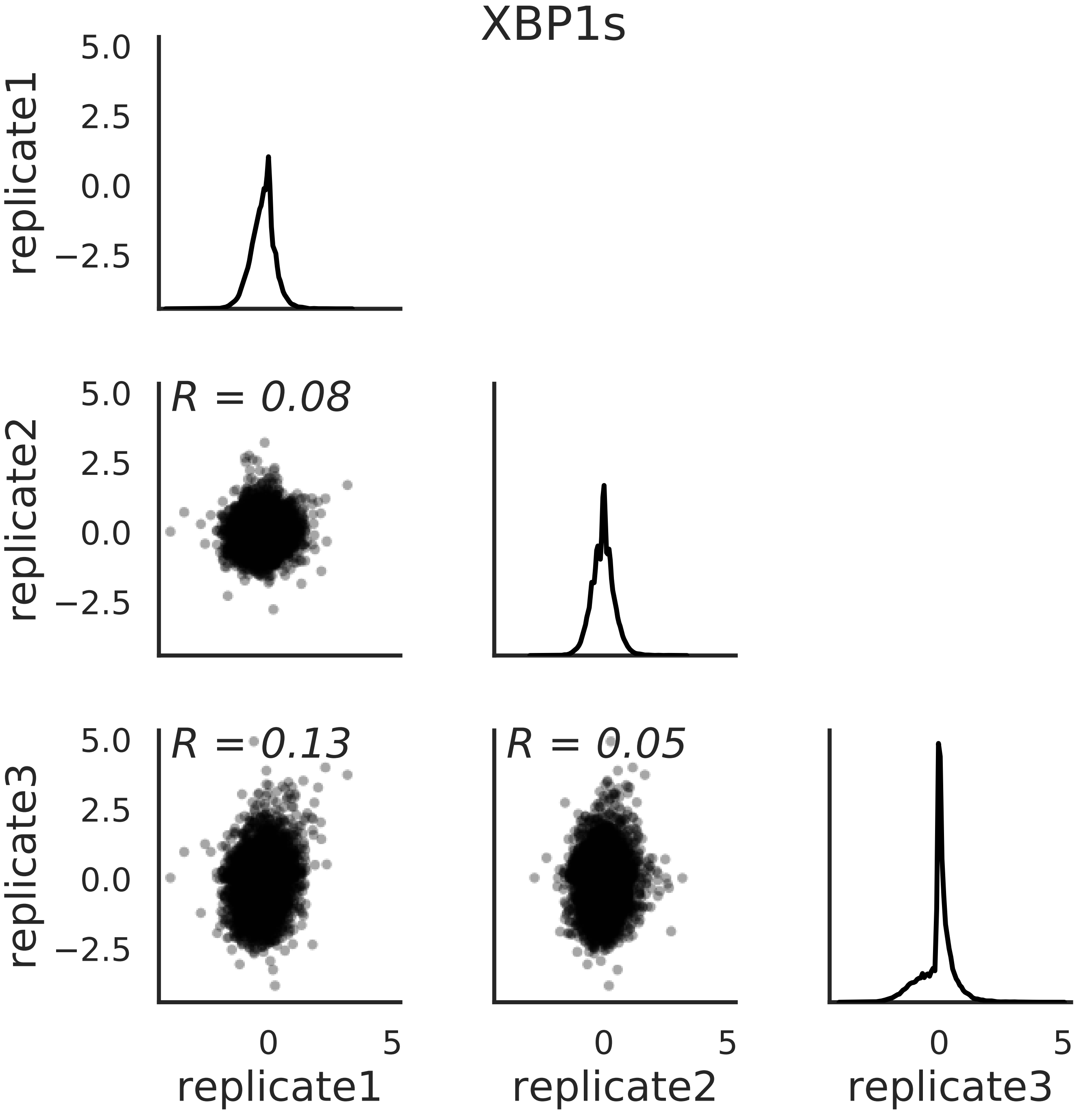

### summary_XBP1s-positivesitediffselcorr.pdf

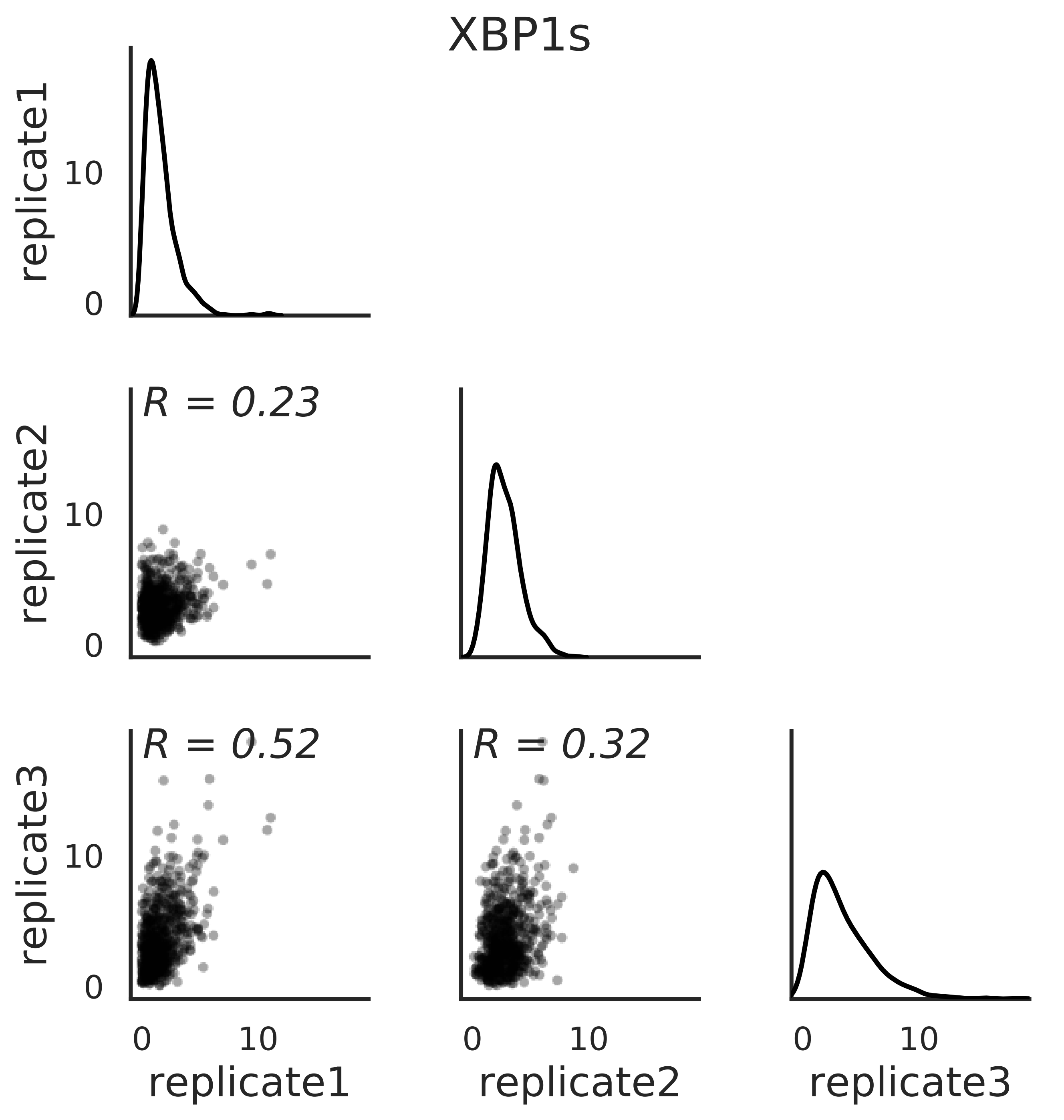

### summary_XBPsATF6-absolutesitediffselcorr.pdf

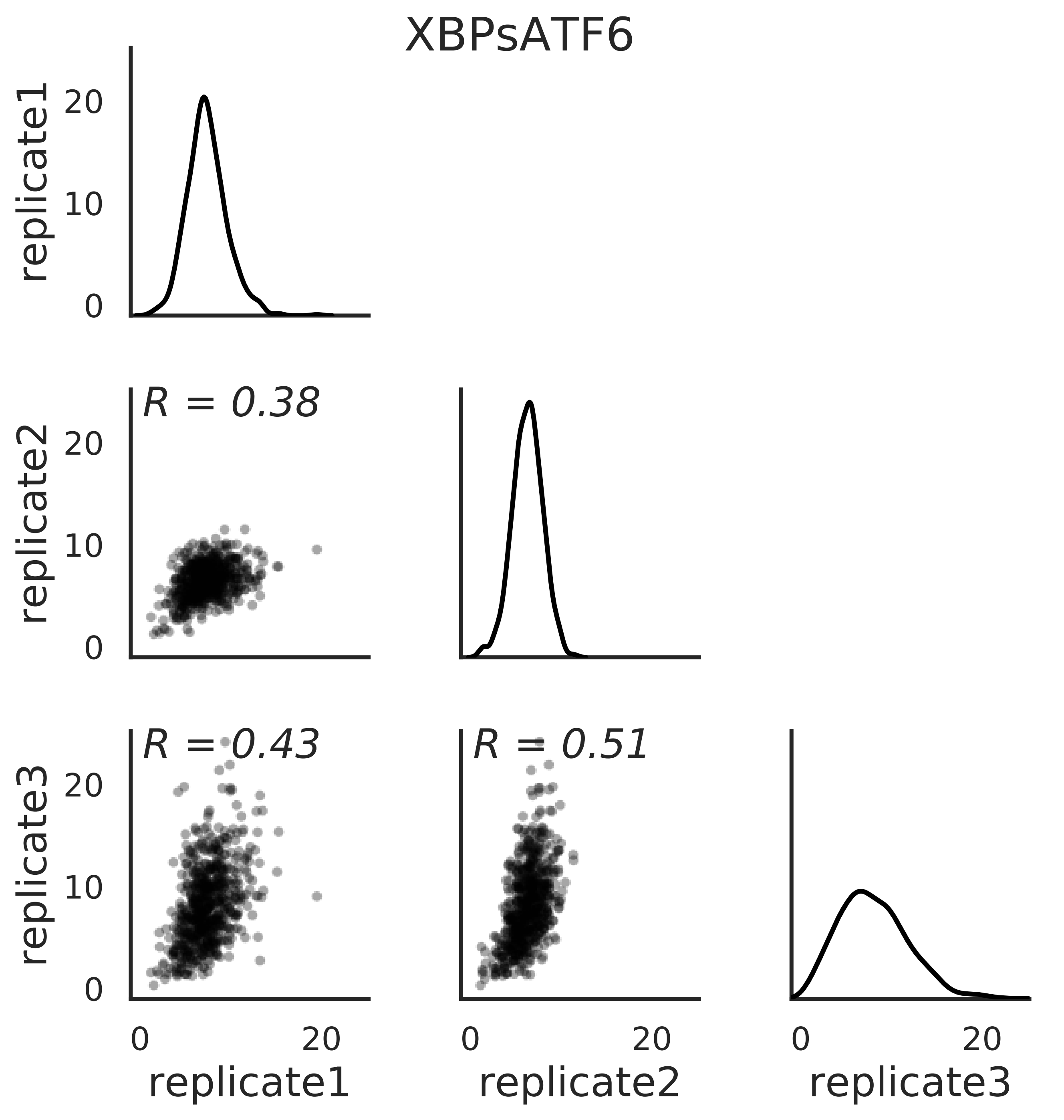

### summary_XBPsATF6-maxmutdiffselcorr.pdf

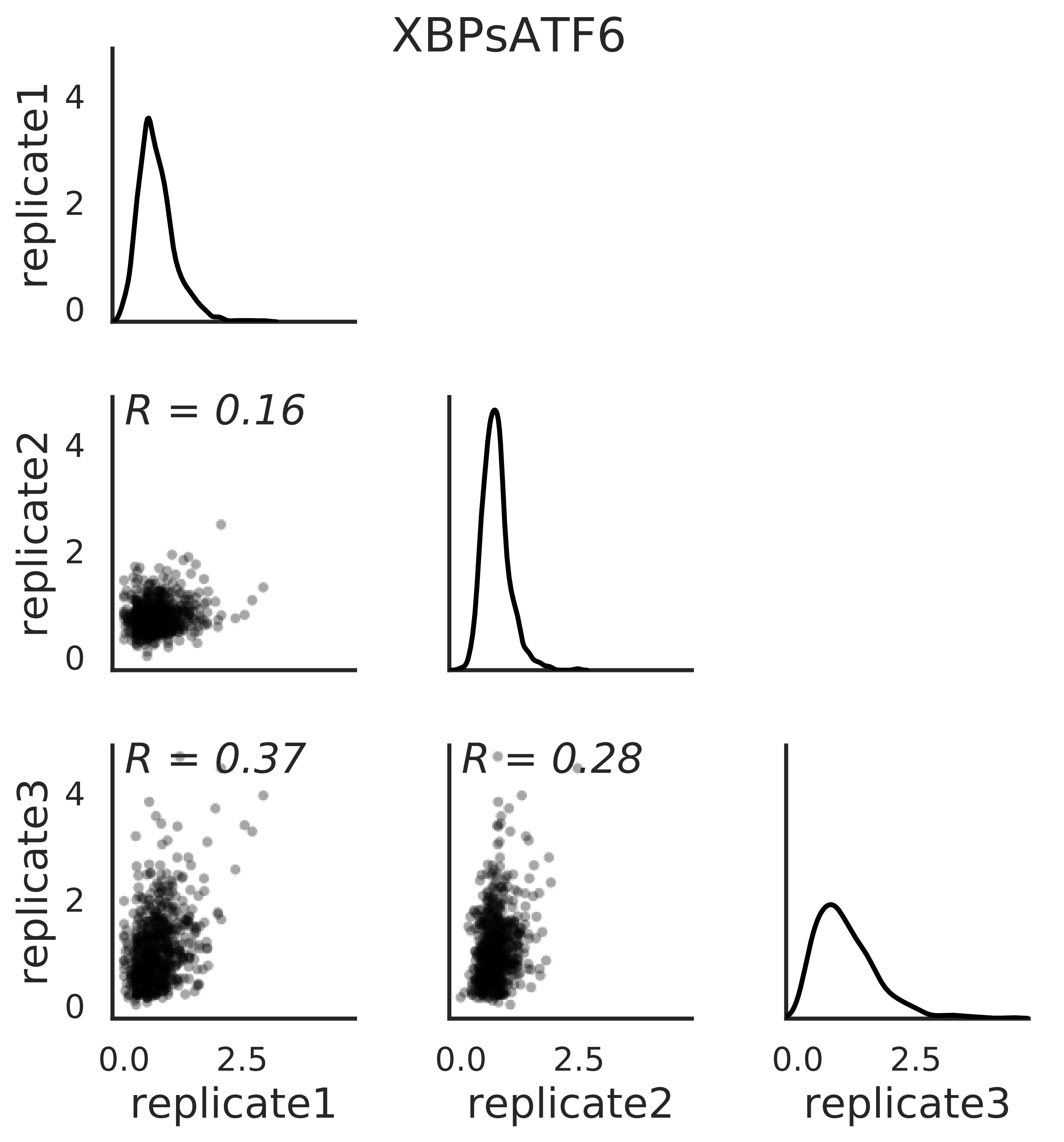

### summary_XBPsATF6-mutdiffselcorr.pdf

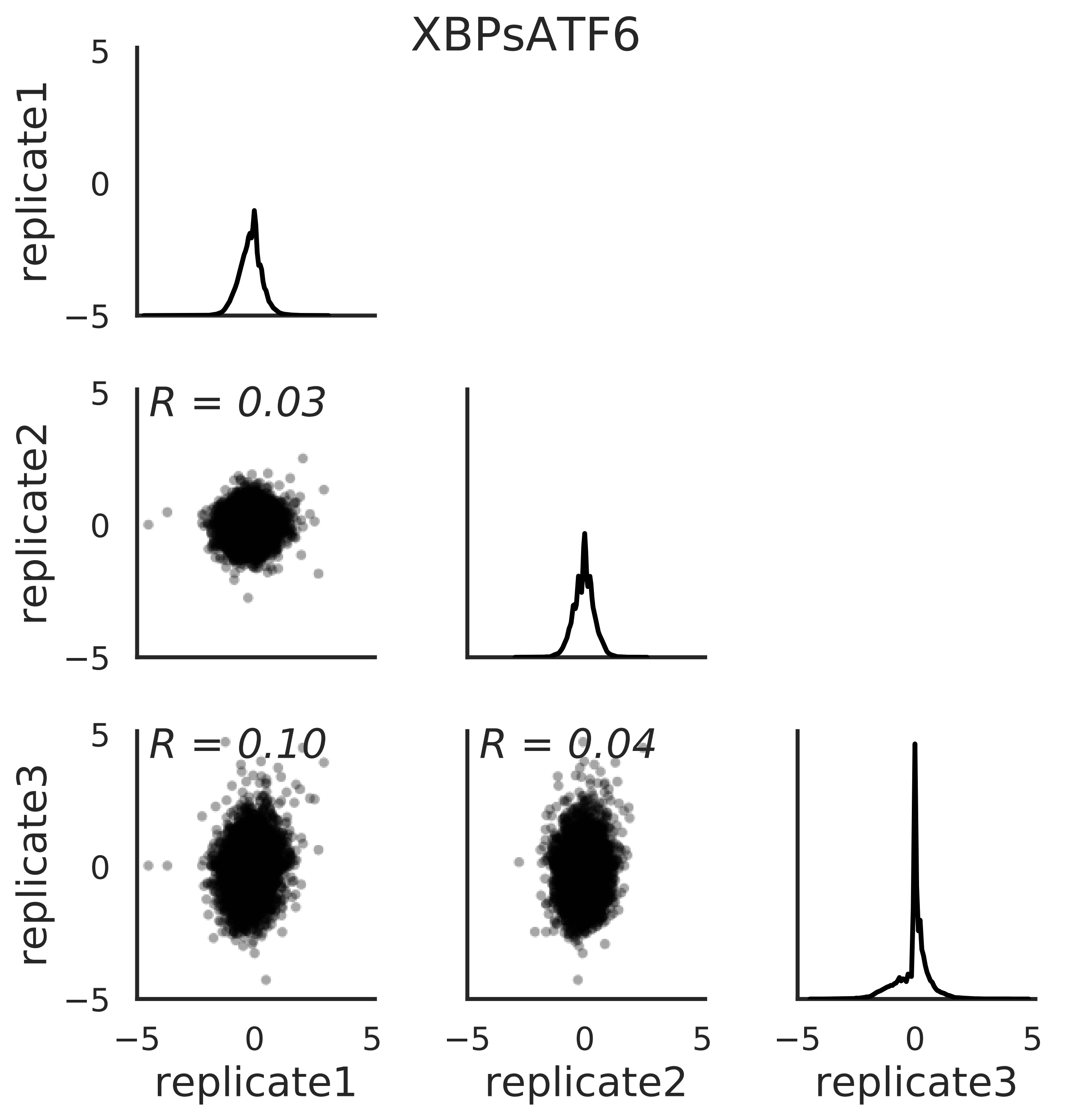

### summary_XBPsATF6-positivesitediffselcorr.pdf

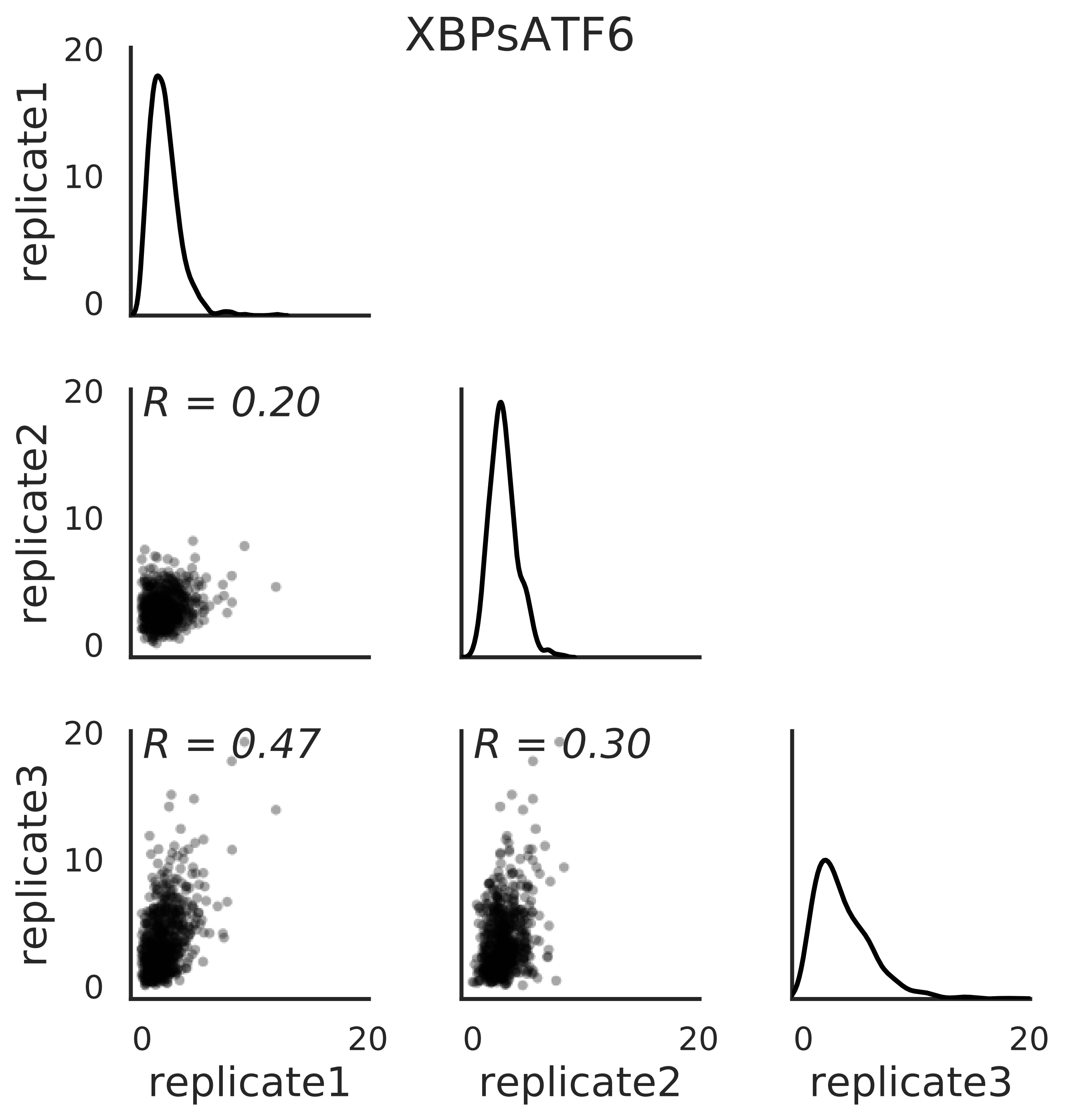

### XBP1s-replicate1_diffsel.pdf

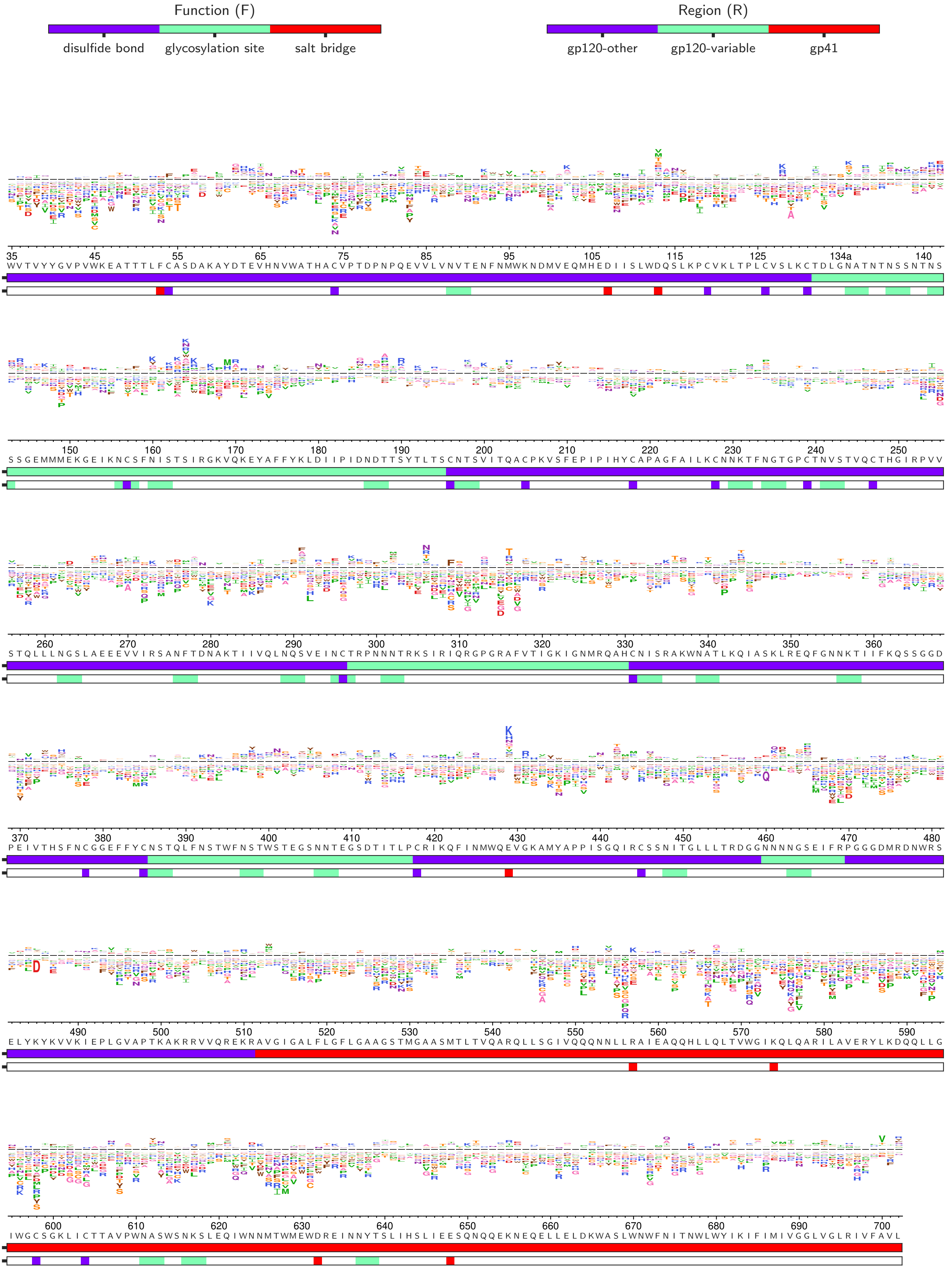

### XBP1s-replicate2_diffsel.pdf

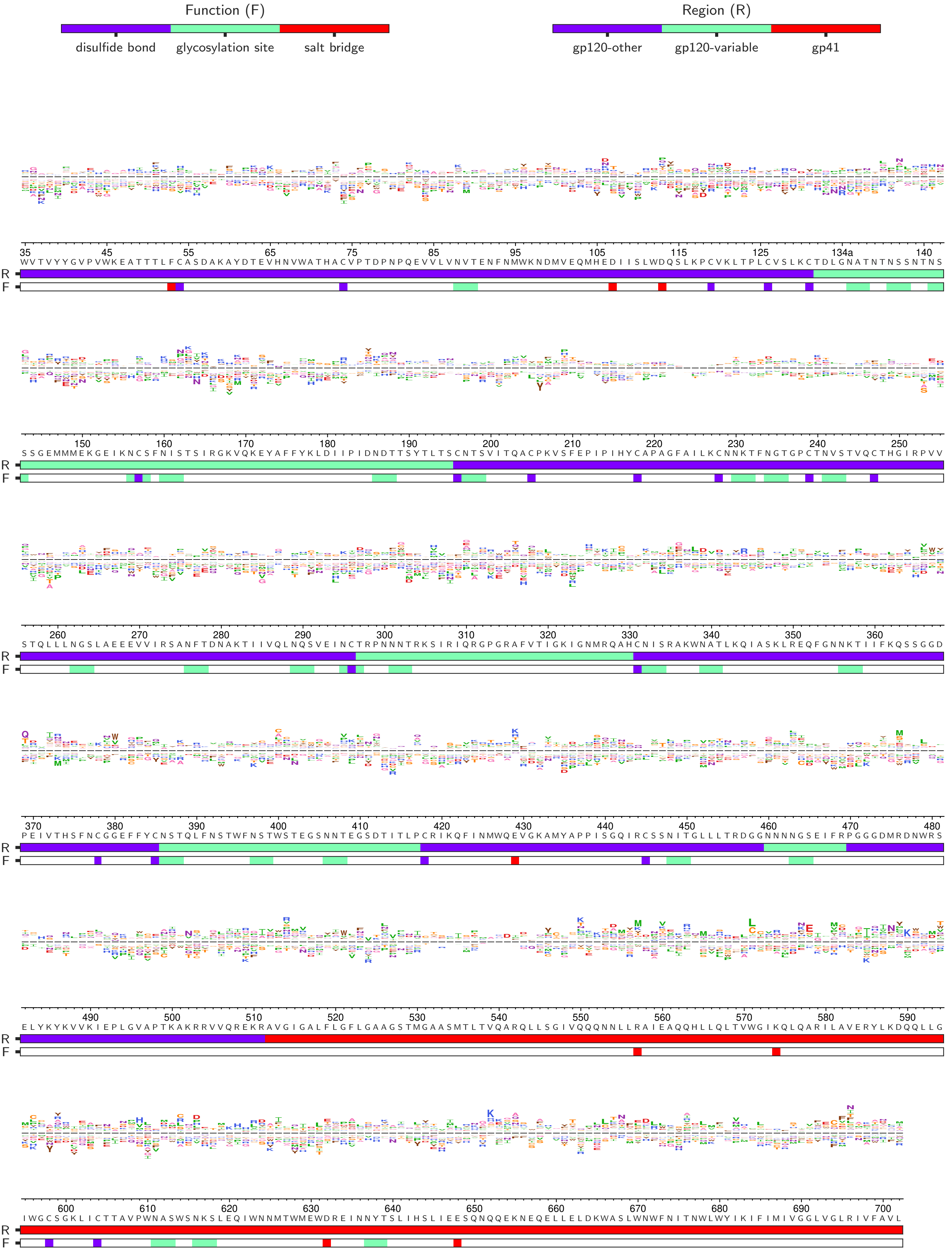

### XBP1s-replicate3_diffsel.pdf

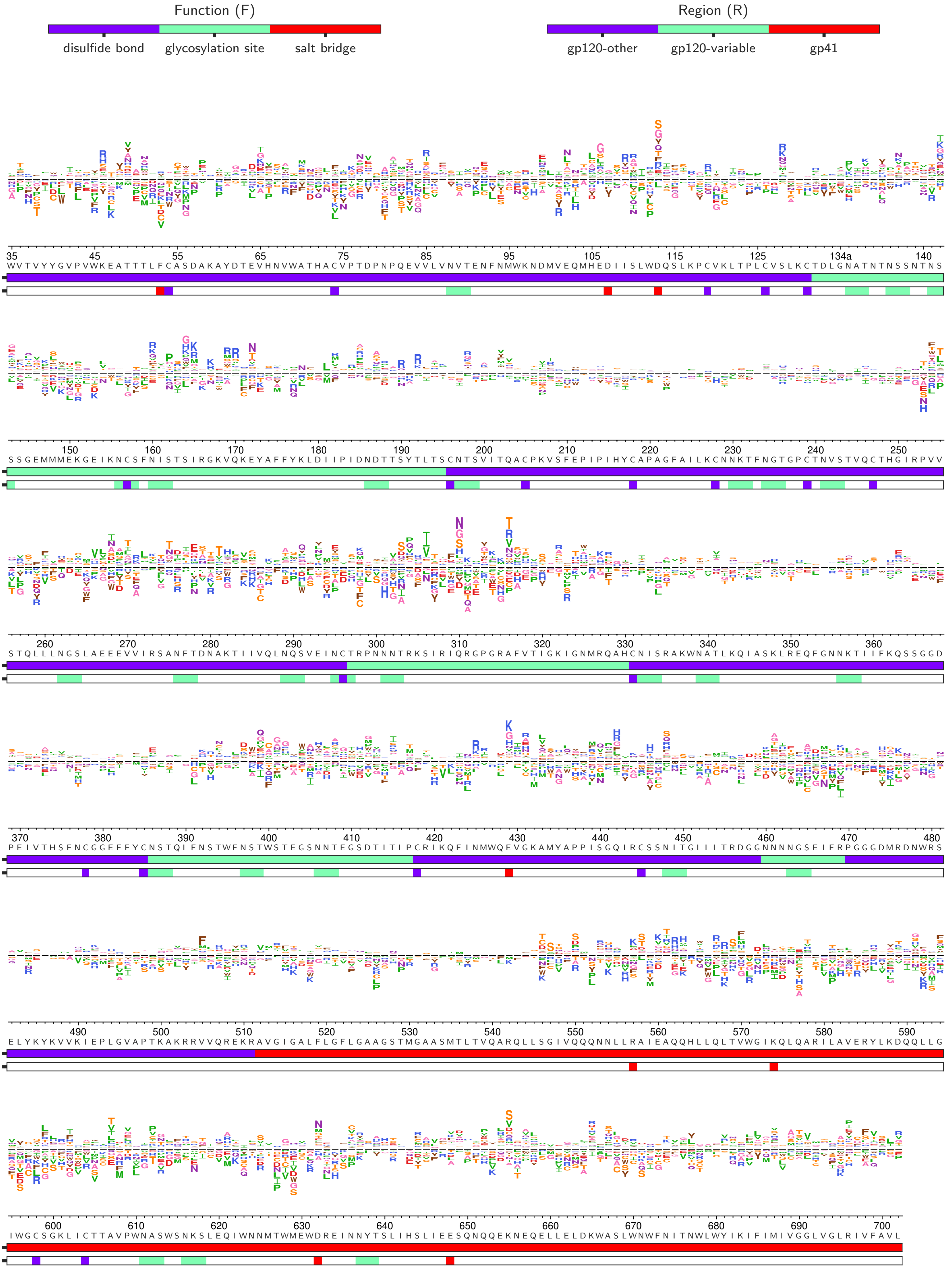

### XBP1s_diffsel.pdf

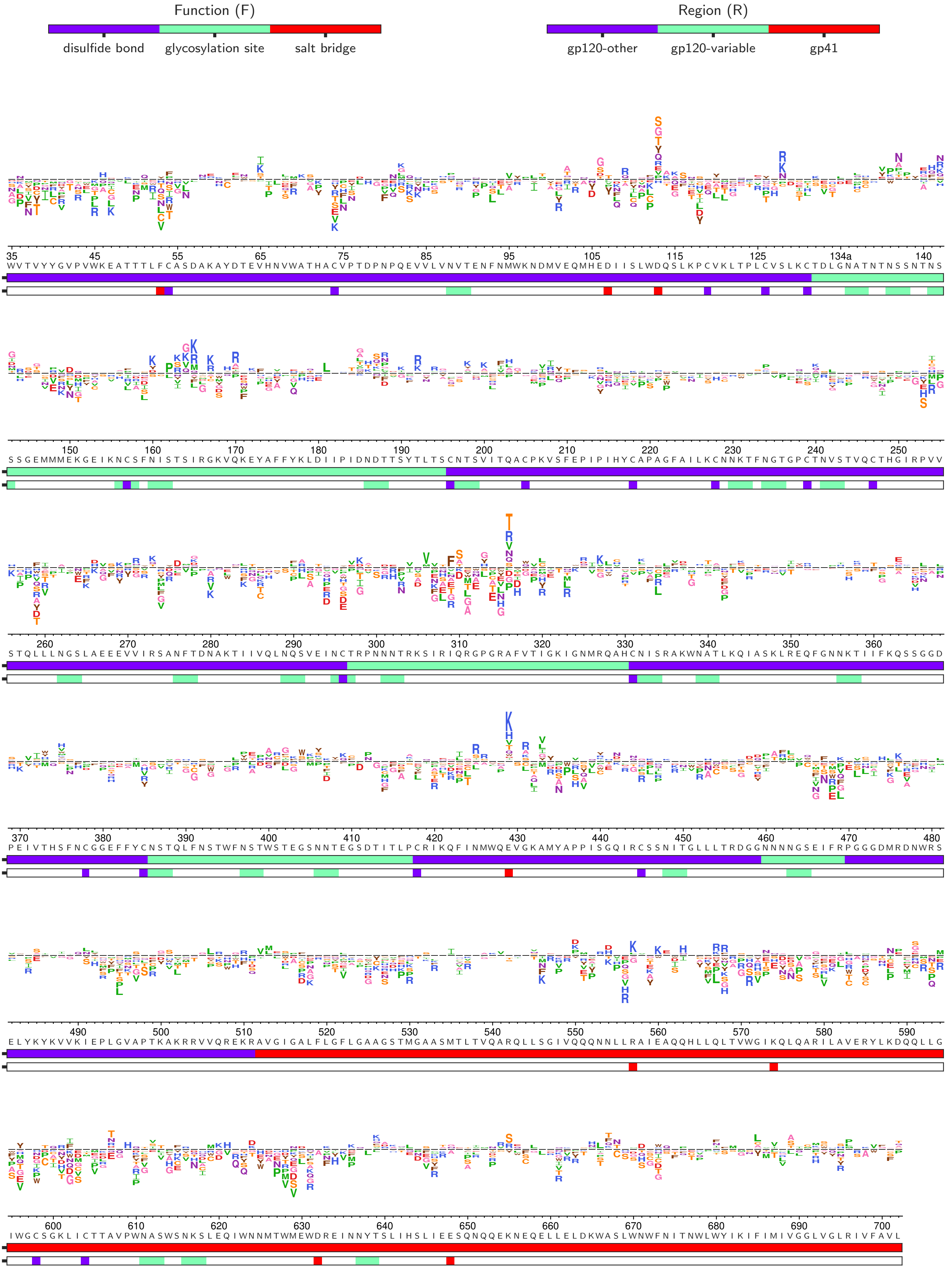

### XBP1sATF6-replicate1_diffsel.pdf

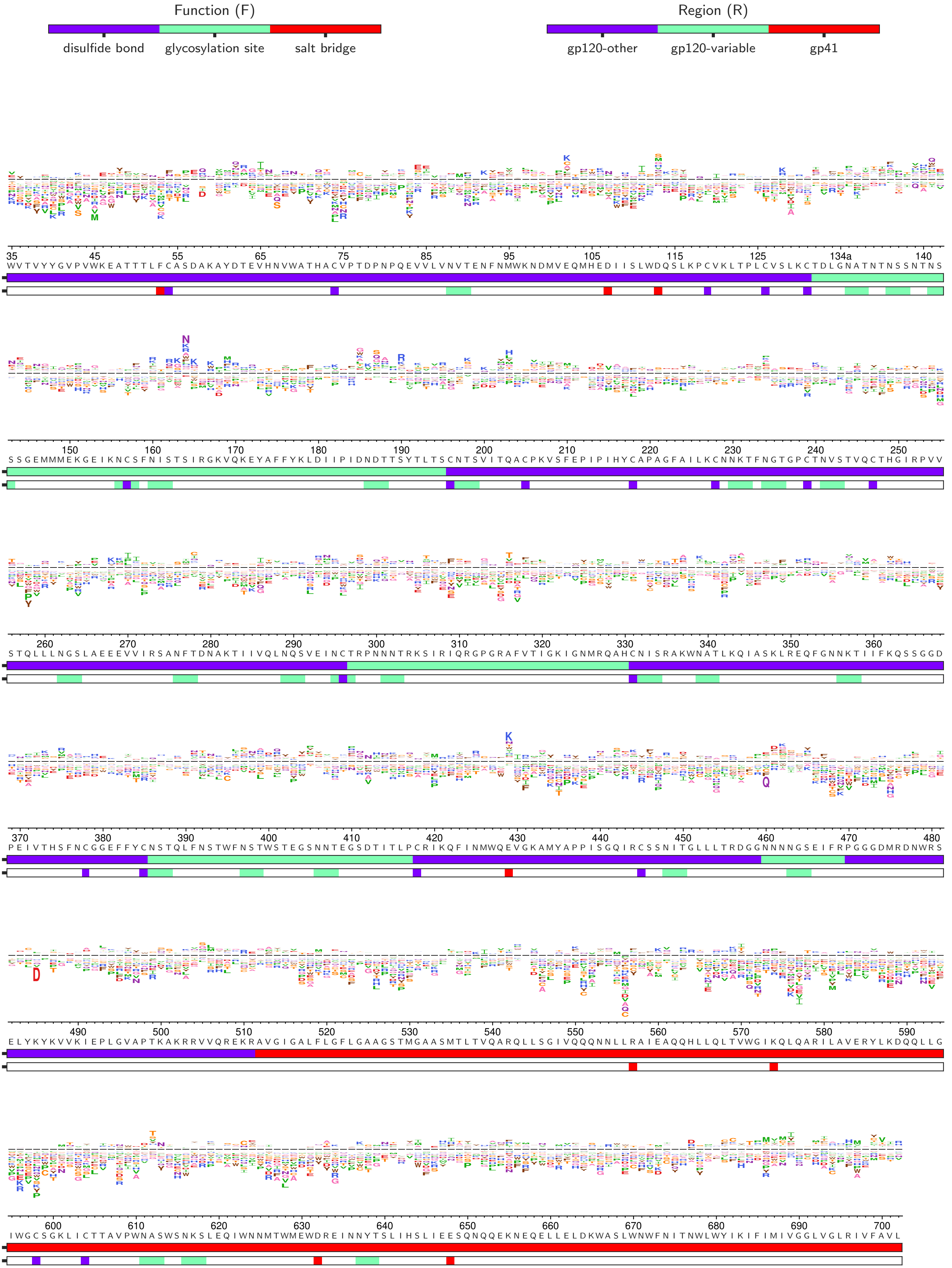

### XBP1sATF6-replicate2_diffsel.pdf

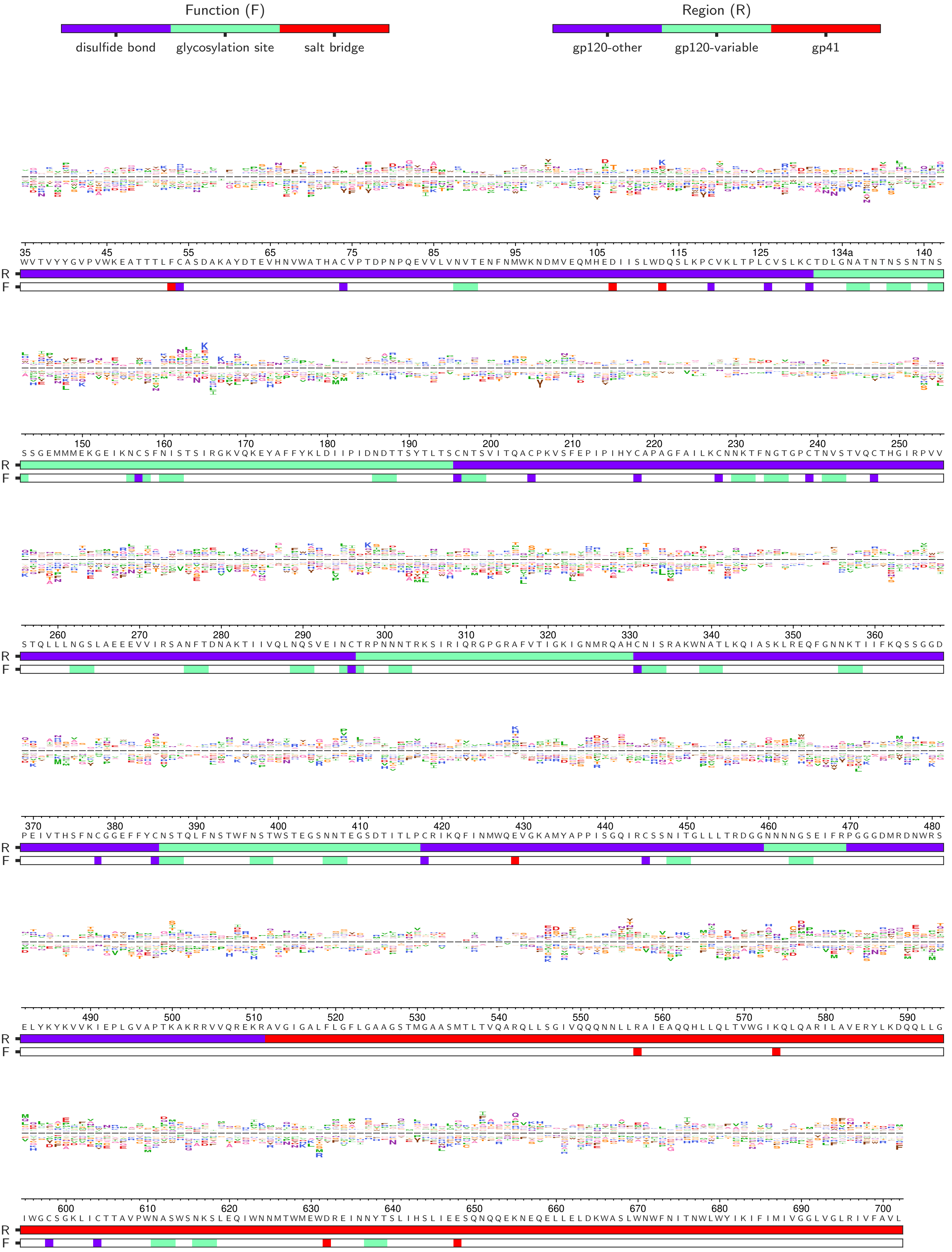

### XBP1sATF6-replicate3_diffsel.pdf

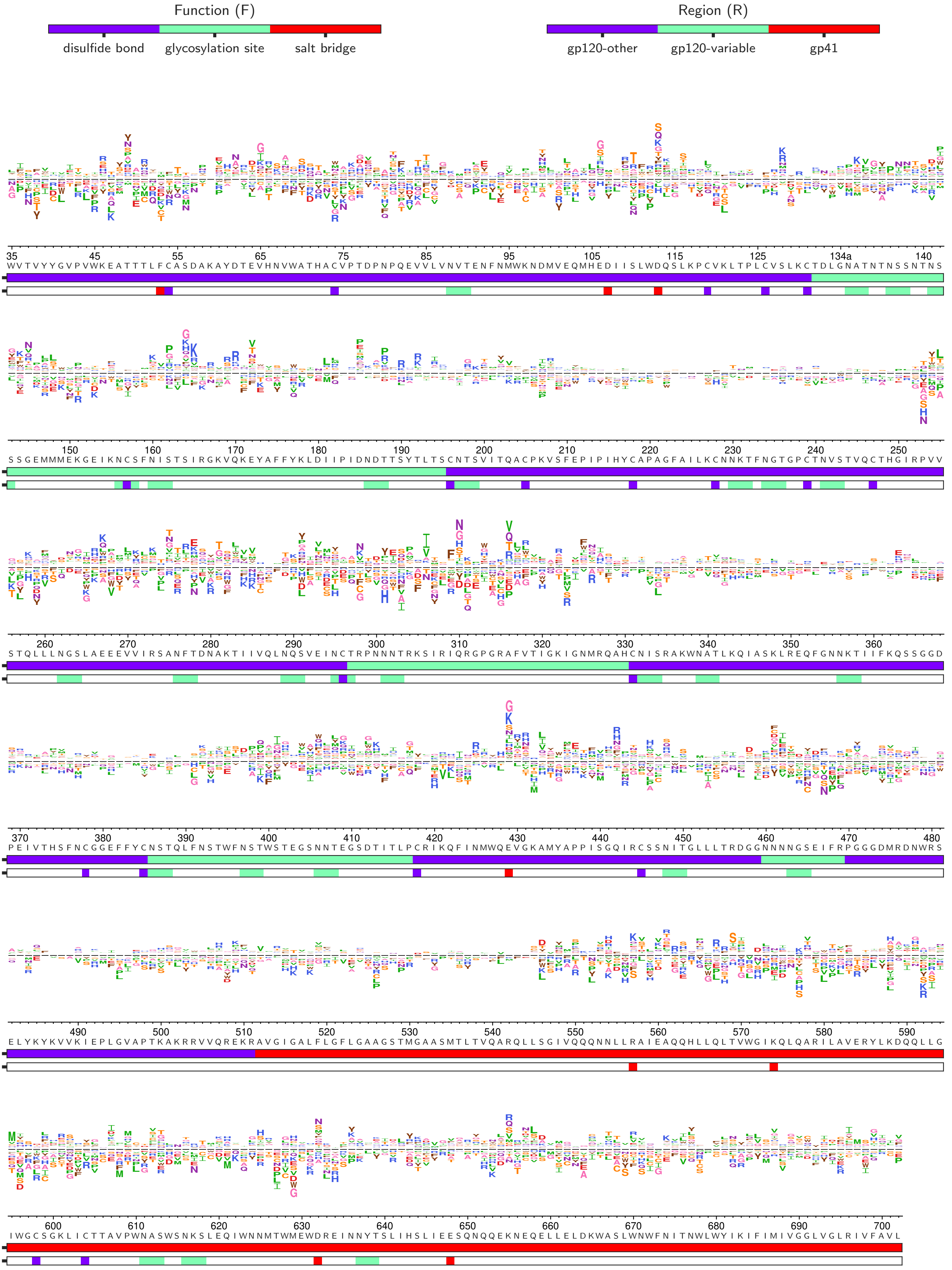

### XBP1sATF6_diffsel.pdf

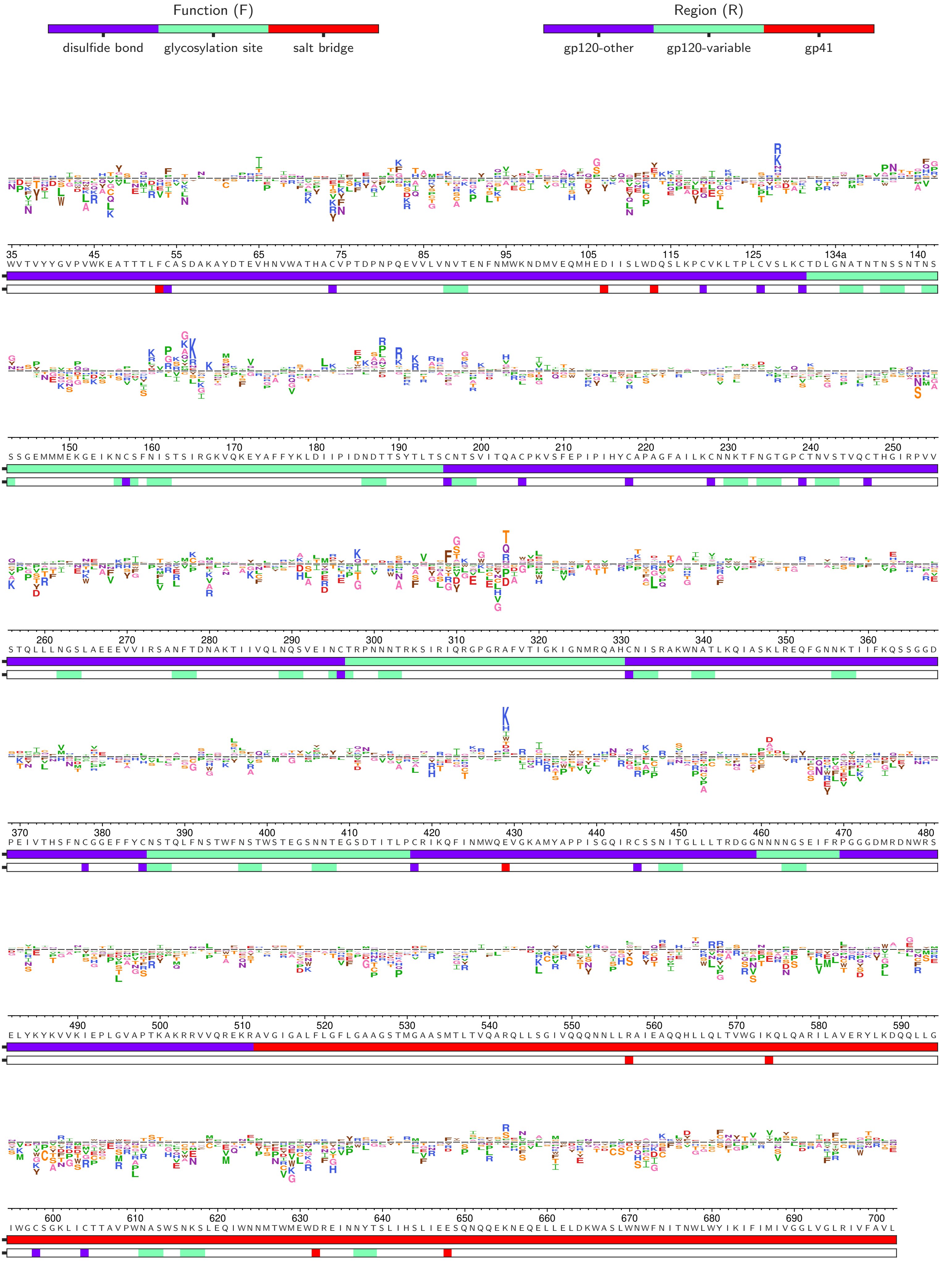
